# “Divergence and gene flow history at two large chromosomal inversions involved in long-snouted seahorse ecotype formation”

**DOI:** 10.1101/2023.07.04.547634

**Authors:** Laura Meyer, Pierre Barry, Florentine Riquet, Andrew Foote, Clio Der Sarkissian, Regina Cunha, Christine Arbiol, Frédérique Cerqueira, Erick Desmarais, Anaïs Bordes, Nicolas Bierne, Bruno Guinand, Pierre-Alexandre Gagnaire

## Abstract

Chromosomal inversions can play an important role in divergence and reproductive isolation by building and maintaining distinct allelic combinations between evolutionary lineages. Alternatively, they can take the form of balanced polymorphisms that segregate within populations over time until one arrangement becomes fixed. Many questions remain about how these different inversion polymorphisms arise, how the mechanisms responsible for their long-term maintenance interact, and ultimately how they contribute to speciation. The long-snouted seahorse (*Hippocampus guttulatus*) is known to be subdivided into partially isolated lineages and marine-lagoon ecotypes differentiated by structural variation. Here, we aim to characterise these differences along the entire genome, and to reconstruct their history and role in ecotype formation. We generated a near chromosome-level reference genome assembly and described genome-wide patterns of diversity and divergence through the analysis of 112 whole-genome sequences from Atlantic, Mediterranean, and Black Sea populations. Combined with linked-read sequencing data, we found evidence for two megabase-scale chromosomal inversions showing contrasted allele frequency patterns across the species range. We reveal that these inversions represent ancient intraspecific polymorphisms, one being likely maintained by divergent selection, and the other by associative overdominance. Haplotype combinations characterising Mediterranean ecotypes also suggest the existence of potential interactions between the two inversions, possibly driven by environment-dependent fitness effects. Lastly, we detected gene flux eroding divergence between inverted alleles at varying levels between the two inversions, with a likely impact on their long-term dynamics.

## Introduction

How genetic differences accumulate between nascent species and contribute to the buildup of reproductive isolation remains a fundamental question in evolutionary biology (Westram et al., 2022a). Mechanisms that bring distant sites into linkage disequilibrium and maintain strong allelic associations in the presence of gene flow are key factors favouring the emergence of reproductive isolation (Butlin, 2005; Ortiz-Barrientos et al., 2016; Tigano & Friesen, 2016). Structural variants (SVs) such as chromosomal inversions, which both establish and maintain haplotypes by reducing recombination, combine these two properties (Faria & Navarro, 2010; Hoffmann & Rieseberg, 2008; Mérot et al., 2020a; Wellenreuther et al., 2019; Zhang et al., 2021). For these reasons, SVs are often the focus of speciation genomics studies, particularly those interested in ecotype formation (e.g. Atlantic cod, Berg et al., 2016; rough periwinkles, Faria et al., 2019a; deer mice, Harringmeyer & Hoekstra, 2022; seaweed flies, Mérot et al., 2020b; sunflowers, Todesco et al., 2020; monkeyflowers, Lowry & Willis, 2010). Despite the growing number of studies implicating inversions in divergence between ecotypes in a variety of organisms, many questions remain about the conditions in which these inversions arise, their long-term maintenance, and their contribution to the strengthening of reproductive isolation (Faria et al., 2019b; Kirkpatrick, 2010; Wellenreuther & Bernatchez, 2018; Westram et al., 2022b).

The most immediate role of inversions is generally ascribed to recombination suppression, which protects combinations of locally advantageous or co-adapted alleles locked within an inversion (Butlin, 2005; Kirkpatrick, 2010; Noor et al., 2001; Rieseberg, 2001). Inversions that span large chromosome segments are likely to bring various types of mutations from remote functional sites into linkage disequilibrium, thus increasing the chance of creating and preserving haplotype combinations of large effect on the phenotype. Many studies have directly linked inversions to traits that differ between ecotypes, with functional implications in local adaptation or reproductive isolation (Campbell et al., 2021; Funk et al., 2021; Gould et al., 2017; Hager et al., 2022; Jay et al., 2022; Lundberg et al., 2023). Because these usually complex phenotypes are likely to involve several genes that are inherited as a single block, inversions are sometimes referred to as ‘supergenes’ (Lamichhaney et al., 2016; Matschiner et al., 2022; Schwander et al., 2014; Thompson & Jiggins, 2014). Besides these alleles directly providing local fitness advantage, a single inversion can be expected to carry other selected sites, such as recessive deleterious mutations that have been captured by chance (Berdan et al., 2021; Jay et al., 2021; Nei et al., 1967). The presence of many selected loci, along with the potential interactions between them, means that the dynamics of an inversion may be driven by multiple simultaneous processes (Faria et al., 2019b; Guerrero et al., 2012). What is more, the allelic contents of an inversion are not immutable and could change over time, for instance, by the accumulation of new mutations. Another mechanism that can modify the allelic contents of inverted haplotypes is the exchange of genetic material between alternate arrangements through gene flux (Cheng et al., 2012; Faria et al., 2019b; Matschiner et al., 2022; Navarro et al., 1997; Schaeffer & Anderson, 2005). This can occur through double crossover events with the formation of an inversion loop or through non-crossover gene conversion during the repair of double-stranded breaks (Korunes & Noor, 2019; Matschiner et al., 2022). Changes in allelic content could affect the processes governing inversion frequency (Berdan et al., 2021) and eventually impact its evolutionary trajectory and contribution to reproductive isolation.

Considering the lifetime evolution of inversions provides a particularly useful framework for studying their evolutionary dynamics and their potential role in speciation (Faria et al., 2019b). Disentangling the roles of different processes acting on an inversion at different stages of its evolution allows us to go further than simply taking inventory of the inversions present in a given system. For the sake of clarity, Faria et al. (2019b) distinguished two main classes of inversion polymorphisms, which they refer to as Type I and Type II. Briefly, Type I inversions are expected to show (near) local fixation of one haplotype and maintenance of between-population variation. This pattern may be caused by divergent selection acting on alternate haplotypes conferring local adaptation and/or a form of bistable selection involving frequency dependence, incompatibility selection or assortative mating. A Type I polymorphism may show signs of underdominance either due to direct effects (loss of unbalanced recombinant gametes during meiosis) or due to the allelic contents of the inversion (Faria et al., 2019b; Kirkpatrick & Barton, 2006). Over time, Type I inversions likely contribute to the accumulation of Dobzhansky-Müller (DM) incompatibilities and reinforcement mechanisms via coupling with other reproductive isolation polymorphisms (Kulmuni et al., 2020; Navarro & Barton, 2003). However, inversions are not inevitably subject to divergent selection but can alternatively be maintained by a form of balancing selection (Jay et al., 2021; Mérot, Llaurens, et al., 2020; Wellenreuther & Bernatchez, 2018b; Yeaman, 2013; Yeaman & Whitlock, 2011). Faria et al. (2019b) recognise these Type II inversions as polymorphisms that are maintained by mechanisms such as frequency-dependence, disassortative mating, or antagonistic pleiotropy. Type II inversions might also contain overdominant loci or show heterokaryote advantage through associative overdominance (Marion & Noor, 2023). Indeed, some inversions can be expected to suffer from high mutation load, due to deleterious mutations hitchhiking with selected genes, or simply accumulating in low recombination regions (Hill & Robertson, 1966; Nei et al., 1967). It is important to note that these various processes associated with Type I and Type II polymorphisms are not necessarily mutually exclusive and might act in concert to shape the evolutionary dynamics of a given system.

The long-snouted seahorse (*Hippocampus guttulatus*) is one of two seahorse species occurring along European coastlines. A previous study that addressed population structure in this species found evidence for the presence of semi-isolated lineages with sharp spatial boundaries (Riquet et al., 2019). Using a set of SNP markers, this study showed the presence of a northern and a southern lineage in the Atlantic, while Mediterranean populations were structured according to habitat type. In the Mediterranean Sea, the authors observed ecological structure and the presence of marine and lagoon ecotypes. While analyses in this study were not based on a reference genome for the species, indirect evidence indicated that genetic differentiation between seahorse lineages was concentrated in a large genomic island of divergence that suggested the implication of a structural variant such as a chromosomal inversion. We here introduce the first genome-scale study in *H. guttulatus*, aiming to characterise and understand the origin and evolution of genome-wide divergence patterns among lineages. We generate a near chromosome-level assembly of the long-snouted seahorse genome, and describe genetic variation across habitats throughout the range distribution of the species. We found the presence of not one, but two megabase-scale inversions, which may have played an important role in ecotype formation in the Mediterranean Sea. We then attempt to reconstruct the life history of these two inversions, especially focusing on their origin, divergence and subsequent evolution, including gene exchange between the inverted alleles. Our study highlights interactions between two inversions in the same system and the impact of gene flux between arrangements.

## Materials and methods

### Sampling and DNA extraction

Samples were collected across the species range and from different habitats (Supplementary Table S1). Nonlethal fin or tail clips were taken from fresh fish tissue samples and preserved in 95% ethanol. DNA was extracted using the Nucleospin Tissue kit ® (Macherey–Nagel, Germany). Dried seahorse samples were obtained from private collections through a call in a local newspaper. The approximate date (1935-2010) and collection site were recorded based on information provided by the donors. The dorsal fin of each dried seahorse was scratched to collect c.a. 20 μg of tissue powder. Genomic DNA was extracted using a standard CetylTrimethyl Ammonium Bromide (CTAB) Chloroform:Isoamyl alcohol (24:1) protocol (Doyle & Doyle, 1987). Lastly, four alcohol-preserved museum samples (1856-1898) were provided by the National Museum of Natural History (Paris, France). Tissue fragments (1-2 mm) were obtained by internal needlepoint sampling (Haÿ et al., 2020) and subjected to overnight enzymatic digestion (40 µL proteinase K at 25 mg/mL for 500 µL volume, at 56 °C). DNA was extracted using a phenol-chloroform method (Campos & Gilbert, 2012) in a dedicated clean lab facility located at the Institute of Evolutionary Science of Montpellier (ISEM, France).

### Assembly and repeat annotation of the *H. guttulatus* reference genome

We performed high-coverage linked-read sequencing of an Atlantic long-snouted seahorse from the Hossegor lagoon (Bay of Biscay) to generate a high-quality reference genome assembly (hereafter referred to as Hgutt_V1). Fresh gill and muscle tissue were solubilized in a 25 ml solution of TNES-Urea (10 mM Tris-HCl, 120 mM NaCl, 10 mM EDTA, 0.5% SDS, 4 M urea, PH 8) during 4 weeks at 20°C. High molecular weight genomic DNA (HMW gDNA) was isolated using three phenol-chloroform followed by two chloroform extractions after digestion with proteinase K (150 µg/ml). Final precipitation was performed using two volumes of 100% Ethanol. The resulting pellet was washed several times in 80% ethanol and resuspended in ultra pure water by heating to 65°C, before being kept at 40°C during 4 days. The length distribution of extracted DNA molecules was assessed by electrophoresis on a TapeStation Genomic DNA ScreenTape assay (Agilent Technologies). Single-stranded DNA damage was treated with the NEBNext FFPE DNA Repair mix and repaired DNA was then subjected to size selection to remove fragments shorter than 40 kb using a PippinHT instrument (Sage Science) with a 0.75% Agarose Gel Cassette. HMW gDNA was submitted to the 10x Genomics linked-read library preparation following the Chromium Genome Reagent Kit v2 protocol at the MGX sequencing facility (CNRS, Montpellier, France). The genome library was sequenced to ∼100X on a S1 lane of an Illumina NovaSeq6000 in 150 bp paired-end mode by Genewiz Inc (USA), generating ∼0.3 billion reads.

Raw demultiplexed reads were deduplicated using *nubeam-dedup* (Dai and Guan 2020) and processed with *process_10xReads.py* (https://github.com/ucdavis-bioinformatics/proc10xG) to extract the barcode sequence of each read pair and the number of read pairs associated to each barcode. The distribution of the number of read pairs per barcode was then analysed to identify rare barcodes potentially generated by sequencing errors and over-represented barcodes (Supplementary Figure S1). A total of 143.7 million paired-end reads carrying 1.455 million retained barcodes were finally extracted using *proc10xG* scripts and used for linked-read-based *de novo* genome assembly using the *Supernova-2.1.1* software package (Weisenfeld et al., 2017). Assembled scaffolds were outputted in pseudohap style, with a minimum size set to 1 kb. Assembly statistics of the Hgutt_V1 reference genome were computed and visualised with the *BlobToolKit v3.5.2* software suite (Challis et al., 2020), combined with an assessment of genome assembly completeness with *BUSCO 5.4.4* (Manni et al., 2021) using the actinopterygii_odb10 fish dataset containing 3640 conserved genes. Whole-genome alignment was performed with *Minimap2* (Li, 2018) and visualised using *D-GENIES* (Cabanettes & Klopp, 2018)(Cabanettes and Klopp 2018) to compare and anchor *H. guttulatus* scaffolds to the chromosome-scale assembly of the closely related *H. erectus* (Li et al., 2021), provided by the authors. We also used the genome sequence of a north Atlantic long-snouted seahorse (UK), which was assembled by Iridian Genomes by ordering and orienting pre-assembled contigs based on other fish reference genomes (Accession PRJNA481552, hereafter called HguttRefA).

We used *RepeatModeler2* (Flynn et al., 2020) for *de novo* repeat finding and identification of the unique transposable element families present in the seahorse genome. In addition, we searched for tandem repeats (TRs) following the strategy developed in Melters et al. (2013), using the same parameter values to run *Tandem Repeats Finder* v4.09.1 (Benson, 1999) within the *pyTanFinder* pipeline (Kirov et al., 2018). Finally, *RepeatMasker* v4.0.5 (http://repeatmasker.org) was used to perform repeat annotation and masking of the identified repeat elements.

### Library preparation and whole-genome resequencing

Whole-genome sequencing (WGS) libraries were prepared for 90 samples using the Ovation Ultralow System V2 library preparation kit (NuGEN/Tecan) from 100 ng DNA input (when possible) following the manufacturer’s instructions. We used unique dual indexing to minimise the effect of index-hopping and PCR cycles were adapted to the amount of input DNA (9-20 cycles). Libraries were pooled in equimolar ratio and sequenced to different coverage depths on a single S4 flow cell on a NovaSeq6000 instrument (Illumina) to generate 150 bp paired-end reads.

### Sequence processing and alignment

To complement our dataset, WGS data for an additional 26 seahorse samples were obtained from Barry et al. (2022) (TruSeq DNA PCR-free libraries sequenced on Illumina NovaSeq6000, GenBank Sequence Read Archive, accession BioProject ID PRJNA777424). Therefore, our final combined dataset consisted of 112 samples (dataset #1, Fig. 1A, Supplementary Table S1). Raw demultiplexed reads were processed using fastp (v0.23.1) (Chen et al., 2018) with the “ *--merge*” option, in order to stitch together paired-end reads with overlapping sections. Both merged and unmerged reads were aligned to our reference genome using BWA-MEM (BWA v0.7.17; (H. Li, 2013). Picard (v2.26.8) (« Picard toolkit », 2019) was used for sorting read alignments, marking duplicates and adding read groups. DNA damage patterns in older samples were visualised using PMDtools (v0.60) (Skoglund et al., 2014) (Supplementary Figure S2).

**Fig. 1.**
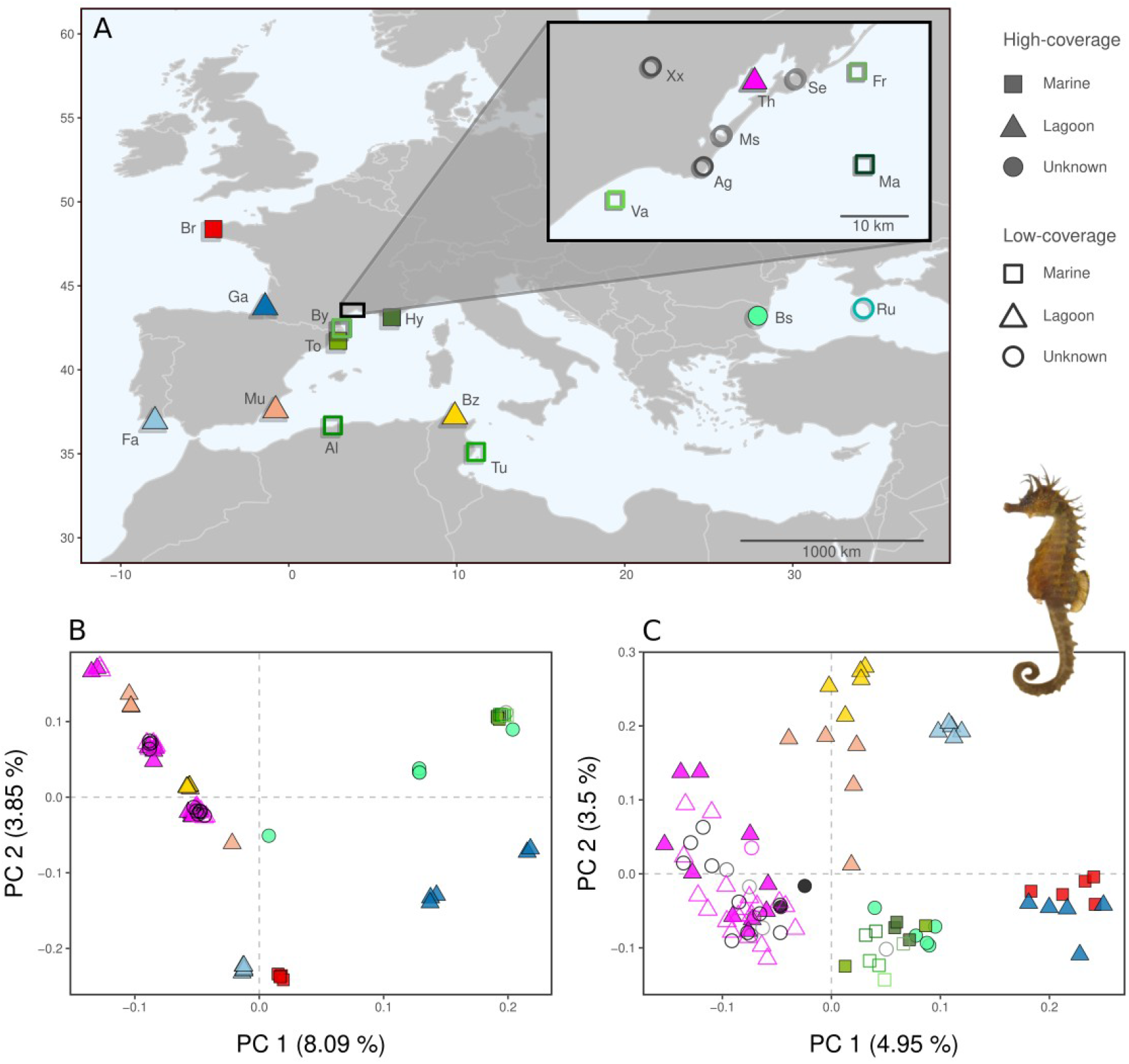
A) Sampling map for all long-snouted seahorse samples (n=112, dataset #1). Triangles: lagoon sites. Squares: marine sites. Circles: unknown habitat type. Filled shapes: high-coverage samples for which variants were called (dataset #2). Unfilled shapes: samples for which variants were not called. Br: Brest. Ga: Port d’Albret. Fa: Faro. Mu: Murcia. To: Tossa de mar. By: Banyuls. Va: Valras. Ag: Agde. Ms: Marseillan. Th: Thau lagoon. Se: Sète. Fr: Frontignan. Hy: Hyères. Ma: unknown Mediterranean marine site. Al: unknown Algerian site. Bz: Bizerte lagoon. Tu: unknown Tunisian site. Bs: Varna. Ru: unknown Russian site. Xx: unknown location. B & C) PCA of 89 individuals with sufficient coverage (>1X) based on IBS distances calculated in ANGSD. PCA was performed using all genome-wide markers (1,478,955 SNPs) (B) and only using a subset of markers at linkage equilibrium (898 SNPs) (C). © Seahorse picture Iglésias 2013.

### Variant calling of medium-to high-coverage samples

Forty-eight samples with sufficient coverage (∼5-40X) were selected for variant calling in such a way that this subset contained five individuals per location (dataset #2, Fig. 1A). Variants were called using the GATK best practices workflow (McKenna et al., 2010; Van der Auwera et al., 2013). Firstly, individual GVCF files were created from bam files with HaplotypeCaller (GATK v.4.1.8.0). This information was then stored in a GVCF database using GenomicsDBImport, and VCF files (one file per scaffold) were generated with GenotypeGVCFs. After concatenation, the resulting VCF was filtered for indels, multiallelic SNPs and missing data (“*--max-missing 0.9*”). Our detailed workflow and commands used are provided in the data availability section.

### Population structure

A Principal component analysis (PCA) was carried out on samples with sufficient coverage (>1X) from dataset #1 without calling genotypes. Genotype likelihoods were calculated using ANGSD (v0.933) (Korneliussen et al., 2014) with the GATK model (“*-GL 2*”) and the following parameters: “*-doMajorMinor 1-minMapQ 30-minQ 20-doMaf 1-doCounts 1-minMaf 0.05-uniqueonly 1-remove_bads 1-C 50-baq 1-doCov 1-doIBS 2-makeMatrix 1-ref referencegenome_Hgutt_V1.fa*”. In addition, the first 5 bp were trimmed off of reads (-*trim 5*) to account for DNA damage patterns due to cytosine deamination in historical samples (Supplementary Figure S2). The analysis was restricted to sites with a strong probability of being SNPs (“*-SNP_pval 1e-6*”). Finally, sites were not considered if all samples had coverage higher than 30X (“*-setMaxDepth*”), in order to avoid artefacts in problematic regions that receive unexpectedly high coverage (low-complexity and duplicated regions), or if less than half of the individuals had data (“*-minInd*”). The same filters were used in subsequent analyses conducted with ANGSD, unless specified otherwise. To perform a PCA while minimising the effect of linkage disequilibrium, we used ANGSD with the “*-sites*” option and provided a file containing one randomly selected SNP per 10 kb window in every 500 kb interval. To complement this genotype likelihood-based approach, PCA was also performed on called genotypes from dataset #2 using the R package SNPRelate (v1.28.0) (Zheng et al., 2012). Lastly, we used local PCA to capture variation in population structure along the genome in order to identify outlying patterns indicating the presence of putative chromosomal inversions. To this end, local PCA was performed in non-overlapping windows of 5 kb using the R package lostruct (v0.0.0.9, H. Li & Ralph, 2019).

### Analysis of large structural variants

Genomic regions that were candidates for the presence of large SVs were more specifically tested for evidence of chromosomal inversions in linked-read sequencing data. Firstly, we used abrupt signal shifts flanking outlying regions in the local PCA to determine the location of putative inversion breakpoints. We performed manual curation of the reference genome assembly in those regions following (Rhie et al., 2021), using the alignment to the genome assembly of *H. erectus* to confirm the suspected breakpoints. We then used *MTG-Link* v2.4.1 (Guichard et al., 2022) to perform local re-assembly of the linked-reads mapping in the regions surrounding breakpoints. The 10X linked-reads data were preprocessed with *EMA* v0.6.2 (Shajii et al., 2018) and *LRez* v2.2.4 (Morisse, Lemaitre, et al., 2021) before running *MTG-Link* with default settings. The locally reassembled contigs were finally aligned against the Hgutt_V1 reference genome with *Minimap2* (H. Li, 2018) to check for local consistency between assemblies and identify possible mis-joins. We characterised repeat content around the inversion breakpoints, looking specifically for the presence of recombinogenic sequences such as long inverted repeats (LIRs). Finally, we used *Leviathan* v1.0.2 (Morisse, Legeai, et al., 2021) for calling structural variants using linked-read data information connecting distant regions that share a higher number of barcodes than expected based on their distance. We ran *Leviathan* on the 10X linked-read data from the Hgutt_V1 assembly mapped to the HguttRefA genome.

Moreover, we followed the haplotagging library construction protocol described in (Meier et al., 2021) to generate complementary linked-read sequencing data for alternate genotypes at the two inversions. Namely, we constructed haplotagging libraries for a DD/AA Mediterranean (Hgutt_MU_09) and a CC/AB Atlantic individual (Hgutt_GA_13), and sequenced them with 2*50 bp paired-end reads to an average coverage depth of 12X.

### Genomic landscape of divergence and introgression

We characterised the genomic landscape of divergence between localities and habitats using dataset #2. Genetic differentiation (*F_ST_*), nucleotide diversity (*π*) and absolute genetic divergence (*D_XY_*) was calculated in 25 kb windows (with “*-m 100”*) using the popgenWindows.py script (Martin, 2018; https://github.com/simonhmartin/genomics_general). We sought to characterise two highly divergent regions on Chr2 and Chr12, which represented suspected inversions. We used BAMscorer v1.4 (Ferrari et al., 2022) to assign inversion genotypes in all samples (including low-coverage samples). This firstly consisted of classifying all samples in dataset #2 with regards to their haplotypes, as based on PCA groupings and individual heterozygosity values calculated with VCFtools (v0.1.16) (Danecek et al., 2011). This reference database was then used to score alignments and to ascertain allelic state and haplotype for all samples.

In order to test for introgression with other seahorse species and to determine evolutionary relationships across the genome, we obtained whole-genome resequencing data for *H. erectus*, *H. hippocampus*, *H. zosterae*, *H. capensis* and *H. comes* from Li et al. (2021) (NCBI BioProjects accession code PRJNA612146), including data for one individual from each species. Reads were processed and aligned to the Hgutt_V1 reference genome as described above. ANGSD was used to call genotypes and to produce a VCF (“*-doBcf*”) containing seven individuals (two *H. guttulatus* that were homozygous for the four alternate inversion haplotypes, i.e. AA-DD and BB-CC, and five individuals from other species). This VCF was filtered to include only fixed sites (no heterozygous genotypes) and was used as input for performing topology weighting using the Twisst pipeline with an iterative sampling of subtrees (Martin & Van Belleghem, 2017). We generated unrooted phylogenies using PhyML (Guindon et al., 2010) as implemented in the script *phyml_sliding_windows.py* (https://github.com/simonhmartin/twisst) for windows of 50 SNPs, allowing a minimum of 25 non-missing genotypes per individual per window (“*--minPerInd 25”*). We produced a genome-wide consensus tree, as well as consensus trees for the two inversions using the averageTree function from phytools (Revell, 2012). Finally, topology weighting was performed using the *twisst.py* script.

### Inferring inversion history from the Ancestral Recombination Graph

To describe the joint genealogies of sampled DNA sequences along the genome, we sought to infer the Ancestral Recombination Graph (ARG) using tsinfer (v0.3.0) (Kelleher et al., 2019). This approach requires an input VCF file that describes a set of phased, diploid genomes, and the ancestral state of each mutation present in the INFO field. We carried out statistical phasing and missing data imputation of per-chromosome VCF files using SHAPEIT (v4.2.2) (Delaneau et al., 2019), assuming constant recombination rate of 1 cM per Mb and an effective population size (*N_e_*) estimated based on genome-wide diversity values (Charlesworth, 2009). To infer ancestral states, we used BLAST searches to determine the fixed allelic state of each SNP position in each of three outgroup species (*H. erectus, H. kuda* [Accession GCA_901007745.1]*, H. comes* [Accession GCA_001891065.2]) using the BLAST+ package (v2.2.28) (Camacho et al., 2009). Flanking regions of 100 bp on either side of each SNP were blasted to the reference genome of each outgroup using *blastn*. The top hit was inspected to determine the state of the homologous position in each outgroup using a custom script. Thereafter, est-sfs (v2.04) (Keightley & Jackson, 2018) was run using the Kimura 2-parameter model to infer ancestral state probabilities and phased VCFs were annotated with the most likely ancestral variant (bcftools v1.9; Danecek et al., 2021).

We reconstructed the ARG by running tsinfer on our phased, oriented VCF containing all chromosomes. Sample objects were created using the CYVCF2 library (v0.30.18; Pedersen & Quinlan, 2017) as in the tskit tutorial (https://tskit.dev/tsinfer/docs/stable/tutorial.html). The tree sequence was then inferred using a recombination rate of 1e-8 and a mismatch ratio of 1. The inferred trees were dated using tsdate (v0.1.5) with “*N_e_=200000*”, “*timepoints=100*” and “*mutation_rate=1e-8*”. We selected a large estimate for *N_e_* based on the diversity values of the most divergent region of the genome (i.e. the inversion on Chr12). Opting for a large *N_e_* value allowed tsdate to assign old ages to deep tree nodes to take into account ancient coalescent times that largely predate within-population coalescent times. To study divergence times along the genome, the time to the most recent common ancestor (TMRCA) for random combinations of haploid samples was extracted from the tree sequence. To characterise local ancestry in suspected inversion regions, we considered the two branches located directly below the oldest node of each tree, and identified the samples below each of these two branches. To assign ancestry to each branch (inverted or non-inverted haplotype), a branch was considered to present a given haplotype if it satisfied one of two conditions: (i) the branch contained more than 75% of the haplotype copies from all of the homokaryotes of that haplotype, and the other branch contained more than 75% of the copies from the opposite haplotype; (ii) the branch contained 100% of the copies from all of the homokaryotes of that haplotype, and the same branch contained less than 75% of the copies from the opposite haplotype.

## Results

### Reference genome

We obtained a near chromosome-level assembly of the *Hippocampus guttulatus* Hgutt_V1 reference genome. The assembly contained 3878 scaffolds spanning 451 Mb (424 Mb in scaffolds of at least 10kb, longest scaffold 28.8 Mb, scaffold N50 = 18.1 Mb, N90 = 5.65 Mbp, Supplementary Figure S3), which is close to the genome size of 424 Mb predicted by GenomeScope (Barry et al., 2020). Genome completeness was very high (Busco C:95.6% [S:94.4%, D:1.2%], F:1.6%, M:2.8%, n:3640) and comparable to the *H. erectus* and the HguttRefA assemblies (Supplementary Figure S4). All scaffolds showed strong homology and conserved synteny relationships with the chromosome-level genome assembly of *H. erectus* (Supplementary Figure S5), suggesting that Hgutt_V1 is a high-quality assembly. Scaffolds anchoring to the *H. erectus* chromosome assembly produced 22 pseudomolecules that were used to characterise the genomic landscape of divergence (Fig. 2).

**Fig. 2.**
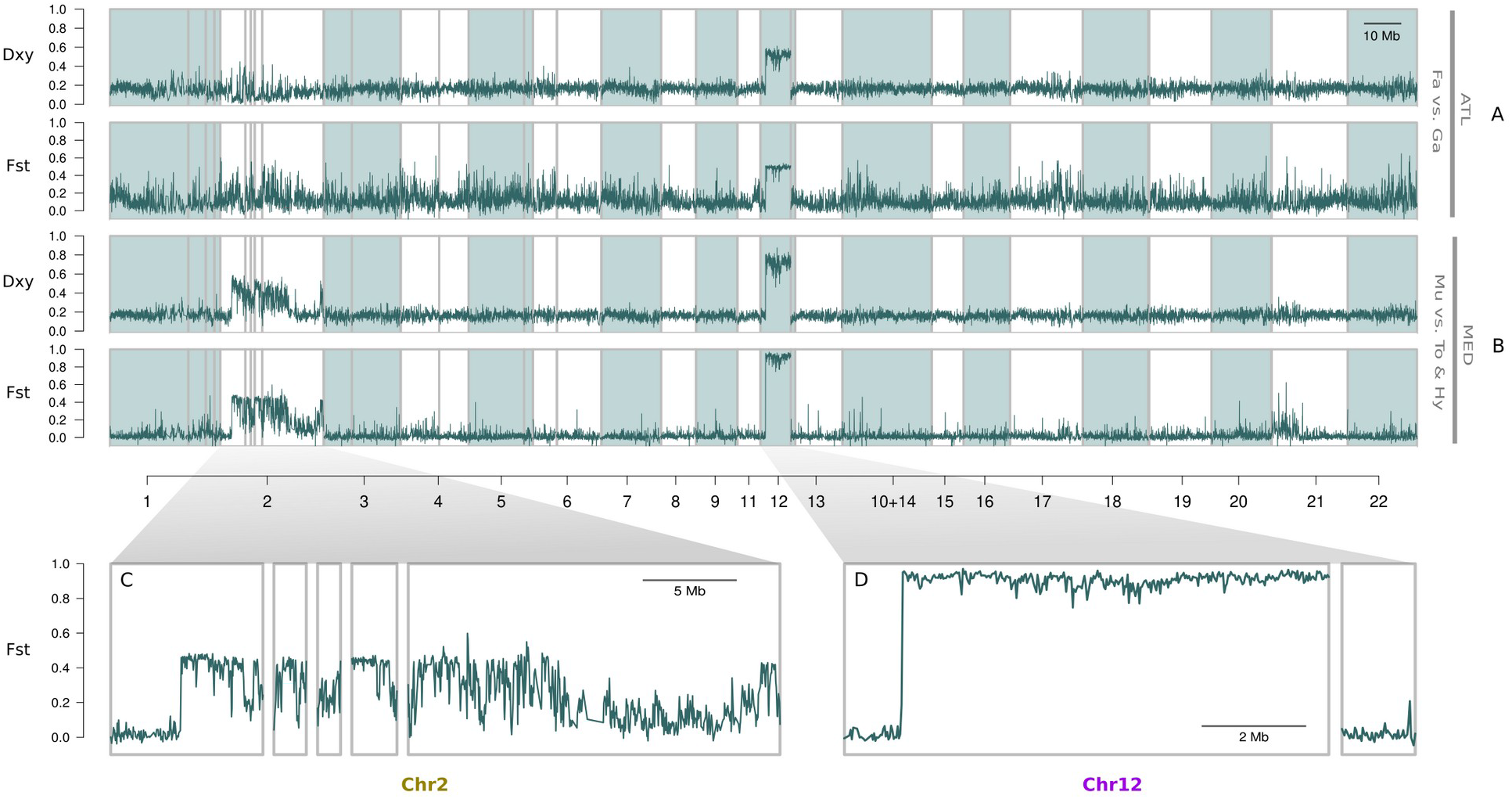
Genomic landscape of absolute divergence (*D_XY_*) and differentiation (Fst) calculated in 25 kb sliding windows between a North Atlantic and a South Atlantic population (A) and between a Mediterranean lagoon and a Mediterranean marine population (B) (5 high-coverage samples per population). *H. guttulatus* scaffolds (grey rectangles) were aligned to the chromosome-level *H. erectus* assembly and are displayed according to their homology with the 22 *H. erectus* chromosomes, alternating white and blue rectangles. Regions of highest divergence clustered on two chromosomes: Chr2 (∼32 Mb) (C) and Chr12 (∼11 Mb) (D), which both carry large chromosomal inversions. Exact ordering and orientation of *H. guttulatus* scaffolds within chromosomes 2 and 12 could not be determined due to rearrangements between species (Supplementary Figure S5).

The repeat landscape showed that interspersed repeats add up to 33% of the genome, with 13% being occupied by unknown repeats (Supplementary Figure S6). Among these unknown repeats, the clustering analysis of similar tandem repeats revealed a highly abundant TR of approximately 500 bp in length, present in more than 5500 copies across the genome, with an accumulated abundance of 2.7 Mb. The consensus monomer of this TR, hereafter called Hgutt_Tan9, is a short sequence of 37bp present in all chromosomes (Supplementary Figure S7).

### Overall genetic structure

We produced genome resequencing data of heterogeneous quality (ranging from <1X to >40X) and therefore characterised overall genetic structure using a genotype likelihood-based approach. We found evidence for pronounced population structure, even between certain sampling locations that were in close geographic proximity (Fig. 1B). This genetic structure could largely be ascribed to markers that are in linkage disequilibrium (LD), since PCA performed on unlinked markers revealed a different pattern (Fig. 1C). In the latter analysis, we observe that Mediterranean marine samples grouped together, separate from the different Mediterranean lagoon populations. We discern a northern and a southern cluster of samples in the Atlantic. By contrast, if all markers are considered (Fig. 1B), samples do not group together based on geographical origin. Instead, we observe three groups spread out along Principal Component (PC) axis 1, as well as three groups along a different axis. Observing three PCA clusters is consistent with the presence of large chromosomal inversions, with the three groups representing different inversion genotypes - homokaryotes with two inverted alleles, heterokaryotes carrying both haplotypes, and non-inverted homokaryotes. The observed separation along PCA axes is thus reminiscent of many segregating sites captured in one or more polymorphic inversions.

We observed heterogeneous landscapes of differentiation along the genome (Fig. 2). In general, genome-wide *F_ST_* values ranged between 0.06 and 0.15 (first and third quartiles) for the comparison between northern and southern Atlantic lineages (Fig. 2A, Supplementary Figure S8). Background genomic differentiation was weaker between Mediterranean marine and lagoon ecotypes, with *F_ST_* values ranging between 0 and 0.03 (Fig. 2B, Supplementary Figure S8). Other contrasts between different populations in the Atlantic and between populations in the Mediterranean yielded similar diversity and divergence landscapes (Supplementary Figures S8, S9 & S10). These relatively homogenous, weak differentiation patterns contrasted with high *F_ST_* and *D_XY_* values found on two chromosomes - Chr2 (Fig. 2C) and Chr12 (Fig. 2D). Chr12 presented an 8.2 Mb long plateau of high *F_ST_* and *D_XY_* values, as has previously been associated with inversions segregating at different frequencies. For Chr2, the pattern was less clear, as the highest values did not occur in one contiguous block, but were split across scaffolds and even showed discontinuities within scaffolds.

### Inversions differentiate ecotypes in the Mediterranean Sea

We identified two highly differentiated blocks that putatively represent chromosomal inversions differentiating seahorse lineages (Fig. 2). The presence of an inversion for the block on Chr12 was first supported by the alignment of Hgutt_V1 against HguttRefA (generated from the opposite homokaryote, (Supplementary Figure S11), as well as with the *H. erectus* assembly. This analysis showed the presence of an 8.2 Mb long inverted segment between the two *H. guttulatus* assemblies, with a putative inversion breakpoint located near position 1.699Mb on scaffold 14 of Hgutt_V1, which corresponds to the abrupt signal shift in divergence at the beginning of the block. Linked-read based local reassembly of a 3 kb region centred on the putative breakpoint confirmed the contiguity of the Hgutt_V1 reference. In the middle of this 3 kb region, we found a long inverted repeat (LIR) consisting of two inverted arrays of the Tan9 monomer, with the internal spacer overlapping the inversion breakpoint (Supplementary Figure S12). The other end of the inversion at the end of scaffold 14 (near 9.877Mb) also contained a tandem repeat of Tan9 monomers (Supplementary Figure S6). Linked-read sequencing data obtained from three individuals showing the three possible genotypes (AA, AB and BB) showed mapping patterns that were consistent with the presence of a large chromosomal inversion (Supplementary Figure S12). Finally, the analysis of linked-reads from a BB genotype mapped to the HguttRefA assembly allowed the direct detection of a 8.2 Mb inversion at the breakpoints expected from all previous analyses.

Although we could not make the same comparison for Chr2, since the two *H. guttulatus* assemblies both carry the same haplotype, other patterns suggested that a second inversion is highly likely. Performing PCA separately on Chr2 and Chr12, we observed three clusters along PC1 with the middle group presenting higher heterozygosity than the outer groups (Fig. 3A & 3B). Using BAMscorer (Ferrari et al., 2022), we characterised inversion genotypes in all samples except for one extremely low-quality individual. These scores were consistent with the groupings observed from the PCA (Fig. 1B, Fig. 3A & 3B) - that is, samples that were classified as heterokaryotes were in the middle PCA group, and homokaryote samples were in the outer groups. These patterns confirm that the multiple blocks of high *F_ST_* on Chr2 are in perfect LD, and that they collectively segregate as a single contiguous variant, despite the discontinuous signature observed in the divergence landscape (Fig. 2C). Consequently, our results indicate that there are two polymorphic inversions segregating in seahorse populations, and that these play an important role in the differentiation between lineages. Classifying inversion genotypes in all samples allowed us to study the distribution and relative frequency of alternate haplotypes in different sampling locations (Fig. 3C & 3F).

**Fig. 3.**
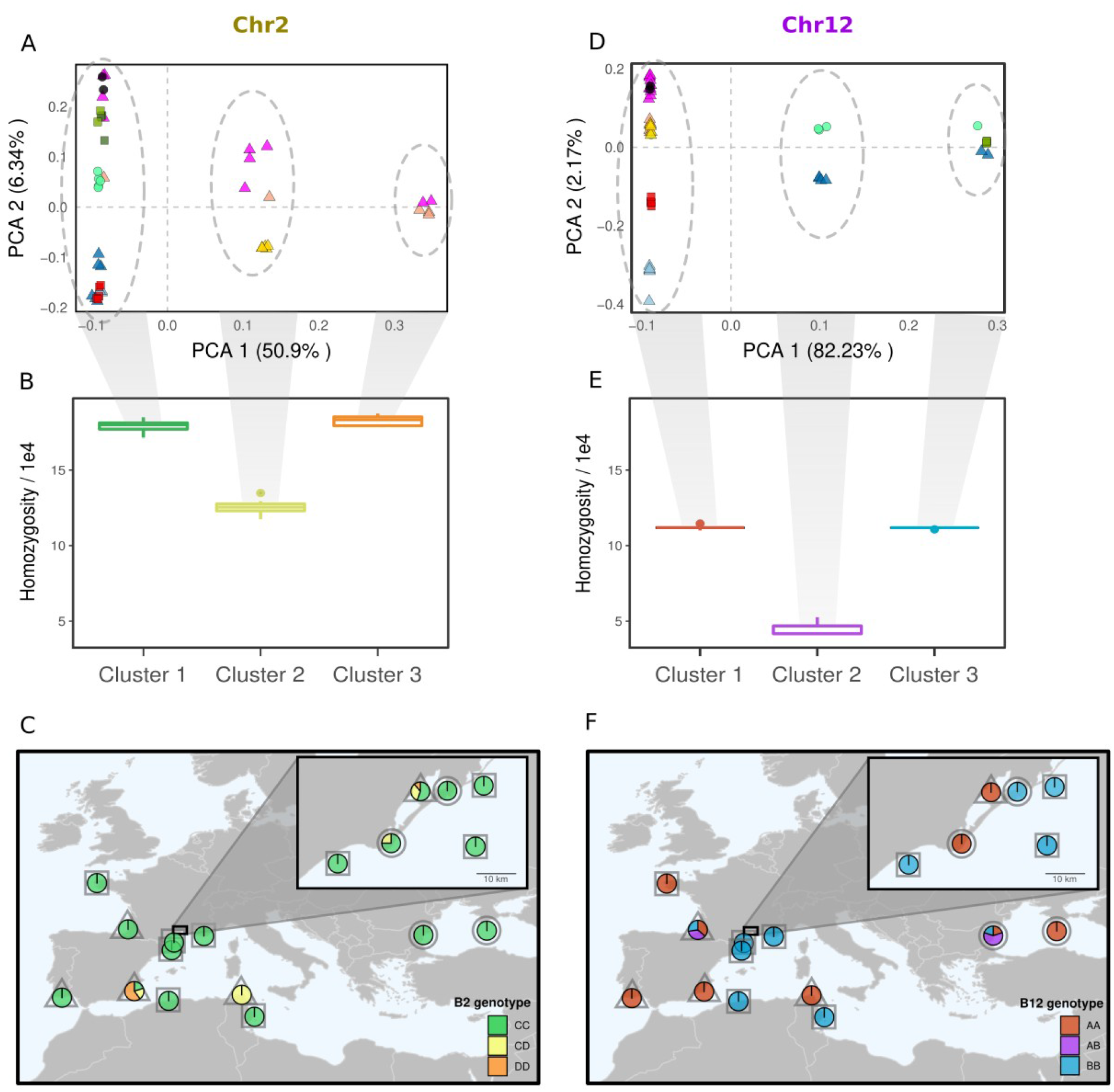
Molecular diversity patterns observed for Chr2 (A-C) and Chr12 (D-F), which carry large inversions (B2 and B12). A & D) Chromosome-wide Principal Component Analysis (PCA) of 48 high-coverage samples (dataset #2) using the same symbols as in Fig. 1. The three clusters for Chr2 (A) and Chr12 (D) are illustrated by grey ellipses. B & E) Boxplots of observed homozygosity for each cluster. Homozygosity was calculated at a chromosome-wide scale and averaged per sample. C & F) Maps of inversion genotype frequencies. Pie charts represent inversion genotypes and grey symbols indicate marine (square), lagoon (triangle) or uncharacterised (circle) habitats. Samples that could not be genotyped visually by PCA were genotyped using BAMscorer.

We found that the inversion on Chr12 (B12) was present in the homozygous state (either AA or BB) in almost all locations (Fig. 3F). Atlantic samples from the northern part of Biscay (Brest), the Gulf of Cadiz (Faro) and samples from Mediterranean lagoons (Thau, Murcia and Bizerte) exclusively presented homokaryotes for the A allele, while Mediterranean marine sites only presented homokaryotes for the B allele. Small-scale variation in inversion genotypes was especially pronounced in the Mediterranean Sea, since samples that were collected only a few kilometres apart showed differential fixation for B12 between marine and lagoon habitats (insert map, Fig. 3F). Samples for which precise location information was not available (grey circles) were either AA or BB homokaryotes, with A alleles probably hailing from lagoon habitats in these locations and B alleles from the sea. Samples of unknown origin (“Xx”) mostly carried the lagoon haplotype, probably reflecting large local lagoon populations and their easy access for artisanal and non-professional fishing activities. There were only two sampling locations which were polymorphic for B12: Port d’Albret (Bay of Biscay) and Varna (Black Sea), which presented samples from each of the three inversion genotypes (AA, AB and BB). The inversion located on Chr2 (B2) showed different distribution patterns compared to that of B12, since this polymorphism was only found in Mediterranean lagoons (Fig. 3C). Both in the Atlantic and in the Black Sea, all samples were fixed for one haplotype (C). The alternative haplotype (D) was found at varying frequencies (28% to 70%) in Thau, Bizerte and Murcia lagoons. All samples caught in the Bizerte lagoon were CD heterokaryotes (n=5), while 3 out of 5 samples from Murcia were DD homokaryotes. All samples carrying one or two D alleles had been classified as an AA homokaryote for B12 (particularly for samples from unknown habitats), indicating that the Chr2 polymorphism is private to Mediterranean lagoon lineages.

Our results thus indicate that two chromosomal inversions largely drive ecotype differentiation in the Mediterranean Sea. Lagoon populations are fixed for one haplotype of B12 (A), whereas populations in the sea only carry the alternate haplotype (B). In addition, there is a second inversion on Chr2 (with haplotypes C or D) differentiating marine and lagoon ecotypes. This inversion presents a polymorphism which is private to the lagoon ecotype. Even though one of these inversion polymorphisms (B12) is present in the Atlantic, it does not show differential fixation according to habitat type here. Instead, Atlantic populations to the north of Hossegor carry the lagoon allele (A), while both A and B alleles are present in the South Atlantic (Riquet et al., 2019). Thus, lineage differentiation between geographic lineages in the Atlantic relies on genome-wide differences rather than loci captured in inversions only. In what follows, we sought to further characterise the inversions and determine their respective histories, including their origins and ages.

### Evolutionary history of the two inversions

When investigating the origin of the inversions segregating in seahorses, we wished to test whether the heterogeneous divergence landscape resulted from the differential erosion of previously divergent genomes or alternatively from the emergence of newly divergent regions. To address this question, we studied coalescence times along the genome as inferred by tsdate (v0.1.5). Outside the inversion regions, we did not find a strong signature of ancient coalescence in the form of deep time to the most recent common ancestor (TMRCA) (Fig. 4G). Only chromosomes carrying inversions showed peaks of older TMRCA (up to 5X higher than in the genome background) and high *D_XY_* values (Fig. 2). What is more, except for the inversions, regions of high *F_ST_* were not shared between Atlantic and Mediterranean contrasts, suggesting a lack of parallelism in the genome background (Supplementary Figure S13).

**Fig. 4.**
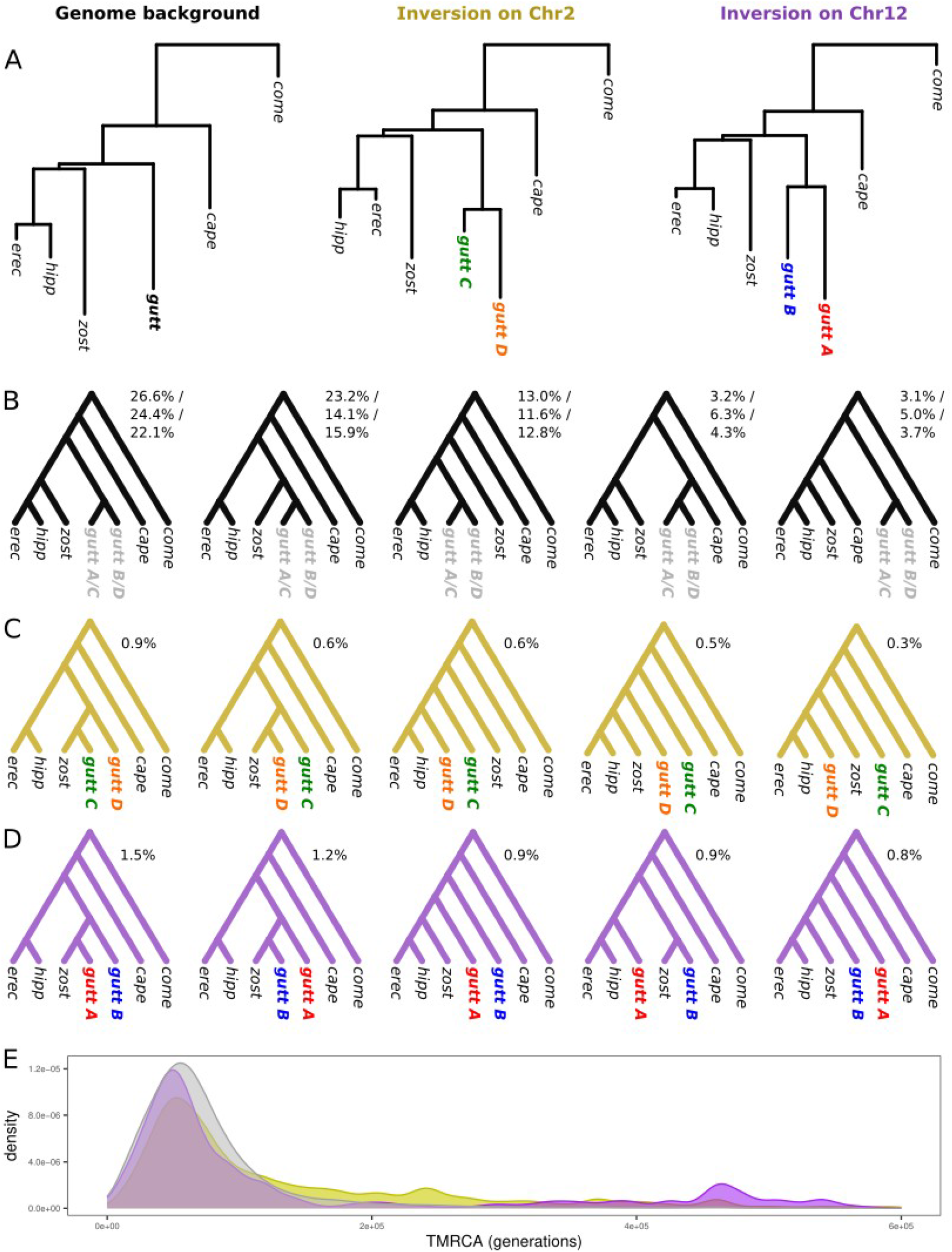
A) Maximum likelihood phylogenies for different regions of the seahorse genome. B) The five most predominant 7-taxon topologies identified with Twisst in the genome background as well as in the inversions. Relative proportions (percentage among all topologies in a specific region) are shown for the genome background (top), for B2 (middle, leaves *gutt C* and *gutt D*) and for B12 (bottom, leaves *gutt A* and *gutt B*). C-D) The five most common topologies in B2 (C) and in B12 (D) which did not place alternate haplotypes as sister lineages. E) Distribution of coalescent times estimated with *tsdate* (v0.1.5) in the genome background (grey), on Chr2 (yellow) and on Chr12 (purple) between a DD/AA individual (HguttLi4) and a CC/BB individual (HguttGa7). Abbreviations: *Come*: *H. comes. Cape*: *H. capensis. Zost*: *H. zosterae. Hipp*: *H. hippocampus. Erec*: *H. erectus*.

Topology weighting conducted using Twisst (Martin & Van Belleghem, 2017) revealed that the most predominant topologies - both in the genome background as well as in the inversions - were consistent with the currently accepted genome-wide phylogeny of the genus (Li et al., 2021; Stiller et al., 2022). However, they simultaneously showed high levels of incomplete lineage sorting (ILS) near the ancestral nodes of the genus. If one of the two haplotypes on either Chr2 or Chr12 had been introduced through introgression from another species, we would have expected the tree topologies in this region to differ from genome-wide topologies. In this case, a *H. guttulatus* individual carrying the introgressed haplotype would have grouped closer to the donor species and not with *H. guttulatus* of the alternate haplotype. The topologies that did not group alternate haplotypes together, amounted to 2.2% and 15.7% of topologies in the B2 and B12 regions, respectively (Fig. 4E & 4F). Even when the two *H. guttulatus* haplotypes were not in the same grouping, they were not placed in distant positions (i.e. they still occurred in the same sub-branch). These topologies did not show any particular haplotype tending to group with a potential donor species, and both haplotypes showed shifted positions, most probably due to ILS. Given these findings, we conclude that the two inversion polymorphisms in long-snouted seahorses were not likely introduced through introgression with a closely related species.

Our results indicate that the B2 and B12 polymorphisms are most likely explained by chromosomal inversion events that took place within the *H. guttulatus* lineage. These inversions have been maintained as intraspecific polymorphisms for a long enough period of time for divergence to have accumulated between haplotypes. The divergence between A and B haplotypes on Chr12 was particularly high, since *D_XY_* values for comparisons between opposite homokaryotes ranged between 0.69 and 0.76% (first and third quartiles). Absolute divergence between opposite B2 homokaryotes was slightly lower and showed more variance (first to third quartile, 0.36 to 0.57%) (Supplementary Figure S9). TMRCA inferred in the B12 region were also higher than for the B2 region, and consensus phylogenies constructed with PhyML showed a deeper split for haplotypes A and B (Fig. 4C) than for C and D (Fig. 4B). We conclude that the inversions probably do not have the same age, and that the B12 polymorphism emerged before B2. Furthermore, it should be noted that divergence might have been underestimated in our study due to reference bias, e.g. reads from an AA individual were mapped to our B reference (Hgutt_V1).

In addition to these inversions, other intra- and inter-chromosomal rearrangements were evidenced in alignments between our reference assembly (Hgutt_V1), HguttRefA and the *H. erectus* assembly. Due to the abundance of these structural rearrangements, we were not able to determine the ancestral arrangement for inversions B2 and B12 through comparison with the *H. erectus* assembly. These analyses showed that *H. erectus* chromosomes Chr11 and Chr12 were fused in the HguttRefA assembly to form chromosome JAOYMQ010000004.1 (Supplementary Figure S11). This could potentially indicate that the A haplotype (carried by HguttRefA) on Chr12 is involved in a chromosomal fusion with Chr11 (contrary to the B haplotype), or alternatively, that this might represent a technical artefact of the HguttRefA assembly, which is based on short-read data. As for B2, we speculate that the discontinuous signal of high *F_ST_* associated with this inversion (Fig. 2C) might be due to additional rearrangements that have altered the collinearity between the C and D haplotypes. The D haplotype might present a different intra-chromosomal structure to the C haplotype, resulting in a heterogeneous divergence landscape when mapped to the C reference. However, our analyses did not allow us to confirm this hypothesis, since both *H. guttulatus* assemblies were CC homokaryotes. Alternatively, regions of low *F_ST_* and *D_XY_* could be due to local erosion of the divergence between haplotypes through recombination.

We wished to test for gene flux between inverted and non-inverted haplotypes, which can be expected in the form of double crossover events or gene conversion. Plotting PC1 coordinates from local PCA along inversion regions allowed us to locate those genome windows that showed the three-cluster pattern typical of inversions (Fig. 5A and 5D). B12 showed a classical triple structure at a large scale, whereas B2 showed more “choppy” patterns, with long stretches of “irregular” patterns separating three-cluster windows. For both inversions, these irregular windows that did not show three groups were regions that showed a breakdown of inversion structure. We used ARG-inferred trees to determine local ancestry along both inversions (Fig. 5B, 5C & 5E). We determined which trees showed the expected topology of opposite haplotypes grouping in two different branches (Fig. 5B), and which trees showed a discordant topology (Fig. 5C). Trees which grouped opposite haplotypes in the same branch indicated regions which were locally introgressed with a tract from the alternate haplotype. This allowed us to perform chromosome painting for each non-recombinant block associated with a particular tree. This approach revealed large-scale haplotypes corresponding to inverted and non-inverted ancestries, while simultaneously showing local ancestry variation at a finer scale. Chromosome painting strongly reflected the patterns observed in local PCA and *F_ST_* landscapes (see Supplementary Figures S14, S15a and S15b for the full dataset). For lower quality samples, we found evidence for phasing errors in the form of switches between maternal and paternal haplotypes, which were visible in heterokaryotic samples. These errors did not prevent us from locating introgressed tracts in homokaryotes, which spanned up to ∼100 kb for B12 and up to ∼500 kb for B2.

**Fig. 5.**
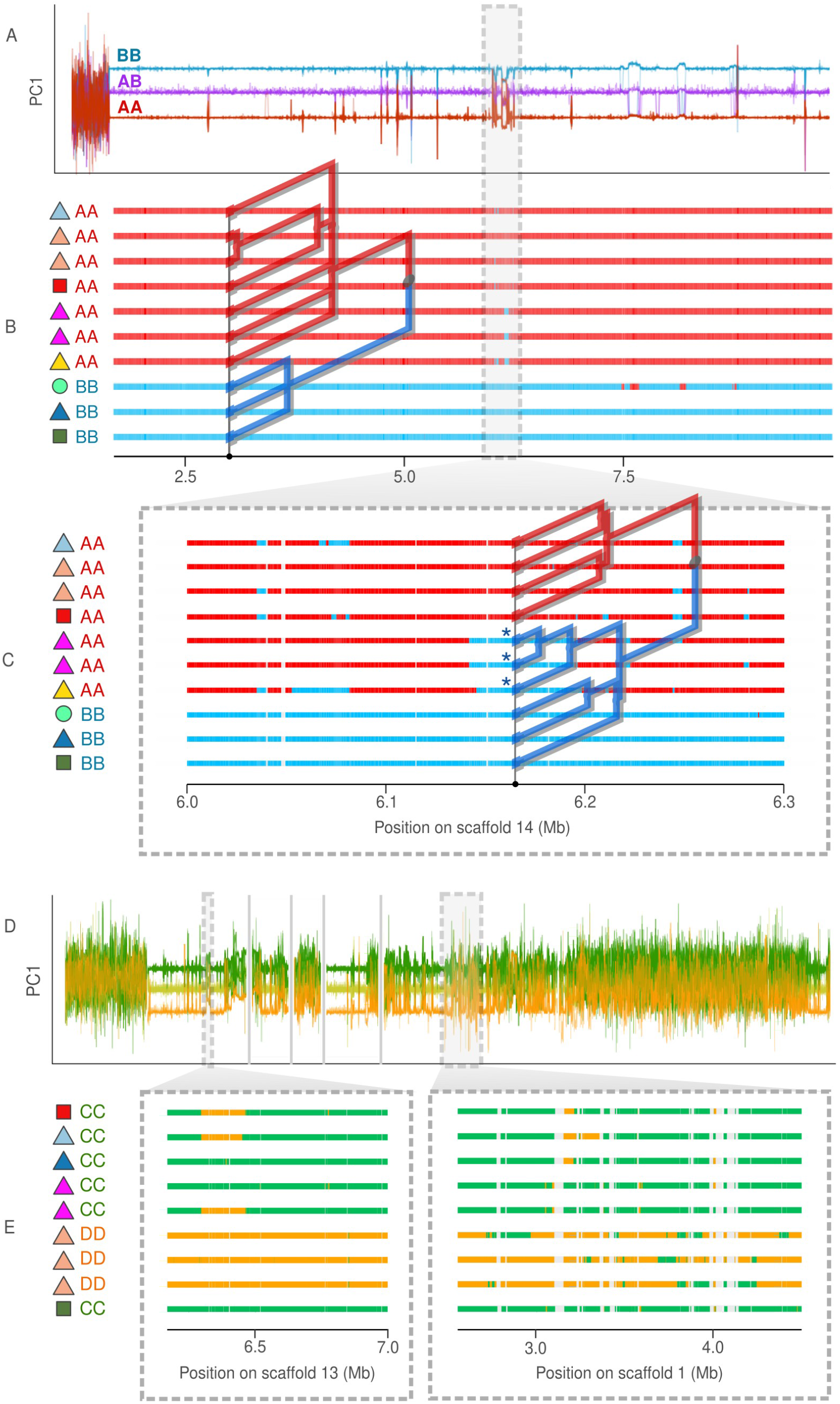
Gene flux between the inverted and non-inverted haplotypes on Chr12 (A-C) and on Chr2 (D-E). A & D) PC1 coordinates from local PCA plotted for non-overlapping 5 kb window along the inversions. Individual lines represent samples and are coloured according to inversion genotype. Vertical grey lines indicate the edges of scaffolds composing Chr2. B & C & E) Chromosome painting showing local ancestry within inversion regions as determined by our ARG-based approach. The superposed trees were constructed with *tsinfer* (v0.3.0) and illustrate how local ancestry at a given position was determined for each leaf. Horizontal coloured bars represent a subsample of all haploid chromosomes and colours indicate inverted or non-inverted ancestry. Symbols on the left of each chromosome indicate sampling location and habitat type (see Fig. 1A), as well as large-scale inversion genotype. C) Enlarged view of an introgressed region of B12, where certain A haplotypes (red) locally show B ancestry (blue). The tree at position 6.175 Mb shows that three A alleles (marked by asterisks) are grouped in the same branch as the B alleles. E) Enlarged views of two regions on Chr2 showing different levels of introgression.

## Discussion

We present the first genome-wide study of intraspecific diversity in *H. guttulatus*, investigating multiple aspects of population structure related to geography, habitat type and genome architecture. We confirmed the presence of two geographical lineages in the Atlantic and marine-lagoon ecotypes in the Mediterranean Sea (Riquet et al., 2019). In the latter region, we observed that marine and lagoon ecotypes show low levels of genome-wide differentiation, while exhibiting substantial frequency differences at two megabase-scale chromosomal inversions. We characterise the origin of these SVs and discuss the possible mechanisms responsible for their long-term maintenance. Lastly, we found evidence for gene flux between inverted alleles and address its role in shaping the dynamics and evolutionary fate of seahorse inversions.

### Evolutionary origin of seahorse inversions

Our results indicate that inversions B2 and B12 have been maintained as polymorphisms for hundreds of thousands of generations, that is, well beyond the mean coalescence time inferred within populations (<100k generations). Estimating coalescence times with *tsdate* is known to be hampered by technical limitations (Brandt et al., 2022) that prevented our ability to precisely determine and compare the ages of B2 and B12. Among the reported limitations, is the observation that *tsdate* tends to underestimate the oldest coalescence times. It is therefore conservative to assume that the ages we inferred for the inverted alleles greatly predate average within-population coalescence times. Consistently, these regions also showed remarkably high levels of raw divergence compared to the rest of the genome (*D_XY_* up to 1%, *vs.* maximum of 0.25% elsewhere). Moreover, the levels of nucleotide diversity associated with each haplotype were, given their respective frequencies, consistent with genome-wide average values. It is therefore likely that sufficient time has elapsed since the appearance of the inverted haplotypes for them to have progressed towards mutation-drift equilibrium.

Our finding that the two seahorse inversions represent ancient polymorphisms thus raises the question of their origin. One way that a divergent inverted haplotype may be introduced is through hybridisation with a closely related species (e.g., Hsieh et al., 2019; Jay et al., 2018). We did not find evidence for introgressed ancestry at either of the inversions, since topology analysis did not support similarity between inversion haplotypes and any potential donor species (Fig. 4 B-F). Instead, all major sources of genealogical conflict could be explained by incomplete lineage sorting (ILS) occurring at deep nodes in the *Hippocampus* genus phylogeny. Another scenario that could have given rise to similar heterogenous divergence landscapes, is secondary contact between previously isolated, divergent lineages. In this case, the inversions could have resisted re-homogenisation by gene flow due to recombination suppression (Lundberg et al., 2023; Rafajlović et al., 2021; Yeaman, 2013). It could have been expected that an erosion of past differentiation outside of the inversions would remain detectable by the presence of short divergent haplotypes produced by recombination between diverged ancestries. However, we did not find any such regions of deep coalescence comparable to that found in the inverted regions. It is thus most likely that the inversions have emerged within the *H. guttulatus* lineage and that they have remained polymorphic for long periods of time.

Molecular mechanisms facilitating the emergence of inversions have been studied in detail using long-read sequencing in deer mice, showing that inversion breakpoints tend to occur in centromeric and telomeric regions and to be flanked by LIRs (Harringmeyer & Hoekstra, 2022). Here, we were able to directly demonstrate, using linked-read data, that the 8.2 Mb inversion B12 occurs near a chromosome extremity and has its breakpoints in an approximately 1 kb LIR bearing a Tan9 monomer. Tandem repeats containing Tan9 are widespread throughout the *H. guttulatus* genome and may therefore facilitate ectopic recombination, leading to an increased rate of formation of new structural variants. Although we could not perform such a detailed analysis for the breakpoint regions of B2, it would be interesting to test whether the emergence of this inversion has also been promoted by the presence of recombinogenic elements (Wang & Leung, 2006), such as Tan9 LIRs.

### Maintenance of polymorphisms B2 and B12

The long-term persistence of ancient inversion polymorphisms has long been a topic of interest in evolutionary biology. In their review outlining the different life-history stages of inversions, Faria et al. (2019b) discuss the evolutionary changes that an inversion might undergo from the moment of its appearance to its loss or fixation. By placing our results in this framework, we sought to distinguish between the mechanisms that might maintain the two inversion polymorphisms in our system, and provide possible explanations for their respective patterns. We propose that the B12 polymorphism is most comparable in its characteristics to Type I inversion polymorphisms, while B2 probably calls on the interaction of forces linked both to Type I and Type II polymorphisms.

The B12 polymorphism corresponds to the genomic island identified by Riquet et al. (2019) (Supplementary Figure S16) as the main region differentiating geographical lineages in the Atlantic and ecotypes in the Mediterranean Sea. B12 shows local fixation for a given haplotype within almost all populations (Fig. 3F), in accordance with the pattern expected for Type I polymorphisms (Faria et al. 2019b). In this scenario, within-population polymorphism is not maintained, but among-population variation is conferred by divergent selection on alternate haplotypes. Given the differential fixation of B12 haplotypes between adjacent lagoon (A haplotype) and marine (B haplotype) sites in the Mediterranean Sea, it is possible that the inversion is either underdominant, under negative epistasis, or affected by selection for locally favoured alleles. Regardless of the specific form of selection at work, lack of recombination between inverted haplotypes ensures that the genes in the A and B haplotypes are inherited together, collectively forming a barrier to gene flow. Over a long period of time, the accumulation of new mutations between inverted haplotypes leads to high levels of divergence, as is reflected by high *D_XY_* values observed between AA and BB homokaryotes (Fig. 2B). Although there is little opportunity for gene flow between alternate haplotypes, recombination can proceed normally within every population that is fixed for a given haplotype, since crossing over is unimpeded in homokaryotes. In this way, it has been suggested that the accumulation of mutation load in inverted regions would not be substantially higher than in the collinear genome (Berdan et al. 2021). Besides being a theoretical prediction, empirical examples are provided by inversions in deer mice that show local fixation and lack of mutation load (Harringmeyer & Hoekstra, 2022), as well as inversions in sunflower populations fixed for one arrangement having lower mutation load compared to polymorphic populations (Huang et al., 2022). We can therefore speculate that the long-term subsistence of B12 inversion as a Type I polymorphism has not been hampered by the accumulation of deleterious mutations.

Regarding the possible selective mechanisms at play, lower fitness of heterozygotes (i.e., underdominance) is expected at Type I polymorphisms over the long term, since divergent selection and independent evolution of haplotypes may lead to the accumulation of DM incompatibilities within the inversion. We identified only seven samples (out of 112) that carried the AB genotype at B12, possibly pointing to a deficit of heterokaryotes at the inter-population level. These seven samples originated from only two populations (Port d’Albret, Bay of Biscay, and Varna, Black Sea) where both haplotypes A and B were present and where heterokaryotes were locally common (4 out of 11 and 3 out of 5 samples, respectively). If B12 heterokaryotes are selected against, these populations might represent underdominant clines, as was put forward by Riquet et al. (2019) for polymorphic populations in the contact zone in the Basque country. Furthermore, we also observe a breakdown of the perfect association that exists between habitat and B12 genotype outside of the Mediterranean Sea. Atlantic and Black Sea populations do not carry different inversion haplotypes in marine and lagoon environments, as already observed by Riquet et al. (2019). This partial decoupling from habitat type could be explained by predominantly intrinsic rather than extrinsic incompatibility between A and B. The difference in patterns observed between the Atlantic and the Mediterranean Sea may also involve epistatic interactions with loci on other chromosomes, such as the B2 inversion (see below). In any case, it remains possible that the B12 inversion is to some extent directly involved in differential adaptation to marine and lagoon environments, particularly in the Mediterranean Sea.

In contrast to B12, inversion B2 is maintained as a polymorphism in Mediterranean lagoons, rather than being differentially fixed between marine and lagoon environments. Classical work that has addressed the long-term maintenance of inversion polymorphisms has evoked an important role of balancing selection in maintaining within-population variation (reviewed in, e.g. Llaurens et al., 2017; Wellenreuther & Bernatchez, 2018). This results in Type II polymorphisms *sensu* Faria et al. (2019b), which could explain the distribution of the B2 polymorphism in the Mediterranean Sea. The preponderance of CD genotypes (Th: 12 out of 36; Mu: 3 out of 5; Bz: 5 out of 5) in Mediterranean lagoons may indeed indicate increased fitness in heterokaryotes. The limited distribution of the B2 polymorphism combined with intermediate haplotype frequencies at the populational level, might result in a reduced effective population size and a higher mutation load associated with both haplotypes (Berdan et al., 2021). Additionally, the B2 inversion is nearly three times longer than B12, suggesting that it is more likely to have captured recessive deleterious mutations that are inherited as a single block (Connallon & Olito, 2022). For such inversions that suffer from high mutation load, theory predicts that polymorphism could be maintained by associative overdominance (Berdan et al., 2021), which is also supported by empirical studies in insects (Jay et al., 2021; Yang et al., 2002). For these reasons, we suggest that compensation of the mutation load in heterokaryotes ensures the persistence of the B2 polymorphism in Mediterranean lagoons.

The simultaneous action of forces associated with both Type I and Type II polymorphisms has been shown to potentially give rise to a range of different inversion equilibrium frequencies (Faria et al., 2019b). For instance, selection for heterokaryotes combined with local adaptation can result in polymorphism in one environment and fixation in the other. It is therefore plausible that a similar commixture of forces could be at work to explain the fixation of B2 haplotype C in marine *H. guttulatus* populations, while polymorphism is maintained in Mediterranean lagoons. If the B2 polymorphism is maintained by associative overdominance, we could ask why the D haplotype is not present outside of Mediterranean lagoons. The answer may potentially be connected to local adaptation alleles that are carried in the haplotypes of B2, or in epistatic association with B12. If D provides local advantage in lagoons, or if D/A combinations are favoured over C/A, it could be maintained as a polymorphism (but not fixed) in these environments. On the other hand, D would be outcompeted elsewhere due to lack of selective advantage and high mutation load. Although current knowledge does not allow us to answer all questions regarding the maintenance of the B2 polymorphism, we suspect that it involves an interaction of forces which could include divergent selection of differing strengths and forms of balancing selection (e.g. associative overdominance). All of these processes should also be affected by the level and direction of effective migration connecting different populations.

### Long-term fate of inversion polymorphisms in the long-snouted seahorse

Inversion haplotypes are not expected to persist for an undefined period of time if the evolutionary mechanisms underlying polymorphism are subject to change. An inversion might eventually fix one arrangement throughout the species range, or alternatively, couple with other genomic components of reproductive isolation and fix differentially between incipient species (Faria et al. 2019b). The long-term fate of an inversion thus depends on the processes that drive its dynamics throughout its lifetime. For example, Berdan et al. (2021) studied the “feedback loop” between allelic content and haplotype frequency, caused by the accumulation of deleterious mutations, which in turn affects the frequencies of the different karyotypes. Another process that can influence the long-term fate of an inversion, is the exchange of genetic material between alternate haplotypes through gene conversion or double crossover during meiosis (i.e., gene flux). In the current section we discuss these various points and attempt to make predictions about the long-term fates of the B2 and B12 polymorphisms segregating in *H. guttulatus*.

Theoretical (reviewed in, e.g. Hoffmann & Rieseberg, 2008) and empirical studies (e.g. Huang et al., 2020; Lohse et al., 2015; Noor et al., 2001) have found that, under certain conditions, the presence of (Type I) inversion polymorphisms may facilitate speciation. In fish, multiple empirical studies have highlighted the role of chromosomal inversions in local adaptation, ecotype formation and speciation (Berg et al., 2016; Cayuela et al., 2020; Jones et al., 2012; Le Moan et al., 2021; Matschiner et al., 2022; Pettersson et al., 2019; Tigano et al., 2021). Mediterranean ecotypes of long-snouted seahorse show differential fixation at inversion B12, which is under divergent selection (Type I polymorphism, see previous section) and could eventually facilitate speciation between marine and lagoon ecotypes. We found evidence for very low levels of gene flux taking place in this inversion, suggesting that gene exchange in heterokaryotes is either rare or selected against. Since exchanged segments were generally longer than a few kb and occurred in the middle of the inversion, they most likely resulted from double crossovers rather than gene conversion, in line with theoretical expectations for long inversions (Navarro et al., 1997), but contrary to what was found in cod where gene conversion predominates (Matschiner et al., 2022). Introgressed tracts of B ancestry within A haplotypes were found across the species range, but were nonetheless generally small in size (up to 50 kb) (Supplementary Fig. S14). This suggests that introgression took place a long time ago and that the remaining tracts we observe represent recombined segments that have passed the filter of selection. Evidence for introgression in the opposite direction (A into B) was observed only in the Black Sea population, where both haplotypes segregate and opportunities for gene flux are increased. These tracts of A ancestry within B haplotypes were slightly longer (up to 100 kb) and more numerous, potentially suggesting more recent gene flux.

Whatever the direction of gene flux, the fact that introgressed segments occupy only few and relatively narrow genomic regions in B12, argues for selection acting against introgressed ancestry. This is consistent with the existence of multiple selected mutations that have accumulated within the inversion over the long term (e.g., DM incompatibilities; Navarro & Barton, 2003). If B12 is responsible for strong underdominance and further becomes involved in reinforcement by coupling with premating isolating mechanisms, it may eventually strengthen reproductive isolation between lagoon and marine populations (Faria et al., 2019b). It can be noted that some level of genome-wide differentiation is already observed between Mediterranean ecotypes, thus indicating a significant reduction in effective gene flow at a small spatial scale relative to dispersal. However, the contribution of B12 to gene flow reduction is more uncertain in the Atlantic and the Black Sea. Although a weak association has been observed between B12 and habitat type along the Portuguese coasts, no associated genetic structure was detected in the genomic background (Riquet et al., 2019). Surprisingly, we found an increase in the frequency of the A haplotype in Faro lagoon over the last ten years, but further study will be necessary to confirm whether this observation is a true temporal trend, an indication of cryptic microhabitat variation, or sampling noise.

If the B12 polymorphism lends itself more to speciation than to universal fixation, the long-term fate of the B2 (i.e. Type II) polymorphism is less straightforward to predict. Since the frequencies of this inversion result from a balance between different processes, namely a form of divergent along with balancing selection, it is unclear whether B2 will eventually undergo differential fixation between habitats, or universal fixation of one arrangement. Furthermore, our large-scale assembly of Chr2 is still only partly resolved, and we cannot rule out the possibility that additional chromosomal rearrangements have affected its divergence landscape and evolutionary trajectory. For example, multiple inversions occurring in the same region might have extended the block of high LD, as is the case for adjacent (Jay et al., 2021) or nested inversions (Maggiolini et al., 2020). In contrast to B12, we also found ample evidence for the erosion of divergence in B2 through gene flux, as illustrated by the detection of many introgressed segments (Supplementary Fig. S15a and S15b) and “suspension bridge” patterns in the *F_ST_* landscape (Fig. 2C). The intensity of gene flux was clearly heterogeneous along this chromosome, since we found evidence for relatively recent double crossover events (i.e. larger introgressed segments than in B12) as well as one completely eroded region, where low *F_ST_* values (comparable to genome background) are bordered by high differentiation segments. Gene flux is therefore likely to impact the fate of this inversion, especially if the dynamics of B2 are largely driven by mutation load, as we discussed in the previous section. Recombination between haplotypes might favour the removal of deleterious mutations, as has been shown through simulations, where even low levels of gene conversion were sufficient to mitigate mutation load (Berdan et al., 2021). Given enough time, gene flux between B2 haplotypes could thus weaken the associative overdominance that is suspected of maintaining them.

The framework laid out by Faria et al. (2019b) does not specifically consider the dynamics of multiple inversions co-existing in the same system, but leaves such interactions as an outstanding question. The presence of other polymorphic inversions could indeed impact the establishment, maintenance, and long-term fate of a given inversion, for example through epistatic interactions or coupling. The two inversion polymorphisms segregating in *H. guttulatus* might present a case study for such potential interactions.

Timing and demographic context could be important in determining these dynamics, since these factors could influence the allelic contents of a new inversion and its probability to establish. For example, if the B12 polymorphism was already established and fixed between certain populations by the time B2 emerged, it could have affected the trajectory of the new B2 polymorphism. Furthermore, it is possible that there are epistatic interactions between B2 and B12, which co-occur in lagoon populations. Epistatic interactions could potentially explain why there is a strong association between B12 and habitat type in the Mediterranean, while the absence of B2 polymorphism in the Atlantic would only produce weak associations of B12 with habitat. The question therefore remains as to whether B2 contributes to speciation between marine and lagoon lineages in the Mediterranean through coupling with B12. Future directions for research could address these aspects by focusing on polymorphic populations, attempting to characterise the fitnesses of different karyotypes in different environments, and by quantifying mutation load. Studying gene flux may also reveal more about the loci contained in each arrangement, as we should expect erosion of divergence in regions carrying selectively neutral or disadvantageous mutations, and maintenance in regions that are under divergent selection.

## Acknowledgements

The data were partly produced and analysed with the support of the *GenSeq* genotyping and sequencing platform and the *MBB* Montpellier Bioinformatics Biodiversity platform, both being supported by ANR program “Investissements d’avenir” (ANR-10-LABX-04-01). We thank Rémy Dernat and Khalid Belkhir for their assistance in data storage, management and processing. DNA extraction of historical samples was carried out in a clean lab at ISEM, Montpellier (Plateforme d’ADN dégradé, LabEx CeMEB). We thank the Montpellier GenomiX platform for constructing the reference genome’s 10X Chromium library. We are grateful to the National Museum of Natural History (MNHN Paris, Agnès Dettaï) and colleagues who provided us with samples, as well as those who facilitated or participated in sampling: Jorge Palma and Rita Castilho (CCMAR, Portugal), Cristina Mena (Hippocampus association, Spain), Patrick Louisy (peau-bleue association, CPIE Bassin de Thau, France), Lucy Woodall (University of Oxford, UK), and citizens from the Sète region who provided dried seahorses samples. We thank Huixian Zhang of the Key Laboratory of Tropical Marine Bio-Resources and Ecology, South China Sea Institute of Oceanology, for providing the *Hippocampus erectus* reference genome. Finally, we thank Thomas Broquet for his valuable inputs on the manuscript. This work was supported by the ANR grant CoGeDiv ANR-17-CE02-0006-01 to PAG, and by a Languedoc-Roussillon Region “Chercheur(se)s d’avenir” grant to NB (Connect7 project), with the support of the Occitanie Regional Council’s program «Key challenge BiodivOc». Mobility during the project was partly funded by an ESEB Godfrew Hewitt Mobility Award as well as a Laura Corrigan conservation grant.

## Data accessibility

Our reference genome assembly will be deposited in GenBank, and raw sequence reads from whole-genome re-sequencing data (112 individuals) will be deposited in the GenBank Sequence Read Archive under the accession code BioProject ID XXXXXX. Supplementary File S2 contains scripts and commands used.

## Author contributions

LM, BG, NB and P-AG conceived the study. LM wrote the manuscript with inputs from all co-authors. Sampling was conducted by NB, FR, PB, RC and P-AG. CA performed HMW DNA extraction and preparation for the reference genome 10X library. DNA extraction for WGS libraries was performed by LM, PB, and FR. Aspects of degraded/historical DNA extraction and analysis were handled by AF, CDS, BG and LM. WGS library preparation was done by LM and FC. FC, AB and ED were responsible for haplotagging library construction and sequencing. P-AG produced the reference genome and analysed linked-read sequencing data. PB provided scripts for genotyping high-coverage samples. LM performed all other bioinformatics and population genomic analyses. P-AG managed financial support.

## Notes

### Competing Interest Statement

The authors have declared no competing interest.

## References

1. Barry, P., Broquet, T., & Gagnaire, P.-A. (2020). Life tables shape genetic diversity in marine fishes.

2. Barry, P., Broquet, T., & Gagnaire, P.-A. (2022). Age-specific survivorship and fecundity shape genetic diversity in marine fishes. Evolution letters, 6(1), 46–62.

3. Benson, G. (1999). Tandem repeats finder: A program to analyze DNA sequences. Nucleic Acids Research, 27(2), 573. https://doi.org/10.1093/nar/27.2.573

4. Berdan, E. L., Blanckaert, A., Butlin, R. K., & Bank, C. (2021). Deleterious mutation accumulation and the long-term fate of chromosomal inversions. PLOS Genetics, 17(3), e1009411. https://doi.org/10.1371/journal.pgen.1009411

5. Berg, P. R., Star, B., Pampoulie, C., Sodeland, M., Barth, J. M. I., Knutsen, H., Jakobsen, K. S., & Jentoft, S. (2016). Three chromosomal rearrangements promote genomic divergence between migratory and stationary ecotypes of Atlantic cod. Scientific Reports, 6(1), Article 1. https://doi.org/10.1038/srep23246

6. Butlin, R. K. (2005). Recombination and speciation. Molecular Ecology, 14(9), 2621–2635. https://doi.org/10.1111/j.1365-294X.2005.02617.x

7. Cabanettes, F., & Klopp, C. (2018). D-GENIES: Dot plot large genomes in an interactive, efficient and simple way. PeerJ, 6, e4958. https://doi.org/10.7717/peerj.4958

8. Camacho, C., Coulouris, G., Avagyan, V., Ma, N., Papadopoulos, J., Bealer, K., & Madden, T. L. (2009). BLAST+: Architecture and applications. BMC Bioinformatics, 10(1), 421. https://doi.org/10.1186/1471-2105-10-421

9. Campbell, M. A., Anderson, E. C., Garza, J. C., & Pearse, D. E. (2021). Polygenic Basis and the Role of Genome Duplication in Adaptation to Similar Selective Environments. Journal of Heredity, 112(7), 614–625. https://doi.org/10.1093/jhered/esab049

10. Campos, P. F., & Gilbert, T. M. P. (2012). DNA extraction from formalin-fixed material. Methods in Molecular Biology (Clifton, N.J.), 840, 81–85. https://doi.org/10.1007/978-1-61779-516-9_11

11. Cayuela, H., Rougemont, Q., Laporte, M., Mérot, C., Normandeau, E., Dorant, Y., Tørresen, O. K., Hoff, S. N. K., Jentoft, S., Sirois, P., Castonguay, M., Jansen, T., Praebel, K., Clément, M., & Bernatchez, L. (2020). Shared ancestral polymorphisms and chromosomal rearrangements as potential drivers of local adaptation in a marine fish. Molecular Ecology, 29(13), 2379-2398. https://doi.org/10.1111/mec.15499

13. Challis, R., Richards, E., Rajan, J., Cochrane, G., & Blaxter, M. (2020). BlobToolKit – Interactive Quality Assessment of Genome Assemblies. G3 Genes|Genomes|Genetics, 10(4), 1361-1374. https://doi.org/10.1534/g3.119.400908

14. Charlesworth, B. (2009). Effective population size and patterns of molecular evolution and variation. Nature Reviews Genetics, 10(3), 195–205. https://doi.org/10.1038/nrg2526

15. Chen, S., Zhou, Y., Chen, Y., & Gu, J. (2018). fastp: An ultra-fast all-in-one FASTQ preprocessor. Bioinformatics, 34(17), i884–i890. https://doi.org/10.1093/bioinformatics/bty560

16. Cheng, C., White, B. J., Kamdem, C., Mockaitis, K., Costantini, C., Hahn, M. W., & Besansky, N. J. (2012). Ecological Genomics of Anopheles gambiae Along a Latitudinal Cline: A Population-Resequencing Approach. Genetics, 190(4), 1417–1432. https://doi.org/10.1534/genetics.111.137794

17. Connallon, T., & Olito, C. (2022). Natural selection and the distribution of chromosomal inversion lengths. Molecular Ecology, 31(13), 3627–3641. https://doi.org/10.1111/mec.16091

18. Danecek, P., Auton, A., Abecasis, G., Albers, C. A., Banks, E., DePristo, M. A., Handsaker, R. E., Lunter, G., Marth, G. T., Sherry, S. T., McVean, G., Durbin, R., & 1000 Genomes Project Analysis Group. (2011). The variant call format and VCFtools. Bioinformatics, 27(15), 2156-2158. https://doi.org/10.1093/bioinformatics/btr330

19. Danecek, P., Bonfield, J. K., Liddle, J., Marshall, J., Ohan, V., Pollard, M. O., Whitwham, A., Keane, T., McCarthy, S. A., Davies, R. M., & Li, H. (2021). Twelve years of SAMtools and BCFtools. GigaScience, 10(2), giab008. https://doi.org/10.1093/gigascience/giab008

20. Delaneau, O., Zagury, J.-F., Robinson, M. R., Marchini, J. L., & Dermitzakis, E. T. (2019). Accurate, scalable and integrative haplotype estimation. Nature Communications, 10(1), Article 1. https://doi.org/10.1038/s41467-019-13225-y

21. Doyle, J. J., & Doyle, J. L. (Éds.). (1987). A rapid DNA isolation procedure for small quantities of fresh leaf tissue. PHYTOCHEMICAL BULLETIN.

22. Faria, R., Chaube, P., Morales, H. E., Larsson, T., Lemmon, A. R., Lemmon, E. M., Rafajlović, M., Panova, M., Ravinet, M., Johannesson, K., Westram, A. M., & Butlin, R. K. (2019a). Multiple chromosomal rearrangements in a hybrid zone between Littorina saxatilis ecotypes. Molecular Ecology, 28(6), 1375–1393. https://doi.org/10.1111/mec.14972

23. Faria, R., Johannesson, K., Butlin, R. K., & Westram, A. M. (2019b). Evolving Inversions. Trends in Ecology & Evolution, 34(3), 239–248. https://doi.org/10.1016/j.tree.2018.12.005

24. Faria, R., & Navarro, A. (2010). Chromosomal speciation revisited: Rearranging theory with pieces of evidence. Trends in Ecology & Evolution, 25(11), 660–669. https://doi.org/10.1016/j.tree.2010.07.008

25. Ferrari, G., Atmore, L. M., Jentoft, S., Jakobsen, K. S., Makowiecki, D., Barrett, J. H., & Star, B. (2022). An accurate assignment test for extremely low-coverage whole-genome sequence data. Molecular Ecology Resources, 22(4), 1330–1344. https://doi.org/10.1111/1755-0998.13551

26. Flynn, J. M., Hubley, R., Goubert, C., Rosen, J., Clark, A. G., Feschotte, C., & Smit, A. F. (2020). RepeatModeler2 for automated genomic discovery of transposable element families. Proceedings of the National Academy of Sciences, 117(17), 9451–9457. https://doi.org/10.1073/pnas.1921046117

27. Funk, E. R., Mason, N. A., Pálsson, S., Albrecht, T., Johnson, J. A., & Taylor, S. A. (2021). A supergene underlies linked variation in color and morphology in a Holarctic songbird. Nature Communications, 12(1), Article 1. https://doi.org/10.1038/s41467-021-27173-z

28. Gould, B. A., Chen, Y., & Lowry, D. B. (2017). Pooled ecotype sequencing reveals candidate genetic mechanisms for adaptive differentiation and reproductive isolation. Molecular Ecology, 26(1), 163–177. https://doi.org/10.1111/mec.13881

29. Guerrero, R. F., Rousset, F., & Kirkpatrick, M. (2012). Coalescent patterns for chromosomal inversions in divergent populations. Philosophical Transactions of the Royal Society B: Biological Sciences, 367(1587), 430–438. https://doi.org/10.1098/rstb.2011.0246

30. Guichard, A., Legeai, F., Tagu, D., & Lemaitre, C. (2022). MTG-Link: Leveraging barcode information from linked-reads to assemble specific loci (p. 2022.09.27.509642). bioRxiv. https://doi.org/10.1101/2022.09.27.509642

31. Guindon, S., Dufayard, J.-F., Lefort, V., Anisimova, M., Hordijk, W., & Gascuel, O. (2010). New Algorithms and Methods to Estimate Maximum-Likelihood Phylogenies: Assessing the Performance of PhyML 3.0. Systematic Biology, 59(3), 307–321. https://doi.org/10.1093/sysbio/syq010

32. Hager, E. R., Harringmeyer, O. S., Wooldridge, T. B., Theingi, S., Gable, J. T., McFadden, S., Neugeboren, B., Turner, K. M., Jensen, J. D., & Hoekstra, H. E. (2022). A chromosomal inversion contributes to divergence in multiple traits between deer mouse ecotypes. Science, 377(6604), 399–405. https://doi.org/10.1126/science.abg0718

33. Harringmeyer, O. S., & Hoekstra, H. E. (2022). Chromosomal inversion polymorphisms shape the genomic landscape of deer mice. Nature Ecology & Evolution, 1-15. https://doi.org/10.1038/s41559-022-01890-0

34. Haÿ, V., Mennesson, M. I., Dettaï, A., Bonillo, C., Keith, P., & Lord, C. (2020). Needlepoint non-destructive internal tissue sampling for precious fish specimens. Cybium: Revue Internationale d’Ichtyologie, 44(1), 73–79. https://doi.org/10.26028/cybium/2020-441-010

35. Hill, W. G., & Robertson, A. (1966). The effect of linkage on limits to artificial selection. Genetics Research, 8(3), 269–294. https://doi.org/10.1017/S0016672300010156

36. Hoffmann, A. A., & Rieseberg, L. H. (2008). Revisiting the Impact of Inversions in Evolution: From Population Genetic Markers to Drivers of Adaptive Shifts and Speciation? Annual review of ecology, evolution, and systematics, 39, 21–42. https://doi.org/10.1146/annurev.ecolsys.39.110707.173532

37. Hsieh, P., Vollger, M. R., Dang, V., Porubsky, D., Baker, C., Cantsilieris, S., Hoekzema, K., Lewis, A. P., Munson, K. M., Sorensen, M., Kronenberg, Z. N., Murali, S., Nelson, B. J., Chiatante, G., Maggiolini, F. A. M., Blanché, H., Underwood, J. G., Antonacci, F., Deleuze, J.-F., & Eichler, E. E. (2019). Adaptive archaic introgression of copy number variants and the discovery of previously unknown human genes. Science, 366(6463), eaax2083. https://doi.org/10.1126/science.aax2083

38. Huang, K., Andrew, R. L., Owens, G. L., Ostevik, K. L., & Rieseberg, L. H. (2020). Multiple chromosomal inversions contribute to adaptive divergence of a dune sunflower ecotype. Molecular Ecology, 29(14), 2535–2549. https://doi.org/10.1111/mec.15428

39. Huang, K., Ostevik, K. L., Elphinstone, C., Todesco, M., Bercovich, N., Owens, G. L., & Rieseberg, L. H. (2022). Mutation Load in Sunflower Inversions Is Negatively Correlated with Inversion Heterozygosity. Molecular Biology and Evolution, 39(5), msac101. https://doi.org/10.1093/molbev/msac101

40. Jay, P., Chouteau, M., Whibley, A., Bastide, H., Parrinello, H., Llaurens, V., & Joron, M. (2021). Mutation load at a mimicry supergene sheds new light on the evolution of inversion polymorphisms. Nature Genetics, 53(3), 288–293. https://doi.org/10.1038/s41588-020-00771-1

41. Jay, P., Leroy, M., Le Poul, Y., Whibley, A., Arias, M., Chouteau, M., & Joron, M. (2022). Association mapping of colour variation in a butterfly provides evidence that a supergene locks together a cluster of adaptive loci. Philosophical Transactions of the Royal Society B: Biological Sciences, 377(1856), 20210193. https://doi.org/10.1098/rstb.2021.0193

42. Jay, P., Whibley, A., Frézal, L., Rodríguez de Cara, M. Á., Nowell, R. W., Mallet, J., Dasmahapatra, K. K., & Joron, M. (2018). Supergene Evolution Triggered by the Introgression of a Chromosomal Inversion. Current Biology, 28(11), 1839–1845.e3. https://doi.org/10.1016/j.cub.2018.04.072

43. Jones, F. C., Grabherr, M. G., Chan, Y. F., Russell, P., Mauceli, E., Johnson, J., Swofford, R., Pirun, M., Zody, M. C., White, S., Birney, E., Searle, S., Schmutz, J., Grimwood, J., Dickson, M. C., Myers, R. M., Miller, C. T., Summers, B. R., Knecht, A. K.,…Kingsley, D. M. (2012). The genomic basis of adaptive evolution in threespine sticklebacks. Nature, 484(7392), Article 7392. https://doi.org/10.1038/nature10944

44. Keightley, P. D., & Jackson, B. C. (2018). Inferring the Probability of the Derived vs. The Ancestral Allelic State at a Polymorphic Site. Genetics, 209(3), 897–906. https://doi.org/10.1534/genetics.118.301120

44. Kelleher, J., Wong, Y., Wohns, A. W., Fadil, C., Albers, P. K., & McVean, G. (2019). Inferring whole-genome histories in large population datasets. Nature Genetics, 51(9), Article 9. https://doi.org/10.1038/s41588-019-0483-y

45. Kirkpatrick, M. (2010). How and Why Chromosome Inversions Evolve. PLOS Biology, 8(9), e1000501. https://doi.org/10.1371/journal.pbio.1000501

46. Kirkpatrick, M., & Barton, N. (2006). Chromosome inversions, local adaptation and speciation. Genetics, 173(1), 419–434. https://doi.org/10.1534/genetics.105.047985

47. Kirov, I., Gilyok, M., Knyazev, A., & Fesenko, I. (2018). Pilot satellitome analysis of the model plant, Physcomitrellapatens, revealed a transcribed and high-copy IGS related tandem repeat. Comparative Cytogenetics, 12(4), 493–513. https://doi.org/10.3897/CompCytogen.v12i4.31015

48. Korneliussen, T. S., Albrechtsen, A., & Nielsen, R. (2014). ANGSD: Analysis of Next Generation Sequencing Data. BMC Bioinformatics, 15(1), 356. https://doi.org/10.1186/s12859-014-0356-4

49. Korunes, K. L., & Noor, M. A. F. (2019). Pervasive gene conversion in chromosomal inversion heterozygotes. Molecular Ecology, 28(6), 1302–1315. https://doi.org/10.1111/mec.14921

50. Kulmuni, J., Butlin, R. K., Lucek, K., Savolainen, V., & Westram, A. M. (2020). Towards the completion of speciation: The evolution of reproductive isolation beyond the first barriers. Philosophical Transactions of the Royal Society B: Biological Sciences, 375(1806), 20190528. https://doi.org/10.1098/rstb.2019.0528

51. Lamichhaney, S., Fan, G., Widemo, F., Gunnarsson, U., Thalmann, D. S., Hoeppner, M. P., Kerje, S., Gustafson, U., Shi, C., Zhang, H., Chen, W., Liang, X., Huang, L., Wang, J., Liang, E., Wu, Q., Lee, S. M.-Y., Xu, X., Höglund, J.,…Andersson, L. (2016). Structural genomic changes underlie alternative reproductive strategies in the ruff (Philomachus pugnax). Nature Genetics, 48(1), Article 1. https://doi.org/10.1038/ng.3430

52. Le Moan, A., Bekkevold, D., & Hemmer-Hansen, J. (2021). Evolution at two time frames: Ancient structural variants involved in post-glacial divergence of the European plaice (Pleuronectes platessa). Heredity, 126(4), Article 4. https://doi.org/10.1038/s41437-020-00389-3

53. Li, C., Olave, M., Hou, Y., Qin, G., Schneider, R. F., Gao, Z., Tu, X., Wang, X., Qi, F., Nater, A., Kautt, A. F., Wan, S., Zhang, Y., Liu, Y., Zhang, H., Zhang, B., Zhang, H., Qu, M., Liu, S.,…Lin, Q. (2021). Genome sequences reveal global dispersal routes and suggest convergent genetic adaptations in seahorse evolution. Nature Communications, 12(1), Article 1. https://doi.org/10.1038/s41467-021-21379-x

54. Li, H. (2013). Aligning sequence reads, clone sequences and assembly contigs with BWA-MEM (arXiv:1303.3997). arXiv. http://arxiv.org/abs/1303.3997

55. Li, H. (2018). Minimap2: Pairwise alignment for nucleotide sequences. Bioinformatics, 34(18), 3094–3100. https://doi.org/10.1093/bioinformatics/bty191

56. Li, H., & Ralph, P. (2019). Local PCA Shows How the Effect of Population Structure Differs Along the Genome. Genetics, 211(1), 289–304. https://doi.org/10.1534/genetics.118.301747

57. Llaurens, V., Whibley, A., & Joron, M. (2017). Genetic architecture and balancing selection: The life and death of differentiated variants. Molecular Ecology, 26(9), 2430–2448. https://doi.org/10.1111/mec.14051

58. Lohse, K., Clarke, M., Ritchie, M. G., & Etges, W. J. (2015). Genome-wide tests for introgression between cactophilic Drosophila implicate a role of inversions during speciation. Evolution, 69(5), 1178–1190. https://doi.org/10.1111/evo.12650

59. Lowry, D. B., & Willis, J. H. (2010). A Widespread Chromosomal Inversion Polymorphism Contributes to a Major Life-History Transition, Local Adaptation, and Reproductive Isolation. PLOS Biology, 8(9), e1000500. https://doi.org/10.1371/journal.pbio.1000500

60. Lundberg, M., Mackintosh, A., Petri, A., & Bensch, S. (2023). Inversions maintain differences between migratory phenotypes of a songbird. Nature Communications, 14(1), Article 1. https://doi.org/10.1038/s41467-023-36167-y

61. Maggiolini, F. A. M., Sanders, A. D., Shew, C. J., Sulovari, A., Mao, Y., Puig, M., Catacchio, C. R., Dellino, M., Palmisano, D., Mercuri, L., Bitonto, M., Porubský, D., Cáceres, M., Eichler, E. E., Ventura, M., Dennis, M. Y., Korbel, J. O., & Antonacci, F. (2020). Single-cell strand sequencing of a macaque genome reveals multiple nested inversions and breakpoint reuse during primate evolution. Genome Research, 30(11), 1680–1693. https://doi.org/10.1101/gr.265322.120

62. Manni, M., Berkeley, M. R., Seppey, M., & Zdobnov, E. M. (2021). BUSCO: Assessing Genomic Data Quality and Beyond. Current Protocols, 1(12), e323. https://doi.org/10.1002/cpz1.323

63. Marion, S. B., & Noor, M. A. F. (2023). Interrogating the Roles of Mutation–Selection Balance, Heterozygote Advantage, and Linked Selection in Maintaining Recessive Lethal Variation in Natural Populations. Annual Review of Animal Biosciences, 11(1), 77–91. https://doi.org/10.1146/annurev-animal-050422-092520

64. Martin, S. H., & Van Belleghem, S. M. (2017). Exploring Evolutionary Relationships Across the Genome Using Topology Weighting. Genetics, 206(1), 429–438. https://doi.org/10.1534/genetics.116.194720

65. Matschiner, M., Barth, J. M. I., Tørresen, O. K., Star, B., Baalsrud, H. T., Brieuc, M. S. O., Pampoulie, C., Bradbury, I., Jakobsen, K. S., & Jentoft, S. (2022). Supergene origin and maintenance in Atlantic cod. Nature Ecology & Evolution, 6(4), Article 4. https://doi.org/10.1038/s41559-022-01661-x

66. McKenna, A., Hanna, M., Banks, E., Sivachenko, A., Cibulskis, K., Kernytsky, A., Garimella, K., Altshuler, D., Gabriel, S., Daly, M., & DePristo, M. A. (2010). The Genome Analysis Toolkit: A MapReduce framework for analyzing next-generation DNA sequencing data. Genome Research, 20(9), 1297–1303. https://doi.org/10.1101/gr.107524.110

67. Meier, J. I., Salazar, P. A., Kučka, M., Davies, R. W., Dréau, A., Aldás, I., Box Power, O., Nadeau, N. J., Bridle, J. R., Rolian, C., Barton, N. H., McMillan, W. O., Jiggins, C. D., & Chan, Y. F. (2021). Haplotype tagging reveals parallel formation of hybrid races in two butterfly species. Proceedings of the National Academy of Sciences of the United States of America, 118(25), e2015005118. https://doi.org/10.1073/pnas.2015005118

68. Melters, D. P., Bradnam, K. R., Young, H. A., Telis, N., May, M. R., Ruby, J. G., Sebra, R., Peluso, P., Eid, J., Rank, D., Garcia, J. F., DeRisi, J. L., Smith, T., Tobias, C., Ross-Ibarra, J., Korf, I., & Chan, S. W. (2013). Comparative analysis of tandem repeats from hundreds of species reveals unique insights into centromere evolution. Genome Biology, 14(1), R10. https://doi.org/10.1186/gb-2013-14-1-r10

69. Mérot, C., Llaurens, V., Normandeau, E., Bernatchez, L., & Wellenreuther, M. (2020). Balancing selection via life-history trade-offs maintains an inversion polymorphism in a seaweed fly. Nature Communications, 11(1), Article 1. https://doi.org/10.1038/s41467-020-14479-7

70. Mérot, C., Oomen, R. A., Tigano, A., & Wellenreuther, M. (2020). A Roadmap for Understanding the Evolutionary Significance of Structural Genomic Variation. Trends in Ecology & Evolution, 35(7), 561–572. https://doi.org/10.1016/j.tree.2020.03.002

71. Morisse, P., Legeai, F., & Lemaitre, C. (2021). LEVIATHAN: Efficient discovery of large structural variants by leveraging long-range information from Linked-Reads data (p. 2021.03.25.437002). bioRxiv. https://doi.org/10.1101/2021.03.25.437002

72. Morisse, P., Lemaitre, C., & Legeai, F. (2021). LRez: A C++ API and toolkit for analyzing and managing Linked-Reads data. Bioinformatics Advances, 1(1), vbab022. https://doi.org/10.1093/bioadv/vbab022

73. Navarro, A., & Barton, N. H. (2003). Accumulating postzygotic isolation genes in parapatry: A new twist on chromosomal speciation. Evolution; International Journal of Organic Evolution, 57(3), 447–459. https://doi.org/10.1111/j.0014-3820.2003.tb01537.x

74. Navarro, A., Betrán, E., Barbadilla, A., & Ruiz, A. (1997). Recombination and Gene Flux Caused by Gene Conversion and Crossing Over in Inversion Heterokaryotypes. Genetics, 146(2), 695–709. https://doi.org/10.1093/genetics/146.2.695

75. Nei, M., Kojima, K.-I., & Schaffer, H. E. (1967). FREQUENCY CHANGES OF NEW INVERSIONS IN POPULATIONS UNDER MUTATION-SELECTION EQUILIBRIA. Genetics, 57(4), 741–750. https://doi.org/10.1093/genetics/57.4.741

76. Noor, M. A. F., Grams, K. L., Bertucci, L. A., & Reiland, J. (2001). Chromosomal inversions and the reproductive isolation of species. Proceedings of the National Academy of Sciences, 98(21), 12084–12088. https://doi.org/10.1073/pnas.221274498

77. Ortiz-Barrientos, D., Engelstädter, J., & Rieseberg, L. H. (2016). Recombination Rate Evolution and the Origin of Species. Trends in Ecology & Evolution, 31(3), 226–236. https://doi.org/10.1016/j.tree.2015.12.016

78. Pedersen, B. S., & Quinlan, A. R. (2017). cyvcf2: Fast, flexible variant analysis with Python. Bioinformatics, 33(12), 1867–1869. https://doi.org/10.1093/bioinformatics/btx057

79. Pettersson, M. E., Rochus, C. M., Han, F., Chen, J., Hill, J., Wallerman, O., Fan, G., Hong, X., Xu, Q., Zhang, H., Liu, S., Liu, X., Haggerty, L., Hunt, T., Martin, F. J., Flicek, P., Bunikis, I., Folkvord, A., & Andersson, L. (2019). A chromosome-level assembly of the Atlantic herring genome-detection of a supergene and other signals of selection. Genome Research, 29(11), 1919–1928. https://doi.org/10.1101/gr.253435.119

80. Picard toolkit. (2019). In Broad Institute, GitHub repository. Broad Institute. https://broadinstitute.github.io/picard/

81. M., Rambla, J., Feder, J. L., Navarro, A., & Faria, R. (2021). Inversions and genomic differentiation after secondary contact: When drift contributes to maintenance, not loss, of differentiation. Evolution, 75(6), 1288–1303. https://doi.org/10.1111/evo.14223

82. Revell, L. J. (2012). phytools: An R package for phylogenetic comparative biology (and other things). Methods in Ecology and Evolution, 3(2), 217–223. https://doi.org/10.1111/j.2041-210X.2011.00169.x

83. Rhie, A., McCarthy, S. A., Fedrigo, O., Damas, J., Formenti, G., Koren, S., Uliano-Silva, M., Chow, W., Fungtammasan, A., Kim, J., Lee, C., Ko, B. J., Chaisson, M., Gedman, G. L., Cantin, L. J., Thibaud-Nissen, F., Haggerty, L., Bista, I., Smith, M.,…Jarvis, E. D. (2021). Towards complete and error-free genome assemblies of all vertebrate species. Nature, 592(7856), Article 7856. https://doi.org/10.1038/s41586-021-03451-0

84. Rieseberg, L. H. (2001). Chromosomal rearrangements and speciation. Trends in Ecology & Evolution, 16(7), 351–358. https://doi.org/10.1016/s0169-5347(01)02187-5

85. Riquet, F., Liautard-Haag, C., Woodall, L., Bouza, C., Louisy, P., Hamer, B., Otero-Ferrer, F., Aublanc, P., Béduneau, V., Briard, O., Ayari, T. E., Hochscheid, S., Belkhir, K., Arnaud-Haond, S., Gagnaire, P.-A., & Bierne, N. (2019). Parallel pattern of differentiation at a genomic island shared between clinal and mosaic hybrid zones in a complex of cryptic seahorse lineages. Evolution, 73(4), 817–835. https://doi.org/10.1111/evo.13696

86. Schaeffer, S. W., & Anderson, W. W. (2005). Mechanisms of Genetic Exchange Within the Chromosomal Inversions of Drosophila pseudoobscura. Genetics, 171(4), 1729–1739. https://doi.org/10.1534/genetics.105.041947

87. Schwander, T., Libbrecht, R., & Keller, L. (2014). Supergenes and Complex Phenotypes. Current Biology, 24(7), R288–R294. https://doi.org/10.1016/j.cub.2014.01.056

88. Shajii, A., Numanagić, I., & Berger, B. (2018). Latent Variable Model for Aligning Barcoded Short-Reads Improves Downstream Analyses. Research in computational molecular biology:…Annual International Conference, RECOMB…: proceedings. RECOMB (Conference: 2005-), 10812, 280-282.

89. Sievert, C., Iannone, R., Allaire, J. J., & Borges, B. (2022). flexdashboard: R Markdown Format for Flexible Dashboards. https://pkgs.rstudio.com/flexdashboard/, https://github.com/rstudio/flexdashboard/.

90. Skoglund, P., Northoff, B. H., Shunkov, M. V., Derevianko, A. P., Pääbo, S., Krause, J., & Jakobsson, M. (2014). Separating endogenous ancient DNA from modern day contamination in a Siberian Neandertal. Proceedings of the National Academy of Sciences, 111(6), 2229–2234. https://doi.org/10.1073/pnas.1318934111

91. Stiller, J., Short, G., Hamilton, H., Saarman, N., Longo, S., Wainwright, P., Rouse, G. W., & Simison, W. B. (2022). Phylogenomic analysis of Syngnathidae reveals novel relationships, origins of endemic diversity and variable diversification rates. BMC Biology, 20(1), 75. https://doi.org/10.1186/s12915-022-01271-w

92. Thompson, M. J., & Jiggins, C. D. (2014). Supergenes and their role in evolution. Heredity, 113(1), Article 1. https://doi.org/10.1038/hdy.2014.20

93. Tigano, A., & Friesen, V. L. (2016). Genomics of local adaptation with gene flow. Molecular Ecology, 25(10), 2144–2164. https://doi.org/10.1111/mec.13606

94. Tigano, A., Jacobs, A., Wilder, A. P., Nand, A., Zhan, Y., Dekker, J., & Therkildsen, N. O. (2021). Chromosome-Level Assembly of the Atlantic Silverside Genome Reveals Extreme Levels of Sequence Diversity and Structural Genetic Variation. Genome Biology and Evolution, 13(6), evab098. https://doi.org/10.1093/gbe/evab098

95. Todesco, M., Owens, G. L., Bercovich, N., Légaré, J.-S., Soudi, S., Burge, D. O., Huang, K., Ostevik, K. L., Drummond, E. B. M., Imerovski, I., Lande, K., Pascual-Robles, M. A., Nanavati, M., Jahani, M., Cheung, W., Staton, S. E., Muños, S., Nielsen, R., Donovan, L. A.,…Rieseberg, L. H. (2020). Massive haplotypes underlie ecotypic differentiation in sunflowers. Nature, 584(7822), Article 7822. https://doi.org/10.1038/s41586-020-2467-6

96. Van der Auwera, G. A., Carneiro, M. O., Hartl, C., Poplin, R., del Angel, G., Levy-Moonshine, A., Jordan, T., Shakir, K., Roazen, D., Thibault, J., Banks, E., Garimella, K. V., Altshuler, D., Gabriel, S., & DePristo, M. A. (2013). From FastQ Data to High-Confidence Variant Calls: The Genome Analysis Toolkit Best Practices Pipeline. Current Protocols in Bioinformatics, 43(1), 11.10.1-11.10.33. https://doi.org/10.1002/0471250953.bi1110s43

97. Wang, Y., & Leung, F. C. C. (2006). Long inverted repeats in eukaryotic genomes: Recombinogenic motifs determine genomic plasticity. FEBS Letters, 580(5), 1277–1284. https://doi.org/10.1016/j.febslet.2006.01.045

98. Weisenfeld, N. I., Kumar, V., Shah, P., Church, D. M., & Jaffe, D. B. (2017). Direct determination of diploid genome sequences. Genome Research, 27(5), 757–767. https://doi.org/10.1101/gr.214874.116

99. Wellenreuther, M., & Bernatchez, L. (2018a). Eco-Evolutionary Genomics of Chromosomal Inversions. Trends in Ecology & Evolution, 33(6), 427–440. https://doi.org/10.1016/j.tree.2018.04.002

100. Wellenreuther, M., & Bernatchez, L. (2018b). Eco-Evolutionary Genomics of Chromosomal Inversions. Trends in Ecology & Evolution, 33(6), 427–440. https://doi.org/10.1016/j.tree.2018.04.002

101. Wellenreuther, M., Mérot, C., Berdan, E., & Bernatchez, L. (2019). Going beyond SNPs: The role of structural genomic variants in adaptive evolution and species diversification. Molecular Ecology, 28(6), 1203–1209. https://doi.org/10.1111/mec.15066

102. Westram, A. M., Faria, R., Johannesson, K., Butlin, R., & Barton, N. (2022). Inversions and parallel evolution. Philosophical Transactions of the Royal Society B: Biological Sciences, 377(1856), 20210203. https://doi.org/10.1098/rstb.2021.0203

103. Westram, A. M., Stankowski, S., Surendranadh, P., & Barton, N. (2022). What is reproductive isolation? Journal of Evolutionary Biology, 35(9), 1143–1164. https://doi.org/10.1111/jeb.14005

104. Yang, Y.-Y., Lin, F.-J., & Chang, H. (2002). Comparison of Recessive Lethal Accumulation in Inversion-bearing and Inversion-free Chromosomes in Drosophila. Zoological Studies.

105. Yeaman, S. (2013). Genomic rearrangements and the evolution of clusters of locally adaptive loci. Proceedings of the National Academy of Sciences of the United States of America, 110(19), E1743–1751. https://doi.org/10.1073/pnas.1219381110

106. Yeaman, S., & Whitlock, M. C. (2011). The genetic architecture of adaptation under migration-selection balance. Evolution; International Journal of Organic Evolution, 65(7), 1897–1911. https://doi.org/10.1111/j.1558-5646.2011.01269.x

107. Zhang, L., Reifová, R., Halenková, Z., & Gompert, Z. (2021). How Important Are Structural Variants for Speciation? Genes, 12(7), 1084. https://doi.org/10.3390/genes12071084

108. Zheng, X., Levine, D., Shen, J., Gogarten, S. M., Laurie, C., & Weir, B. S. (2012). A high-performance computing toolset for relatedness and principal component analysis of SNP data. Bioinformatics (Oxford, England), 28(24), 3326–3328. https://doi.org/10.1093/bioinformatics/bts606

