## Supplementary tables and figures for "“Divergence and gene flow history at two large chromosomal inversions involved in long-snouted seahorse ecotype formation”"

#### SUPPLEMENTARY MATERIALS

### Supplementary File S1

Supplementary tables and figures


- Supplementary Table S1. Samples used in the study (dataset #1).
- Supplementary Fig S1. Number of reads per barcode (10X Chromium linked-read sequencing).
- Supplementary Fig S2. PMDtools
- Supplementary Fig S3. Snailplot our reference genome.
- Supplementary Fig S4. Snailplots of other genomes used.
- Supplementary Fig S5: D-genies alignment our reference genome vs. H. erectus.
- Supplementary Fig S6: repeats
- Supplementary Fig S7: Tan9
- Supplementary Fig S8: Fst distributions.
- Supplementary Fig S9: Dxy distributions.
- Supplementary Fig S10: Pi distributions.
- Supplementary Fig S11: D-genies alignment of Hgutt_V1 vs. HguttRefA
- Supplementary Fig S12. Haplotagging
- Supplementary Fig S13. Fst co-plot
- Supplementary Fig S14: all painted chromosomes (Chr12)
- Supplementary Fig S15a and S15b: all painted chromosomes (Chr2)
- Supplementary Fig S16: SNPs Riquet mapped scaffold 14

**Supplementary Table S1:** Samples used in the study (dataset #1). For our sequenced samples, codes are composed of the species name abbreviation, the approximate sampling locality, approximate age and the sample number. Dataset #2 consisted of samples for which variant calling was performed (last column). Br: Brest. Ga: Lac Marin de Port d'Albret. Fa: Faro. Mu: Murcia. To: Tossa de mar. By: Banyuls. Va: Valras. Ag: Agde. Ms: Marseillan. Th: Thau lagoon. Se: Sète. Fr: Frontignan. Hy: Hyères. Ma: unknown Mediterranean marine site. Al: unknown Algerian site. Bz: Bizerte lagoon. Tu: unknown Tunisian site. Bs: Varna. Ru: unknown Russian site. Xx: unknown location.

| Sample | Species | Location | Habitat | Type | Extraction | Library | Variant Calling |
|---|---|---|---|---|---|---|---|
| Hgutt_Ru_1856_77 | H. guttulatus | Ru | unknown | Museum | Phenol-chloroform | OVATION | No |
| Hgutt_Al_1864_80 | H. guttulatus | Al | marine | Museum | Phenol-chloroform | OVATION | No |
| Hgutt_Se_1898_81 | H. guttulatus | Se | unknown | Museum | Phenol-chloroform | OVATION | No |
| Hgutt_Se_1898_94 | H. guttulatus | Se | unknown | Museum | Phenol-chloroform | OVATION | No |
| Hgutt_Ag_1935_17 | H. guttulatus | Ag | unknown | Dried | CTAB | OVATION | No |
| Hgutt_Ag_1935_19 | H. guttulatus | Ag | unknown | Dried | CTAB | OVATION | No |
| Hgutt_Ag_1935_59 | H. guttulatus | Ag | unknown | Dried | CTAB | OVATION | No |
| Hgutt_Ag_1935_70 | H. guttulatus | Ag | unknown | Dried | CTAB | OVATION | No |
| Hgutt_Th_1935_66 | H. guttulatus | Th | lagoon | Dried | CTAB | OVATION | No |
| Hgutt_Va_1935_22 | H. guttulatus | Va | marine | Dried | CTAB | OVATION | No |
| Hgutt_Ba_1950_41 | H. guttulatus | Th | lagoon | Dried | CTAB | OVATION | No |
| Hgutt_Ma_1960_60 | H. guttulatus | Ma | marine | Dried | CTAB | OVATION | No |
| Hgutt_Ms_1960_21 | H. guttulatus | Ms | unknown | Dried | CTAB | OVATION | No |
| Hgutt_Ro_1960_23 | H. guttulatus | Th | lagoon | Dried | CTAB | OVATION | No |
| Hgutt_Ro_1960_43 | H. guttulatus | Th | lagoon | Dried | CTAB | OVATION | No |
| Hgutt_Th_1960_11 | H. guttulatus | Th | lagoon | Dried | CTAB | OVATION | No |
| Hgutt_Th_1960_24 | H. guttulatus | Th | lagoon | Dried | CTAB | OVATION | No |
| Hgutt_Th_1960_42 | H. guttulatus | Th | lagoon | Dried | CTAB | OVATION | No |
| Hgutt_Th_1960_44 | H. guttulatus | Th | lagoon | Dried | CTAB | OVATION | No |
| Hgutt_Th_1960_95 | H. guttulatus | Th | lagoon | Dried | CTAB | OVATION | No |
| Hgutt_Th_1964_45 | H. guttulatus | Th | lagoon | Dried | CTAB | OVATION | No |
| Hgutt_Th_1964_ | H. guttulatus | Th | lagoon | Dried | CTAB | OVATION | No |

| | | | | | | | |
|---|---|---|---|---|---|---|---|
| 65 | | | | | | | |
| Hgutt_Th_1970_30 | H. guttulatus | Th | lagoon | Dried | CTAB | OVATION | No |
| Hgutt_Th_1970_46 | H. guttulatus | Th | lagoon | Dried | CTAB | OVATION | No |
| Hgutt_Il_1975_47 | H. guttulatus | Th | lagoon | Dried | CTAB | OVATION | No |
| Hgutt_Il_1975_48 | H. guttulatus | Th | lagoon | Dried | CTAB | OVATION | Yes |
| Hgutt_Il_1975_68 | H. guttulatus | Th | lagoon | Dried | CTAB | OVATION | Yes |
| Hgutt_Bo_1980_50 | H. guttulatus | Th | lagoon | Dried | CTAB | OVATION | No |
| Hgutt_By_1980_67 | H. guttulatus | By | marine | Dried | CTAB | OVATION | No |
| Hgutt_Th_1980_12 | H. guttulatus | Th | lagoon | Dried | CTAB | OVATION | No |
| Hgutt_Th_1980_49 | H. guttulatus | Th | lagoon | Dried | CTAB | OVATION | No |
| Hgutt_Th_1980_69 | H. guttulatus | Th | lagoon | Dried | CTAB | OVATION | No |
| Hgutt_Tu_1980_25 | H. guttulatus | Tu | lagoon | Dried | CTAB | OVATION | No |
| Hgutt_Tu_1980_31 | H. guttulatus | Tu | lagoon | Dried | CTAB | OVATION | No |
| Hgutt_Me_1982_51 | H. guttulatus | Th | lagoon | Dried | CTAB | OVATION | No |
| Hgutt_Fr_1985_52 | H. guttulatus | Fr | marine | Dried | CTAB | OVATION | No |
| Hgutt_Ma_1985_61 | H. guttulatus | Ma | marine | Dried | CTAB | OVATION | No |
| Hgutt_Ms_1985_18 | H. guttulatus | Ms | unknown | Dried | CTAB | OVATION | No |
| Hgutt_Xx_1985_71 | H. guttulatus | Xx | unknown | Dried | CTAB | OVATION | No |
| Hgutt_Ma_1990_63 | H. guttulatus | Ma | marine | Dried | CTAB | OVATION | No |
| Hgutt_Th_1990_72 | H. guttulatus | Th | lagoon | Dried | CTAB | OVATION | Yes |
| Hgutt_Xx_1990_13 | H. guttulatus | Xx | unknown | Dried | CTAB | OVATION | No |
| Hgutt_Xx_1990_14 | H. guttulatus | Xx | unknown | Dried | CTAB | OVATION | No |
| Hgutt_Xx_1990_15 | H. guttulatus | Xx | unknown | Dried | CTAB | OVATION | No |
| Hgutt_Xx_1990_16 | H. guttulatus | Xx | unknown | Dried | CTAB | OVATION | No |
| Hgutt_Xx_1990_26 | H. guttulatus | Xx | unknown | Dried | CTAB | OVATION | No |
| Hgutt_Xx_1990_27 | H. guttulatus | Xx | unknown | Dried | CTAB | OVATION | No |
| Hgutt_Xx_1990_32 | H. guttulatus | Xx | unknown | Dried | CTAB | OVATION | No |
| Hgutt_Xx_1990_33 | H. guttulatus | Xx | unknown | Dried | CTAB | OVATION | No |

| | | | | | | | |
|---|---|---|---|---|---|---|---|
| Hgutt_Xx_1990_34 | H. guttulatus | Xx | unknown | Dried | CTAB | OVATION | No |
| Hgutt_Xx_1990_53 | H. guttulatus | Xx | unknown | Dried | CTAB | OVATION | No |
| Hgutt_Xx_1990_54 | H. guttulatus | Xx | unknown | Dried | CTAB | OVATION | No |
| Hgutt_Th_1991_55 | H. guttulatus | Th | lagoon | Dried | CTAB | OVATION | No |
| Hgutt_Ms_1995_64 | H. guttulatus | Ms | unknown | Dried | CTAB | OVATION | No |
| Hgutt_Th_1995_56 | H. guttulatus | Th | lagoon | Dried | CTAB | OVATION | No |
| Hgutt_Th_1995_76 | H. guttulatus | Th | lagoon | Dried | CTAB | OVATION | No |
| Hgutt_Xx_1995_20 | H. guttulatus | Xx | unknown | Dried | CTAB | OVATION | Yes |
| Hgutt_Xx_1995_38 | H. guttulatus | Xx | unknown | Dried | CTAB | OVATION | No |
| Hgutt_Xx_1995_39 | H. guttulatus | Xx | unknown | Dried | CTAB | OVATION | No |
| Hgutt_Xx_1995_73 | H. guttulatus | Xx | unknown | Dried | CTAB | OVATION | Yes |
| Hgutt_Th_1996_28 | H. guttulatus | Th | lagoon | Dried | CTAB | OVATION | No |
| Hgutt_Th_1996_74 | H. guttulatus | Th | lagoon | Dried | CTAB | OVATION | No |
| Hgutt_Th_1996_75 | H. guttulatus | Th | lagoon | Dried | CTAB | OVATION | No |
| Hgutt_Bs_2006_83 | H. guttulatus | Bs | unknown | Fresh | Nucleospin | OVATION | Yes |
| Hgutt_Bs_2006_85 | H. guttulatus | Bs | unknown | Fresh | Nucleospin | OVATION | Yes |
| Hgutt_Bs_2006_87 | H. guttulatus | Bs | unknown | Fresh | Nucleospin | OVATION | Yes |
| Hgutt_Bs_2006_89 | H. guttulatus | Bs | unknown | Fresh | Nucleospin | OVATION | Yes |
| Hgutt_Bs_2006_91 | H. guttulatus | Bs | unknown | Fresh | Nucleospin | OVATION | Yes |
| Hgutt_Th_2007_36 | H. guttulatus | Th | lagoon | Dried | CTAB | OVATION | Yes |
| Hgutt_Th_2007_57 | H. guttulatus | Th | lagoon | Dried | CTAB | OVATION | No |
| Hgutt_Me_2010_58 | H. guttulatus | Th | lagoon | Dried | CTAB | OVATION | Yes |
| Hgutt_Br_2014_06 | H. guttulatus | Br | marine | Fresh | Nucleospin | OVATION | Yes |
| Hgutt_Br_2014_07 | H. guttulatus | Br | marine | Fresh | Nucleospin | OVATION | Yes |
| Hgutt_Br_2014_08 | H. guttulatus | Br | marine | Fresh | Nucleospin | OVATION | Yes |
| Hgutt_Br_2014_09 | H. guttulatus | Br | marine | Fresh | Nucleospin | OVATION | Yes |
| Hgutt_Br_2014_10 | H. guttulatus | Br | marine | Fresh | Nucleospin | OVATION | Yes |
| Hgutt_Bz_2014_ | H. guttulatus | Bz | lagoon | Fresh | Nucleospin | OVATION | Yes |

| | | | | | | | |
|---|---|---|---|---|---|---|---|
| 82 | | | | | | | |
| Hgutt_Bz_2014_84 | H. guttulatus | Bz | lagoon | Fresh | Nucleospin | OVATION | Yes |
| Hgutt_Bz_2014_86 | H. guttulatus | Bz | lagoon | Fresh | Nucleospin | OVATION | Yes |
| Hgutt_Bz_2014_88 | H. guttulatus | Bz | lagoon | Fresh | Nucleospin | OVATION | Yes |
| Hgutt_Bz_2014_90 | H. guttulatus | Bz | lagoon | Fresh | Nucleospin | OVATION | Yes |
| Hgutt_To_2016_01 | H. guttulatus | To | marine | Fresh | Nucleospin | OVATION | Yes |
| Hgutt_To_2016_02 | H. guttulatus | To | marine | Fresh | Nucleospin | OVATION | Yes |
| Hgutt_Hy_2017_03 | H. guttulatus | Hy | marine | Fresh | Nucleospin | OVATION | Yes |
| Hgutt_Hy_2017_04 | H. guttulatus | Hy | marine | Fresh | Nucleospin | OVATION | Yes |
| Hgutt_Hy_2017_05 | H. guttulatus | Hy | marine | Fresh | Nucleospin | OVATION | Yes |
| HguttLi1 | H. guttulatus | Th | lagoon | Barry et al. (2022) | Nucleospin | TruSeq | Yes |
| HguttLi3 | H. guttulatus | Th | lagoon | Barry et al. (2022) | Nucleospin | TruSeq | Yes |
| HguttLi4 | H. guttulatus | Th | lagoon | Barry et al. (2022) | Nucleospin | TruSeq | Yes |
| HguttLi5 | H. guttulatus | Th | lagoon | Barry et al. (2022) | Nucleospin | TruSeq | Yes |
| HguttLi6 | H. guttulatus | Th | lagoon | Barry et al. (2022) | Nucleospin | TruSeq | Yes |
| HguttGa10 | H. guttulatus | Ga | lagoon | Barry et al. (2022) | Nucleospin | TruSeq | No |
| HguttGa11 | H. guttulatus | Ga | lagoon | Barry et al. (2022) | Nucleospin | TruSeq | Yes |
| HguttGa12 | H. guttulatus | Ga | lagoon | Barry et al. (2022) | Nucleospin | TruSeq | No |
| HguttGa1 | H. guttulatus | Ga | lagoon | Barry et al. (2022) | Nucleospin | TruSeq | No |
| HguttGa2 | H. guttulatus | Ga | lagoon | Barry et al. (2022) | Nucleospin | TruSeq | No |
| HguttGa3 | H. guttulatus | Ga | lagoon | Barry et al. (2022) | Nucleospin | TruSeq | Yes |
| HguttGa4 | H. guttulatus | Ga | lagoon | Barry et al. (2022) | Nucleospin | TruSeq | No |
| HguttGa6 | H. guttulatus | Ga | lagoon | Barry et al. (2022) | Nucleospin | TruSeq | Yes |
| HguttGa7 | H. guttulatus | Ga | lagoon | Barry et al. (2022) | Nucleospin | TruSeq | Yes |
| HguttGa8 | H. guttulatus | Ga | lagoon | Barry et al. (2022) | Nucleospin | TruSeq | Yes |
| HguttGa9 | H. guttulatus | Ga | lagoon | Barry et al. (2022) | Nucleospin | TruSeq | Yes |
| HguttMu1 | H. guttulatus | Mu | lagoon | Barry et al. (2022) | Nucleospin | TruSeq | Yes |
| HguttMu2 | H. guttulatus | Mu | lagoon | Barry et al. (2022) | Nucleospin | TruSeq | Yes |

| | | | | | | | |
|---|---|---|---|---|---|---|---|
| HguttMu3 | H. guttulatus | Mu | lagoon | Barry et al. (2022) | Nucleospin | TruSeq | Yes |
| HguttMu4 | H. guttulatus | Mu | lagoon | Barry et al. (2022) | Nucleospin | TruSeq | Yes |
| HguttMu5 | H. guttulatus | Mu | lagoon | Barry et al. (2022) | Nucleospin | TruSeq | Yes |
| HguttFa2 | H. guttulatus | Fa | lagoon | Barry et al. (2022) | Nucleospin | TruSeq | Yes |
| HguttFa3 | H. guttulatus | Fa | lagoon | Barry et al. (2022) | Nucleospin | TruSeq | Yes |
| HguttFa4 | H. guttulatus | Fa | lagoon | Barry et al. (2022) | Nucleospin | TruSeq | Yes |
| HguttFa5 | H. guttulatus | Fa | lagoon | Barry et al. (2022) | Nucleospin | TruSeq | Yes |
| HguttFa6 | H. guttulatus | Fa | lagoon | Barry et al. (2022) | Nucleospin | TruSeq | Yes |

**Supplementary Figure S1:** Count distribution of the number of read pairs per barcode in the deduplicated 10X Chromium linked-read sequencing data generated for the reference genome assembly (Hgutt_V1). Molecular barcodes associated with low numbers of read pairs (left part of the distribution shaded in red) reflect noise in the 10X data (e.g. sequencing errors within barcodes), and were therefore excluded before performing *de novo* assembly (1,649,606 read pairs, 1.1% of the total sequencing data). Over-represented molecular barcodes on the right end of the distribution (28,345 read pairs) were also excluded. The resulting average number of read pairs per retained barcode was approximately 100, with a maximum occurrence at 72.

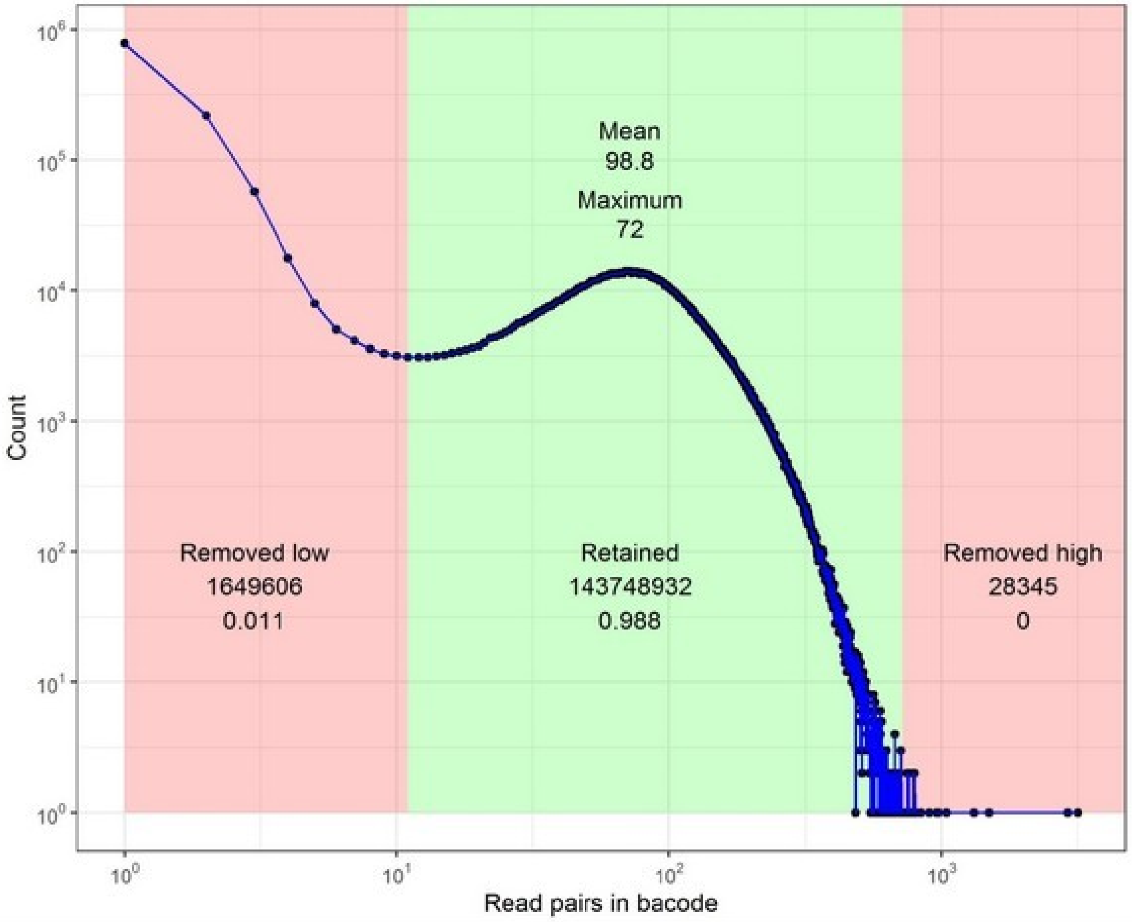

**Supplementary Figure S2:** DNA damage patterns (cytosine deamination) visualised using PMDtools (v0.60) (Skoglund et al., 2014). Results are shown for two historical samples.

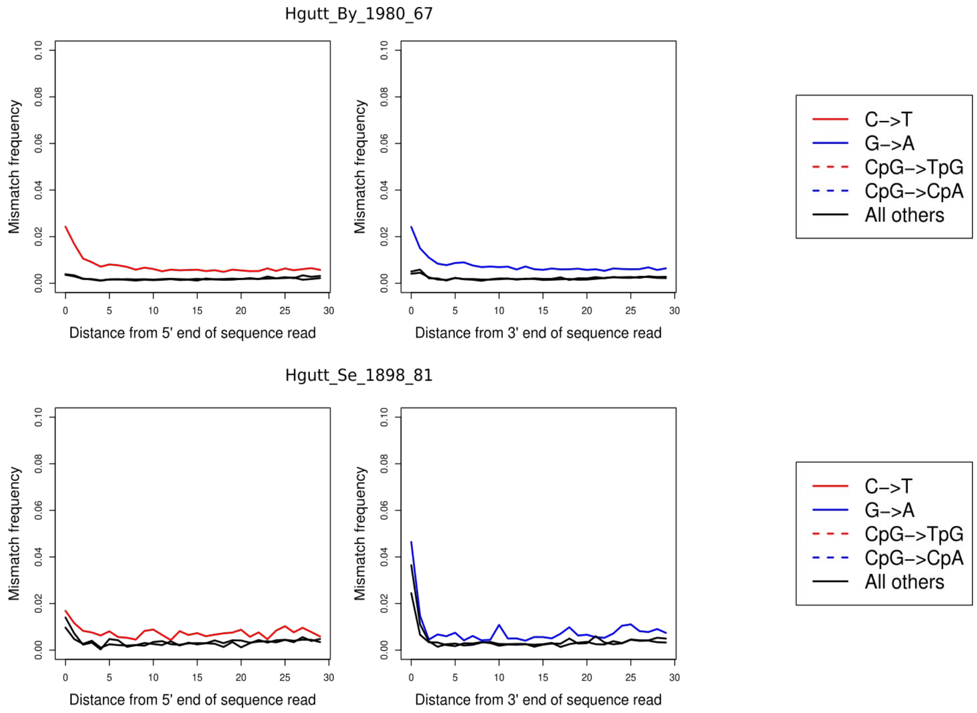

**Supplementary Figure S3:** Snail plot describing the assembly statistics of the *Hippocampus guttulatus* Hgutt_V1 reference genome. The genome contiguity is shown in a circle representing the full assembly length of 451 Mbp, with the N50 length in dark orange and the N90 length in light orange. The longest scaffold was 28.8 Mbp. The completeness BUSCO scores (in shades of green) and base composition (percentage of GC in dark blue, AT in light blue, and N in light grey) appear in top right and bottom left panels, respectively. The plot was generated using *Blobtoolkit* (https://blobtoolkit.genomehubs.org/).

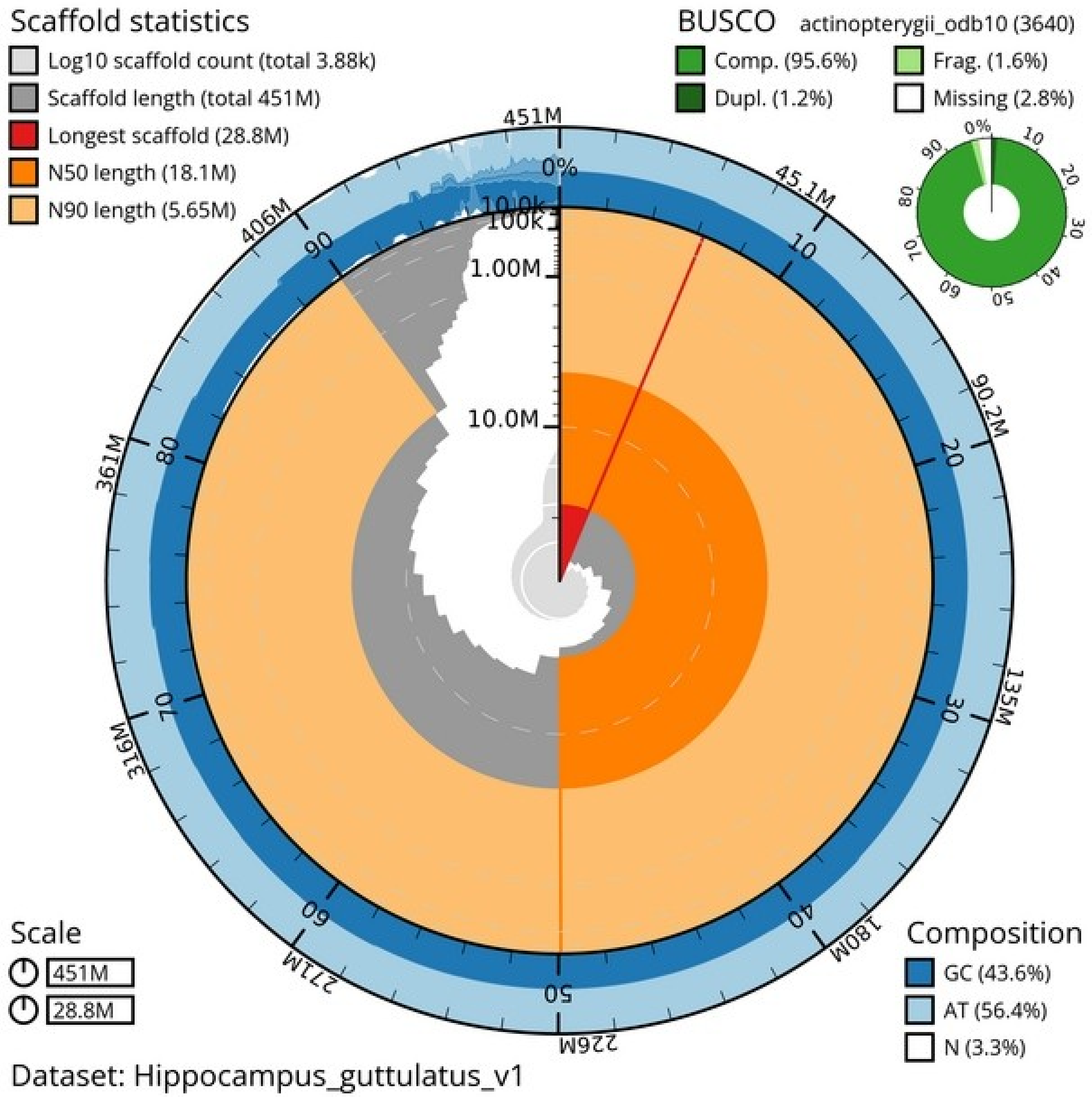

**Supplementary Figure S4:** Snail plots describing the assembly statistics of the *Hippocampus guttulatus* GCA_025802095.1 reference genome (HguttRefA, Jones et al in prep) and the *Hippocampus erectus* reference genome (Li et al 2021).

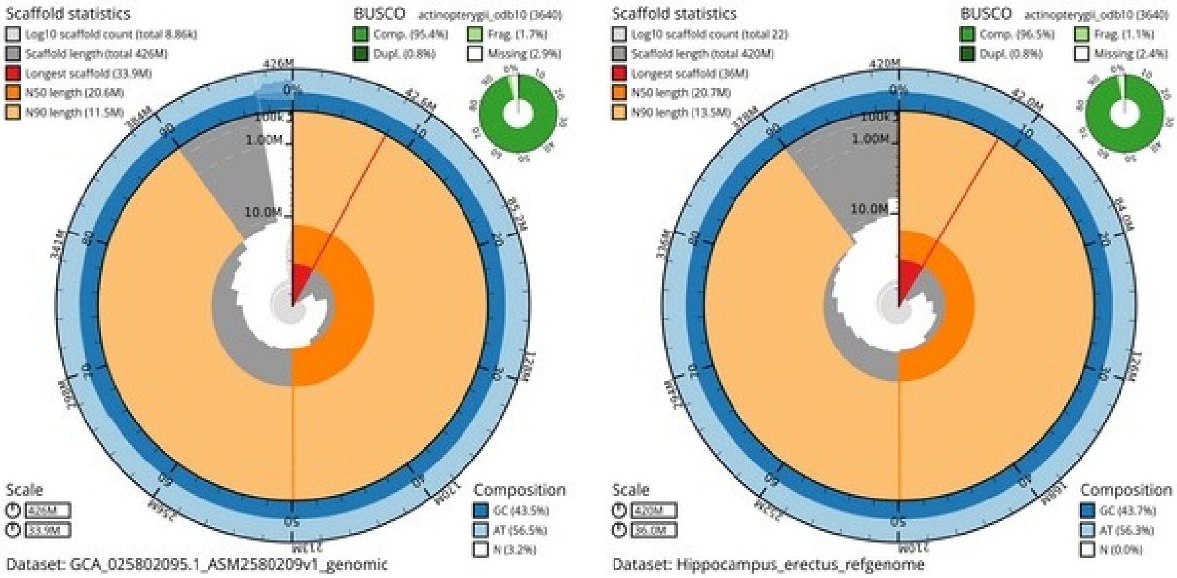

**Supplementary Figure S5:** Genomic alignment dot plot showing the comparison between the chromosome-level assembly of *Hippocampus erectus* (top) and our *H. guttulatus* Hgutt_V1 genome assembly (right). Only shown here are alignment matches passing a minimum size filter and a similarity threshold of 50%. All *H. guttulatus* scaffolds match a single *H. erectus* chromosome (with varied number of intrachromosomal rearrangements), except for scaffold 7 that matches both Chr10 and Chr14 of the *H. erectus* genome. The plot was generated using *D-GENIES* ([https://github.com/genotoul-bioinfo/dgenies](https://github.com/genotoul-bioinfo/dgenies)).

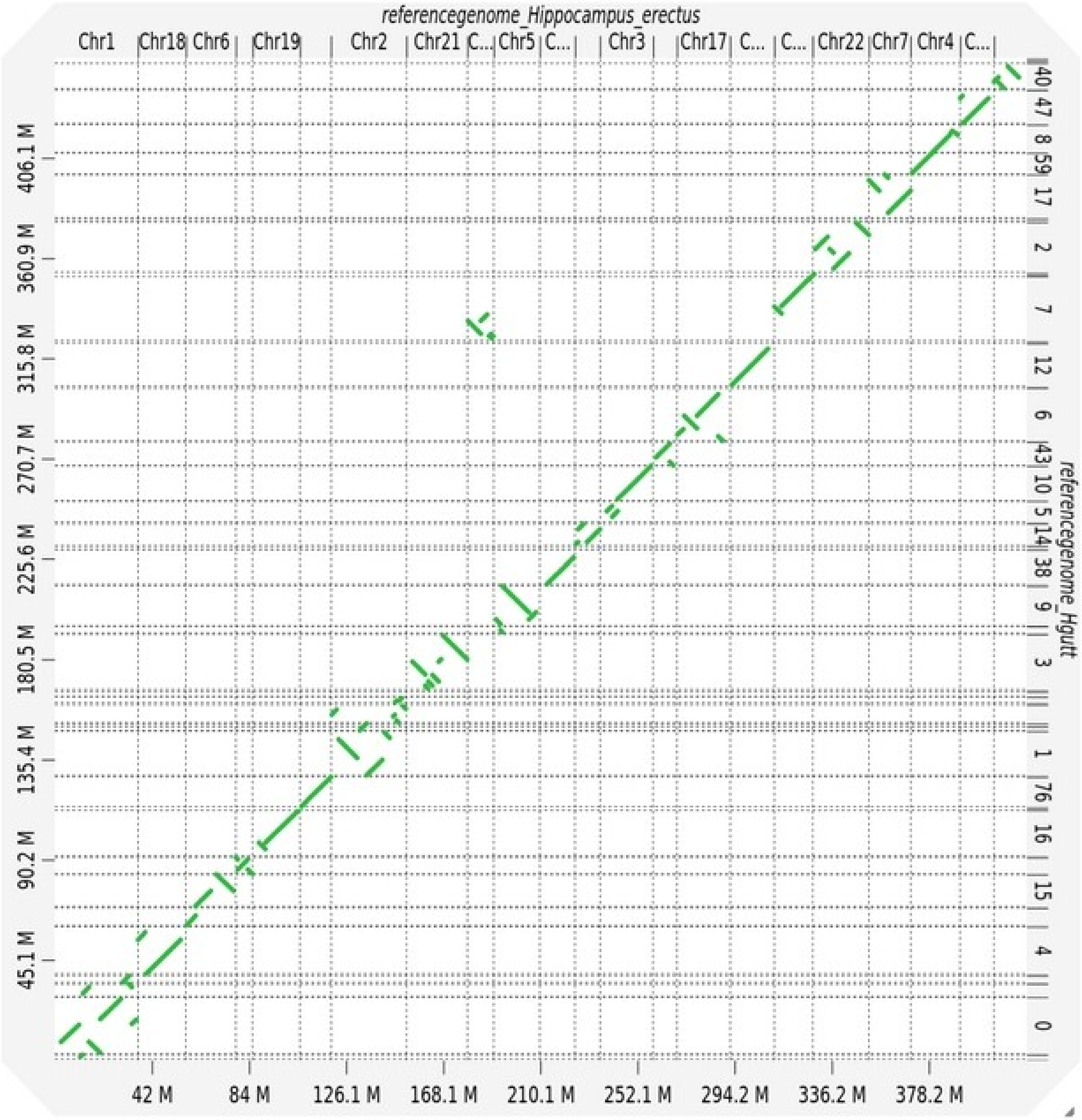

**Supplementary Figure S6:** Repeat landscape showing the percentage of the Hgutt_V1 genome occupied by different categories of interspersed repeat elements (right), and detailed as a function of weighted average Kimura divergence in alignments for each repeat family (left).

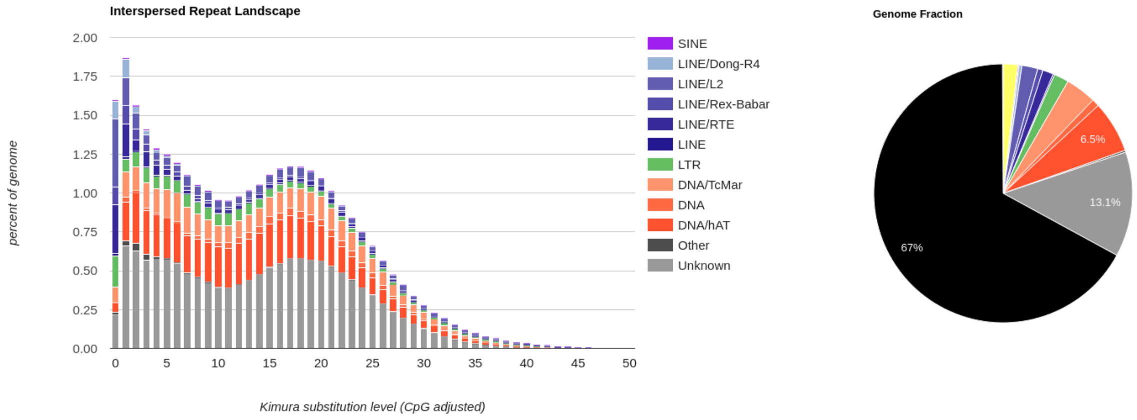

**Supplementary Figure S7:** Repeat landscape showing the number of Tan9 monomer repeats within 10kb windows across Hgutt_V1 scaffolds longer than 1Mb. The consensus sequence of the Tan9 monomer (37bp, GTCGTTTTTTTCGGCTAAAAACGCCTTACTATACATG) was searched with *blastn* with parameters set to M=1, N=-1, Q=2, R=2, W=8, following (Melters et al 2013). Red boxes indicate the Tan9 repeats where the two inversion breakpoints map on Scaffold 14 (Chr12), delimiting an 8.2 Mb-long inversion.

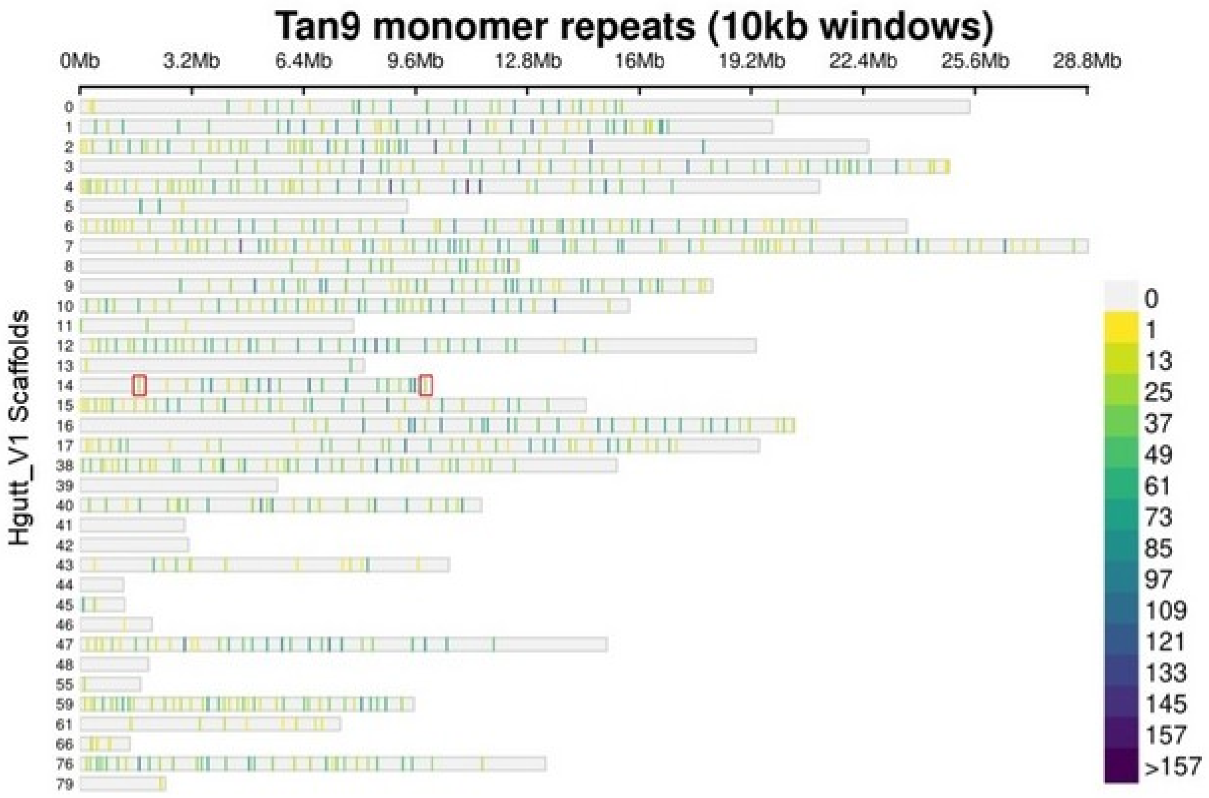

**Supplementary Figure S8:** Distributions of $F_{ST}$ values displayed separately for the genome background, Chr2 and Chr12. Black text gives information about the populations being compared: oceanic basins (“Background” column), total number of C/D alleles in the comparison (“Chr2” column), or total number of A/B alleles in the comparison (“Chr12” column). $F_{ST}$ was calculated for each population pair using the popgenWindows.py script in 25 kb windows (Martin, 2018; https://github.com/simonhmartin/genomics_general). The “Ma” population is here represented by individuals from the Mediterranean marine sites Tossa de mar and Hyères.

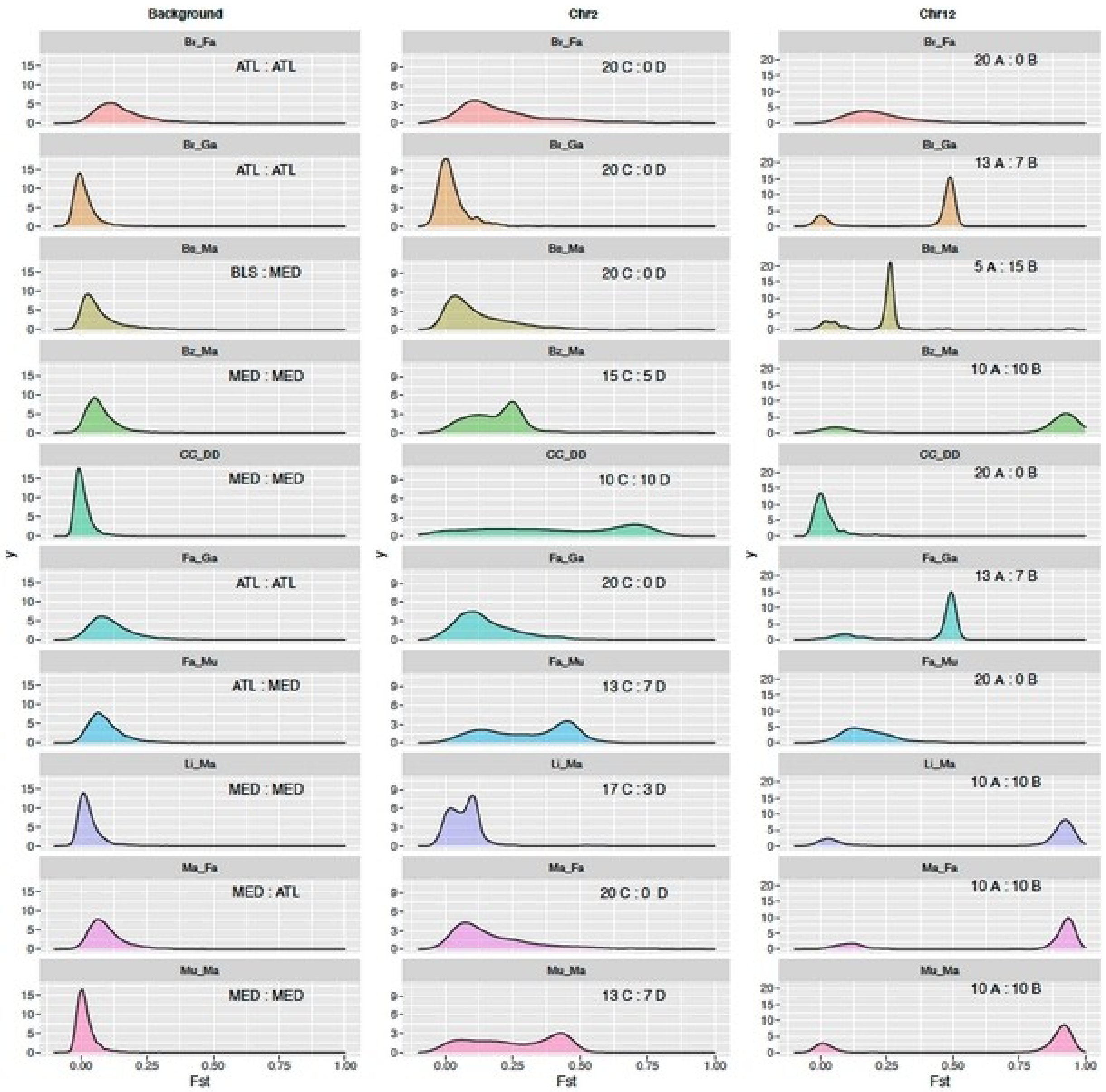

**Supplementary Figure S9:** Distributions of $D_{XY}$ values displayed separately for the genome background, Chr2 and Chr12 (for details see previous figure).

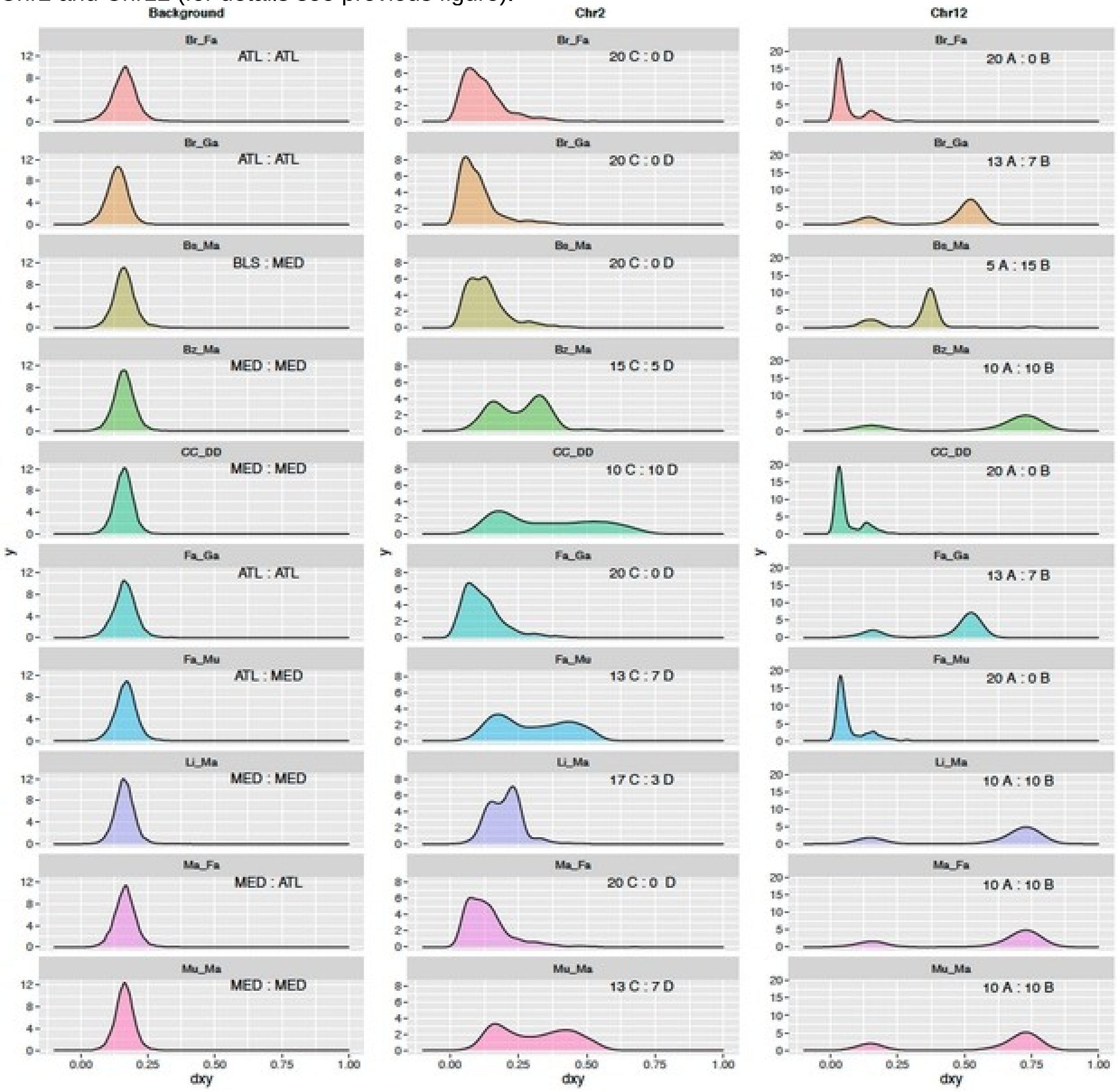

**Supplementary Figure S10:** Distributions of nucleotide diversity ($\pi$) values displayed separately for the genome background, Chr2 and Chr12. Nucleotide diversity was calculated for each population or haplotype group (for details see Supplementary Fig. S8).

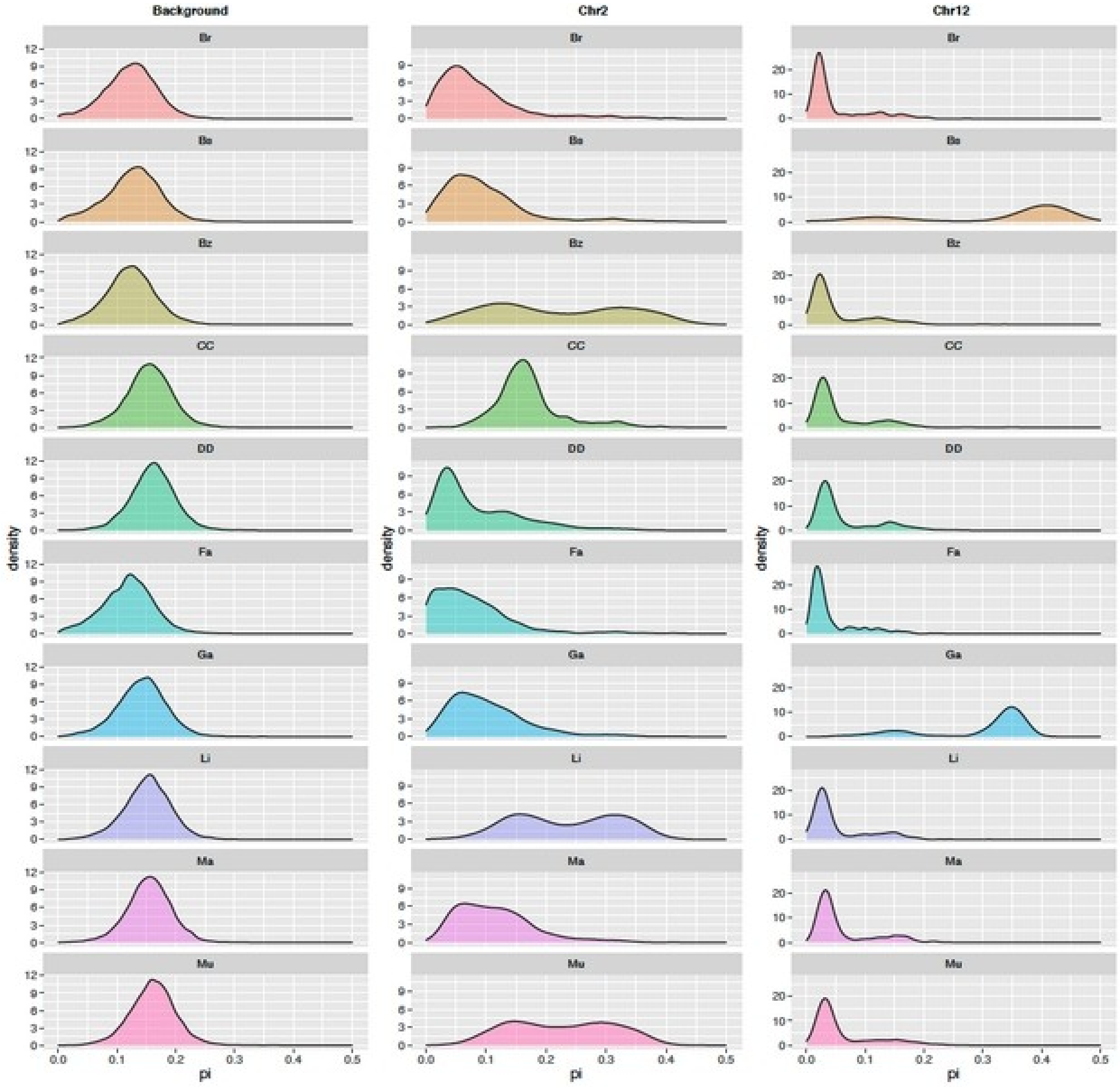

**Supplementary Figure S11:** Genomic alignment dot plot showing the comparison between the chromosome-level assembly of *Hippocampus guttulatus* from the English Channel (HguttRefA) (top) and our Hgutt_V1 genome assembly (right). Only shown here are alignment matches passing a minimum size filter and a similarity threshold of 50%. The zoomed section shows chromosome JAOYMQ010000004.1 which corresponds to *H. erectus* Chr11 (scaffold 61) and Chr12. The junction between Chr11 and Chr12 within chromosome JAOYMQ010000004.1 is possibly an artefact of the assembly pipeline (i.e. the junction is not supported by reads from *H. guttulatus*). The purple asterisk indicates the 8.2 Mb-long region that is inverted between the two *H. guttulatus* reference assemblies. The plot was generated using *D-GENIES* (https://github.com/genotoul-bioinfo/dgenies).

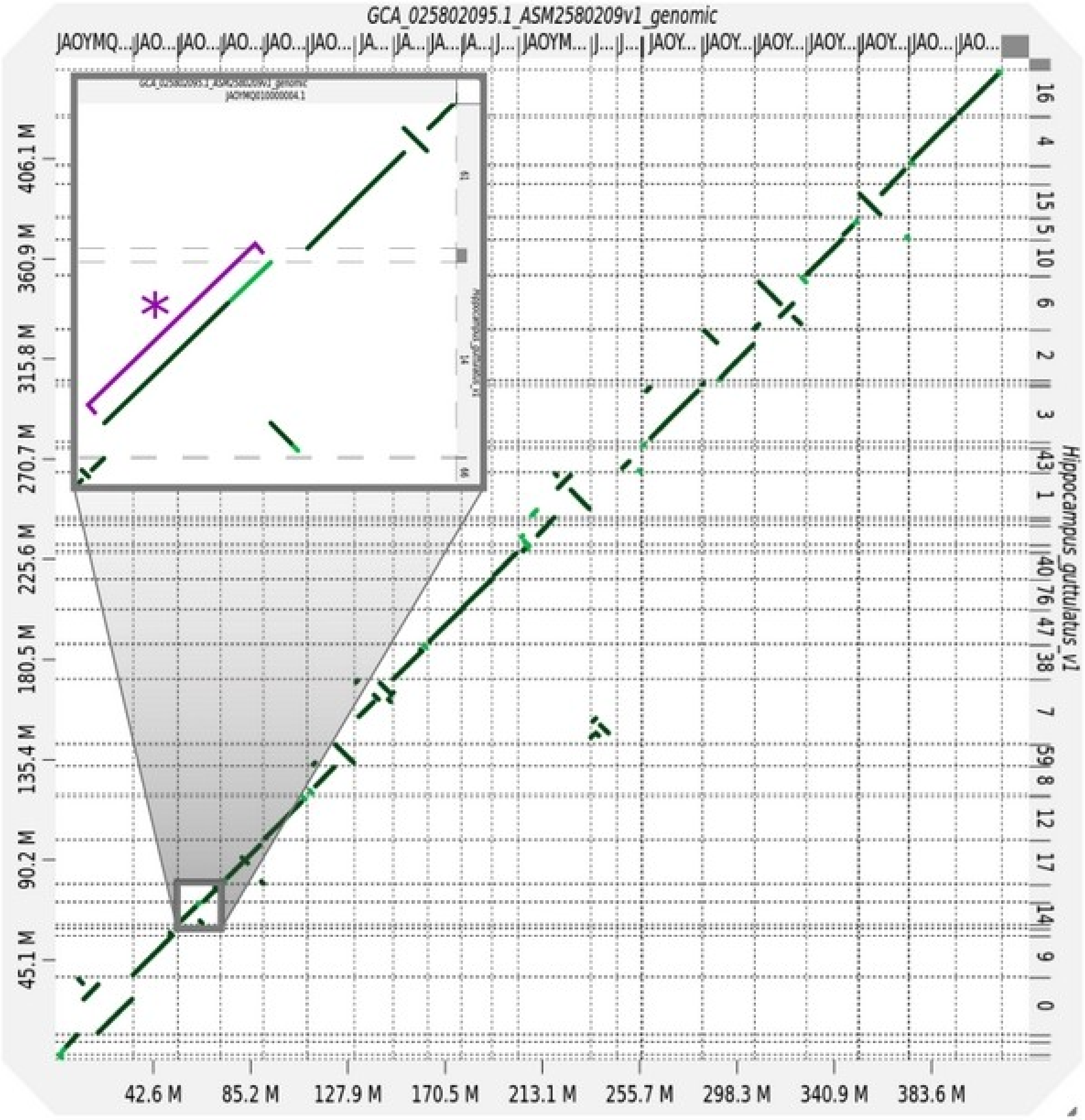

**Supplementary Figure S12:** Direct identification of an 8.2 Mb inversion on Chr12 and characterisation of inversion breakpoints. *Left panel:* The region around the inversion breakpoint localised near position 1,699 Mb (read dotted line) on scaffold 14 of Hgutt_V1 (Blue horizontal line, B arrangement) was reassembled using *MTG-Link*, resulting in a 3kb contig matching the reference (blue rectangle). A long inverted repeat of Tan9 monomers (blue triangles) was found centred at the breakpoint position. Linked-read sequencing data obtained for three individuals with alternate B12 genotypes (i.e. the BB individual used for reference genome assembly sequenced at >100X coverage, plus an AA and an AB obtained by haplotagging and sequenced at ~12X coverage) were mapped against the Hgutt_V1 genome assembly. Horizontal links in each local alignment show linked paired-end reads sharing the same molecular barcode. Long synthetic molecules spanning the 3kb breakpoint region were found for both the BB and the AB genotypes, but not for the AA genotype that showed a ~1.5kb mapping gap directly after the breakpoint. *Right panel:* Linked-reads from the BB genotype were mapped against the HguttRefA assembly (Red horizontal line, A arrangement), and analysed with *Leviathan* to search for structural variants. A total of 63 long synthetic molecules (coloured segments) were found shared by two regions spanning the inversion breakpoints and separated by 8.2 Mb. Read orientation supported a structural variant of type inversion.

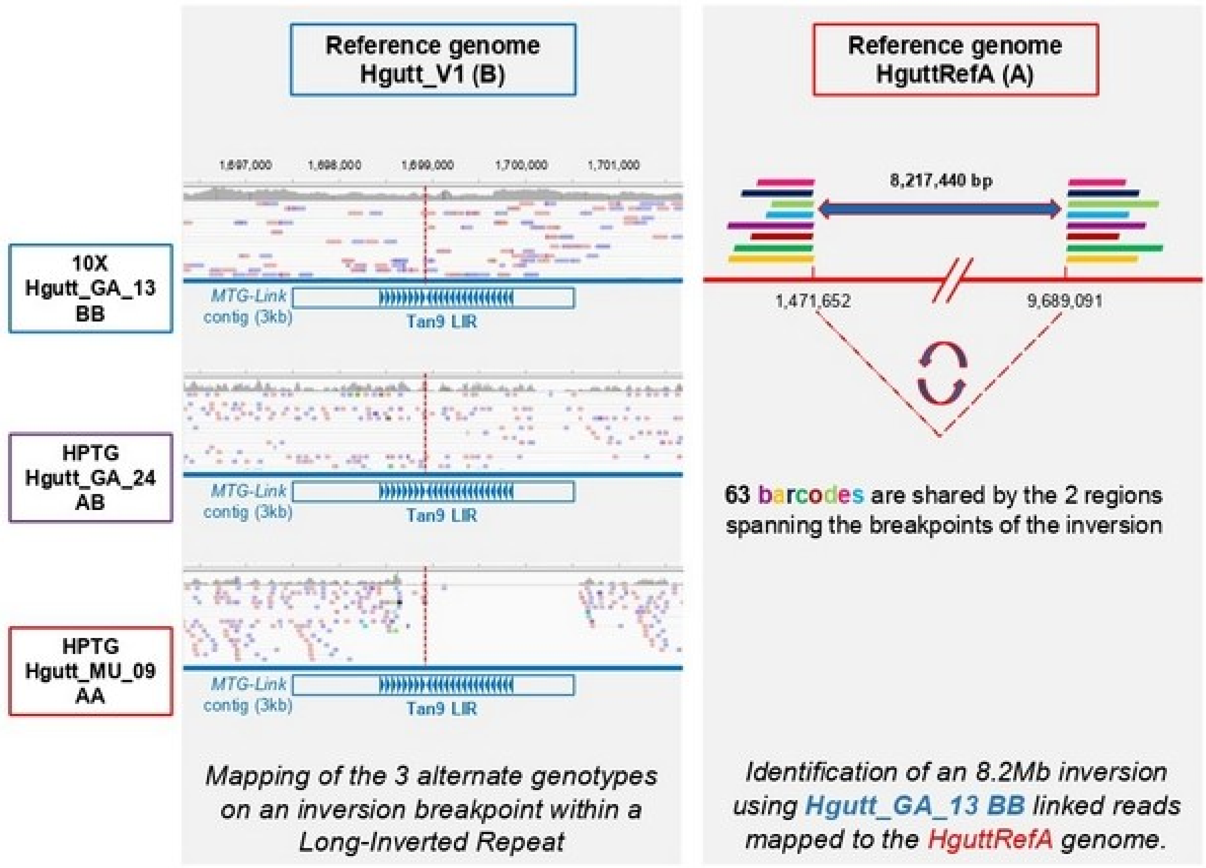

**Supplementary Figure S13:** $F_{ST}$ co-plots between different Mediterranean and Atlantic lineages.

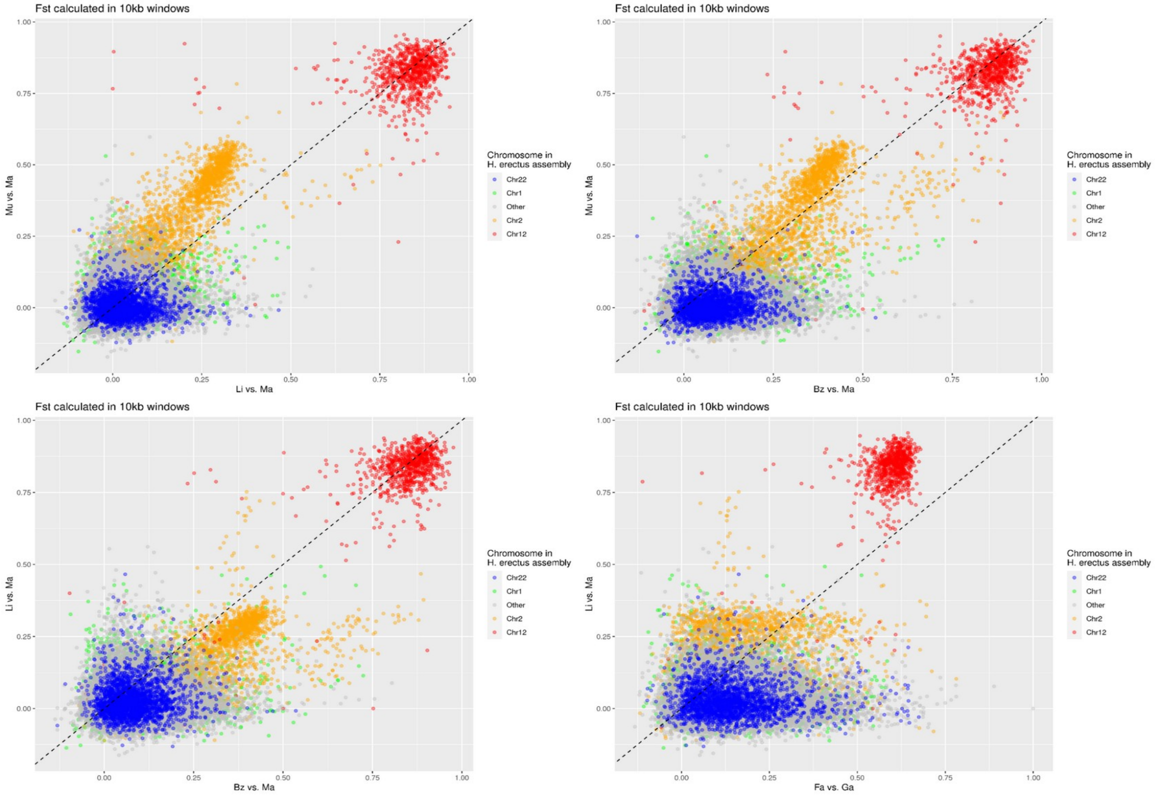

**Supplementary Figure S14:** Chromosome painting in the inversion region on Chr12. Codes indicate inversion genotypes and individual sample names of which the last characters ("-1" or "-2") refer to phased parental haplotypes. Inversion heterokaryotes show phasing errors in the form of haplotype switches. The top panels show the first principal component of local PCA along the inversion.

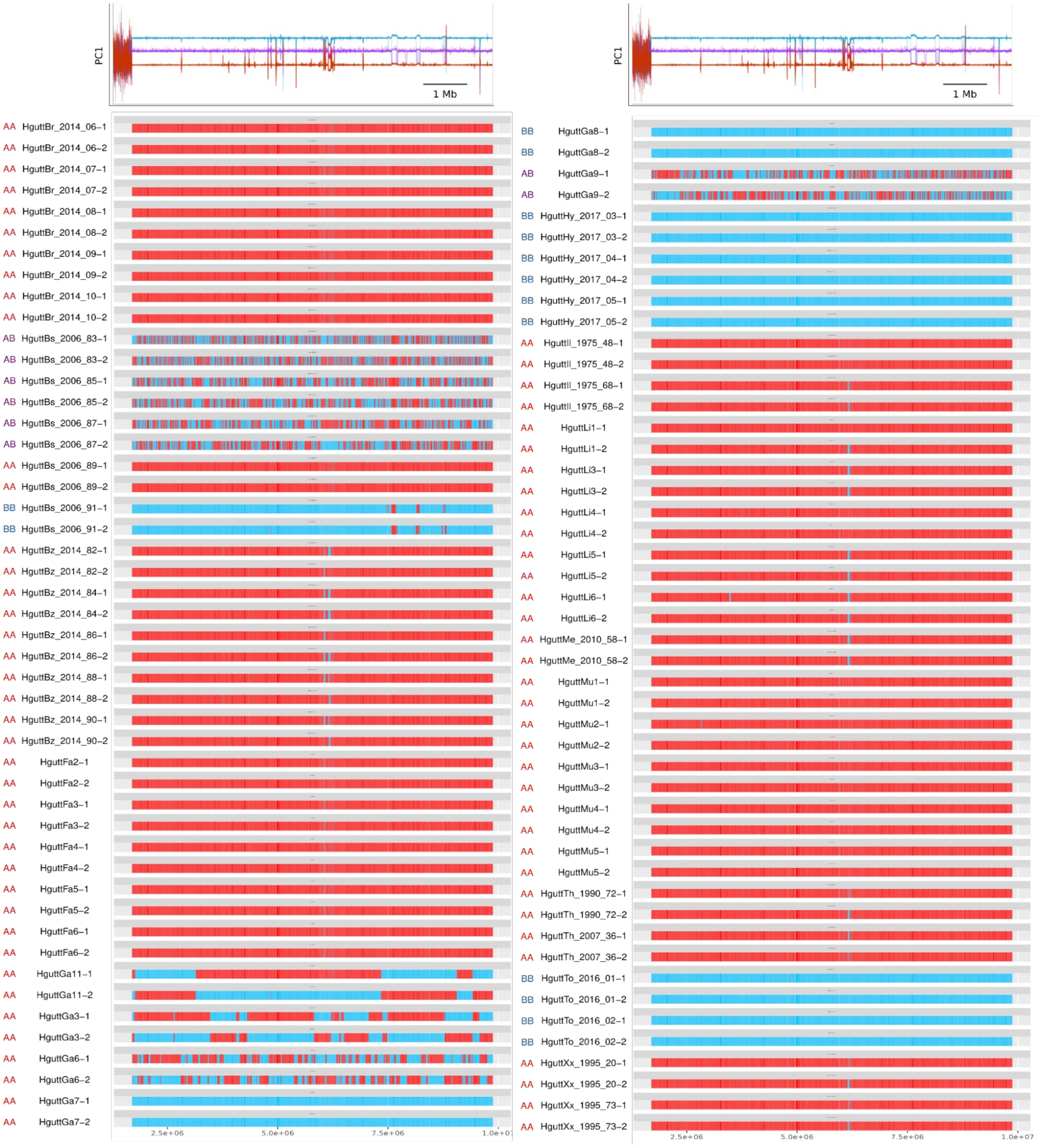

**Supplementary Figure S15a:** Chromosome painting in the inversion region on Chr2. Grey areas represent blocks where it was not possible to determine ancestry.

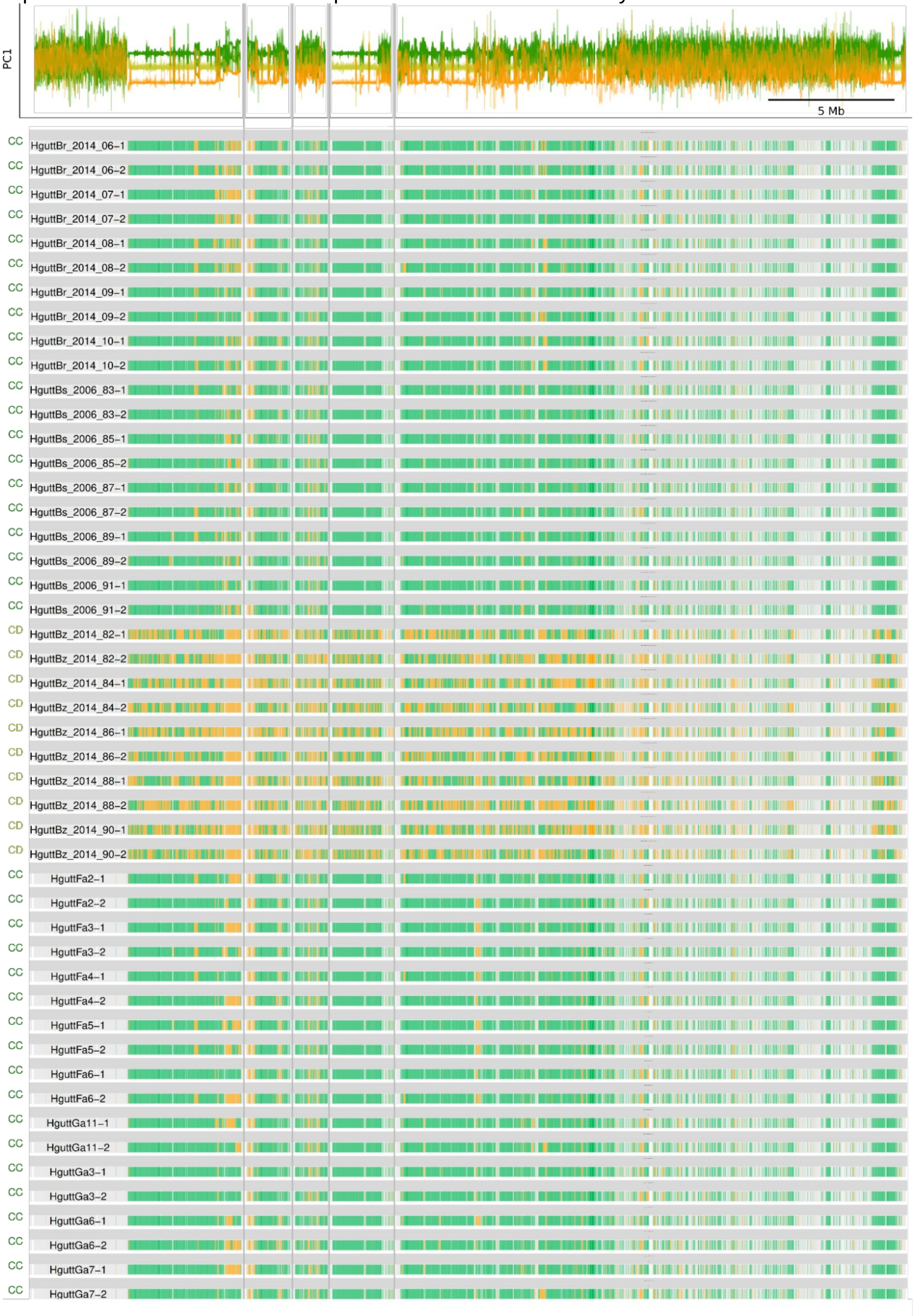

**Supplementary Figure S15b:** Chromosome painting in the inversion region on Chr2 (continued).

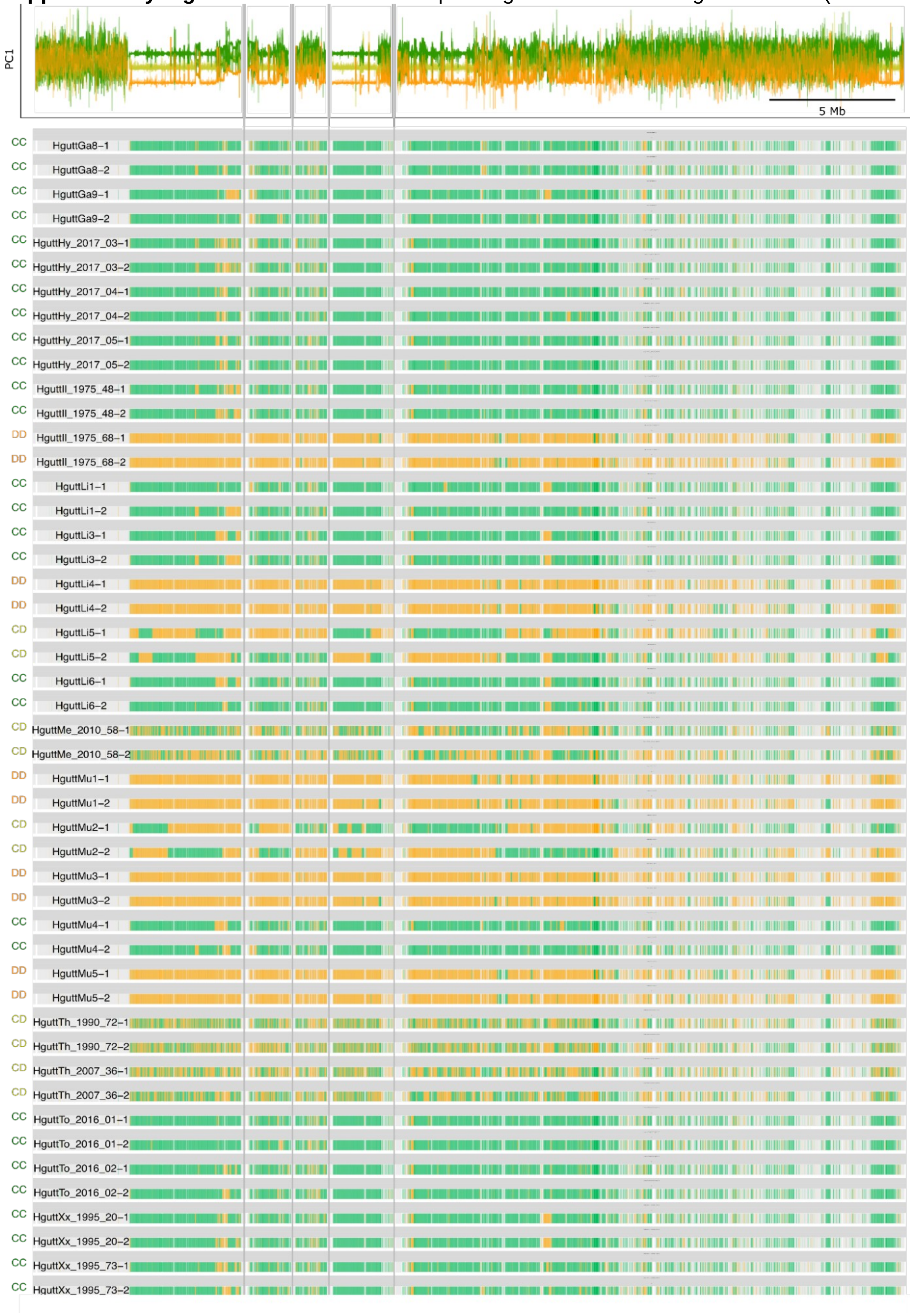

**Supplementary Figure S16:** Outlier SNPs identified in Riquet et al. (2019) mapped onto scaffold 14 (*H. erectus* Chr12) of our reference genome (Hgutt_V1). Allele frequencies calculated from our whole-genome data (n=112) are shown per sampling location. Grey numbers indicate sample sizes.

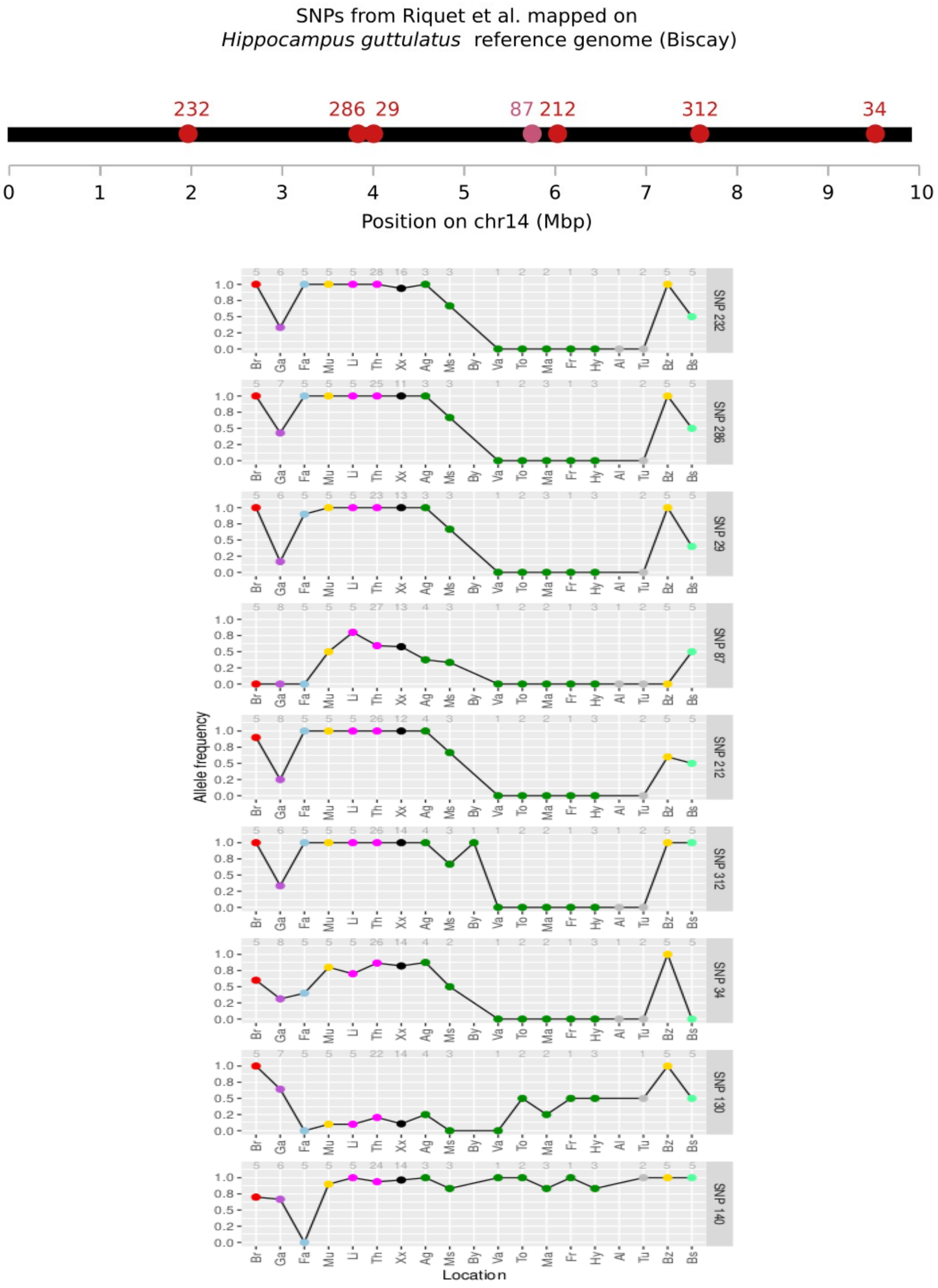

### Supplementary File S2

#### Scripts and commands:

**Reference genome assembly and repeat annotation**

| Step | Command |
|---|---|
| nubeam-dedup | #Deduplication of 10x genomics reads<br>nubeam-dedup -i1 Hgutt_S1_L001_R1_001.fastq.gz -i2 Hgutt_S1_L001_R2_001.fastq.gz -o1 Hgutt_DEDUP_S1_L001_R1_001.fastq.gz -o2 Hgutt_DEDUP_S1_L001_R2_001.fastq.gz -z 6 -r 0 |
| process_10xReads | #Process raw 10x genomics reads<br>process_10xReads.py -a -o Hgutt_DEDUP_PROC_S1 -1 Hgutt_DEDUP_S1_L001_R1_001.fastq.gz -2 Hgutt_DEDUP_S1_L001_R2_001.fastq.gz |
| custom R script | #Identification of rare and over-represented barcodes<br>Rscript --vanilla Autoset_Filters.R Hgutt_DEDUP_PROC_S1_barcodes.txt barocode_list.txt |
| filter_10XReads | #Filters reads for status and barcodes<br>filter_10xReads.py -m 11 -n 720 -o Hgutt_DEDUP_FILTERED_S1 -B Hgutt_DEDUP_PROC_S1_barcodes.txt -1 Hgutt_DEDUP_PROC_S1_R1_001.fastq.gz -2 Hgutt_DEDUP_PROC_S1_R2_001.fastq.gz |
| regen_10XReads | #Return reads to origin format<br>regen_10xReads.py -o Hgutt_DEDUP_REGEN_S1 -1 Hgutt_DEDUP_FILTERED_S1_R1_001.fastq.gz -2 Hgutt_DEDUP_FILTERED_S1_R2_001.fastq.gz |
| Supernova-2.1.1 | #Generate whole genome *de novo* assembly<br>supernova run --id Hgutt_DEDUP_FILTERED_S1_all --description DEDUP_FILTERED_S1_all_reads --fastqs /DEDUP_FILTERED --maxreads=all --accept-extreme-coverage<br><br>#Generate FASTA output for the assembly<br>supernova mkoutput --style=pseudohap2 --asmdir=Hgutt_DEDUP_FILTERED_S1_all/outs/assembly --outprefix=Hippocampus_guttulatus_v1 --minsize=1000 --index -- |

| | |
|---|---|
| | headers=short |
| BlobToolKit (v3.5.2)<br>BUSCO (v5.4.4) | #Create a new BlobDir dataset<br>blobtools create --fasta Hippocampus_guttulatus_v1.fasta --meta Hippocampus_guttulatus_v1.yaml /DATASETS/Hippocampus_guttulatus_v1<br><br>#Run BUSCO analysis<br>busco -i ./Hippocampus_guttulatus_v1.fasta -l actinopterygii -o Hgutt_v1_BUSCO -m geno<br><br>#Add results of BUSCO analysis to the BlobToolKit dataset<br>blobtools add --busco /Hgutt_v1_BUSCO/run_actinopterygii_odb10/full_table.tsv /DATASETS/Hippocampus_guttulatus_v1<br><br>#View results<br>blobtools view --local /DATASETS/Hippocampus_guttulatus_v1 |
| RepeatModeler2 | #Create database for RepeatModeler<br>BuildDatabase -name Hippocampus_guttulatus_v1 Hippocampus_guttulatus_v1.fasta<br><br>#Run RepeatModeler<br>nohup RepeatModeler -database Hippocampus_guttulatus_v1 -threads 20 -LTRStruct >& run.out & |
| Tandem Repeats Finder (v4.09.1) | #Run pyTanFinder<br>pyTanFinder.py Hippocampus_guttulatus_v1.fasta -minM 50 -maxM 2000 -minMN 2 -minA 100000 -px HguttV1_Tan -tp ./trf409.linux64 |
| RepeatMasker (v4.0.5) | #Run RepeatMasker<br>RepeatMasker -lib Hippocampus_guttulatus_v1-families.fa -pa 12 -a Hippocampus_guttulatus_v1.fasta<br><br>#Calculate divergence for each repeat family<br>calcDivergenceFromAlign.pl -s Hippocampus_guttulatus_v1.divsum Hippocampus_guttulatus_v1.fasta.align<br><br>#Generate Repeat Landscape<br>createRepeatLandscape.pl -div ./Hippocampus_guttulatus_v1.divsum -g |

| | |
|---|---|
| | 424000000 > ./Hgutt_Repeat_Landscape.html |

**Whole-genome sequencing data**

Some of the following commands were ran using snakemake (v7.1.1), for which snakefiles can be found at https://github.com/pierrebarry/life_tables_genetic_diversity_marine_fishes.

| Step | Command |
|---|---|
| fastp (v0.23.1) | fastp -i {input.raw_R1} -I {input.raw_R2} -o {output.fastp_R1} -O {output.fastp_R2} --trim_poly_g --correction --low_complexity_filter –html {output.report_html} --json {output.report_json} –report_title {wildcards.sample} --thread $ --dont_overwrite --merge –merged_out {output.merged} |
| bwa mem (v0.7.17) | #paired reads<br>bwa mem -M -t $ referencegenome_{wildcards.species} {input.fastq_R1} {input.fastq_R2} > {output.align_sam_paired})<br><br>#merged reads<br>bwa mem -M -t $ referencegenome_{wildcards.species} {input.fastq_merged} > {output.align_sam_merged})<br><br>#combine sam files<br>picard MergeSamFiles I={input.align_sam_paired} I={input.align_sam_merged} O={output.sam_final} |
| Sam to bam file (picard v2.26.8) | picard SortSam -I {input.align_sam} -O {output.align_bam_picard} -SO coordinate -CREATE_INDEX true -VALIDATION_STRINGENCY LENIENT -TMP_DIR tmp |
| Mark duplicates (picard v2.26.8) | picard -Xmx$g MarkDuplicates -I {input.align_bam_picard} -O {output.markdup_picard} -ASSUME_SORTED TRUE -REMOVE_DUPLICATES FALSE -CREATE_INDEX TRUE -METRICS_FILE {wildcards.sample}_duplicate_metrics.txt -VALIDATION_STRINGENCY LENIENT -TMP_DIR tmp |
| Add read group (picard v2.26.8) | picard AddOrReplaceReadGroups -I {input.markdup_picard} -O {output.markdup_rg_picard} -RGPL ILLUMINA -RGLB lib -RGPU genewiz -RGSM {wildcards.sample} |
| Get stats (samtools v1.10, htslib v1.10.2) | samtools flagstat {input.bamfile} > {output.samtools_flagstat}<br><br>samtools stats -d {input.bamfile} > {output.samtools_stats} |

| | |
|---|---|
| pmdtools<br>(v0.50) | for i in *.bam<br>do<br>echo pmdtools running on: $i<br>samtools view $i \| python /path/pmdtools --deamination > $i_pmdtools.txt<br>done<br><br>#Plotted with their Rscript plotPMD.R |
| GATK HaplotypeCaller (GATK 4.1.8.0) | gatk HaplotypeCaller -R {input.reference_genome} -I {input.markdup_rg_picard} -O {output.gvcf_first} -bamout {output.realign_bam} -ERC GVCF -G StandardAnnotation -G AS_StandardAnnotation -G StandardHCAnnotation --tmp-dir tmp |
| GATK GenomicsDBImport | gatk --java-options '-Xms$g -Xmx$G' GenomicsDBImport<br>--sample-name-map name_Hgutt.txt –genomicsdb-workspace-path {output.joint_genotyping_files} --tmp-dir /home/lmeyer/tmp<br><br>#name_Hgutt.txt contained sample names and the list of *_gvcf_first.g.vcf files |
| GATK GenotypeGVCFs | gatk --java-options '-Xms$g -Xmx$G' GenotypeGVCFs -R {input.reference_genome} -V gendb://Joint_Genotyping_{wildcards.interval} -G StandardAnnotation -O {output.joint_gvcf_first} |
| Merge VCFs (vcftools v0.1.16) | vcf-concat -f /path/list_vcf_files.txt \| bgzip -c > {output.vcf_first} |
| Vcf filtering (bcftools v1.9, vcftools v0.1.16) | #Filter VCF around indels<br>bcftools filter -g 5 --output {output.vcf_indel5bp} {input.vcf}<br><br>#Keep biallelic snps<br>vcftools --gzvcf {input.vcf_indel5bp} --remove-indels --max-alleles 2 --recode --stdout \| bgzip > {output.vcf_indel5bp_snponly}<br><br>#Filter missing data<br>vcftools --gzvcf {input.vcf_indel5bp_snponly} –max-missing {params.per} --recode --stdout \| bgzip > {output.vcf_indel5bp_snponly_missing}<br><br>#Remove sites with extremely high depth<br>vcftools --exclude-positions {list} –vcfgz {output.vcf_indel5bp_snponly_missing} --recode --recode-INFO-all --out {output.vcf_indel5bp_snponly_missing_depth} |
| ANGSD<br>(v0.933) | #ANGSD all SNPs<br>angsd -bam {bam_file_list} -GL 2 -trim 5 -setMaxDepth 2300 -doMajorMinor 1 -minMapQ 30 -minQ 20 -doMaf 1 -SNP_pval 1e-6 -minInd 40 -doCounts 1 -minMaf 0.05 -doGlf 2 -uniqueonly 1 -remove_bads 1 -C 50 -baq 1 -doCov 1 -doIBS 2 -makeMatrix 1 -ref referencegenome_Hgutt_V1.fa -out {prefix} -P $ 2>log.txt |
| ANGSD<br>(v0.933) | #ANGSD markers in linkage equilibrium<br>angsd -bam {bam_file_list} -sites {list_sites} -rf chrs.txt -GL 2 -trim 5 -setMaxDepth 2300 -doMajorMinor 1 -minMapQ 30 -minQ 20 -doMaf 1 -minInd 40 -doCounts 1 -minMaf 0.05 -doGlf 2 -uniqueonly 1 -remove_bads 1 -C 50 -baq 1 -doCov 1 -doIBS 2 -makeMatrix 1 -ref referencegenome_Hgutt_V1.fa -out {prefix} -P $ 2>log.txt |

| | |
|---|---|
| SNPRelate v1.28.0, SeqVartools v1.38.0, R v4.3.0) | showfile.gds(closeall=TRUE)<br>VCF_PATH="/path/file.vcf.gz"<br><br> if (file.exists("file.gds")==F){<br> vcf.fn <- VCF_PATH<br> seqVCF2GDS(vcf.fn, "file.gds")<br> }<br><br>#OPEN GDS (can start from here)<br>genofile <- seqOpen("/path/file.gds")<br><br>#Perform PCA<br>pca <- snpgdsPCA(genofile, num.thread=5,autosome.only = T, maf=0.05) |
| lostruct (v0.0.0.9) | snps<-vcf_windower("/path/file.bcf",size=5000,type='bp')<br><br>pcs <- eigen_windows(snps,k=2)<br><br>write.table(pcs,"filename.txt") |
| genomics_general (python v3.9.13) | #Parse VCF and generate .geno format<br>python /path/parseVCF.py -i file.vcf --skipIndels --minQual 30 --gtf flag=DP min=2 max=100 -o file.geno.gz<br><br># popgenWindows.py<br>#diversity and divergence along the genome<br>#Example for contrast between pops Mu and Ma<br>python /path/popgenWindows.py --windType coordinate -w 25000 -m 100 -g file.geno.gz -o file.csv.gz -f phased -T $ -p Mu HguttMu1,HguttMu2,HguttMu3,HguttMu4,HguttMu5 -p Ma Hgutt_Hy_2017_03,Hgutt_Hy_2017_04,Hgutt_Hy_2017_05,Hgutt_To_2016_01,Hgutt_To_2016_02<br><br>#Scripts available at https://github.com/simonhmartin/genomics_general |
| Heterozygosity (vcftools v0.1.16) | vcftools --gzvcf file.vcf.gz --het --out prefix |
| BAMscorer (v1.4) | #select SNPs<br><br>BAMscorer select_snps file.vcf.gz output_prefix --numchrom $<br><br>#Manual step to get 3 files with the individuals of each karyotype<br>#{OUT}_AA_individuals.txt<br>#{OUT}_BB_individuals.txt<br>#{OUT}_db_individuals.txt<br><br>#score bam files<br>BAMscorer score_bams file.vcf.gz output_prefix path_to_bams |

<table>
<tr><td>Twisst</td><td>#Produce a VCF with ANGSD including other spp.<br>angsd -bam bamlist -r $ -ref referencegenome_Hgutt_V1.fa -GL 2 -doPost 1 -doGeno 1 -trim 5 -setMaxDepth 330 -doMajorMinor 1 -minMapQ 30 -minQ 20 -doMaf 1 -minInd 8 -doCounts 1 -doGlf 2 -uniqueonly 1 -remove_bads 1 -C 50 -baq 1 -doIBS 2 -doCov 1 -makeMatrix 1 -doBcf 1 --ignore-RG 0 -out {prefix} -P $ 2>log.txt<br><br>#Remove lines with heterozygous genotypes<br>bcftools view file.bcf| grep -v "0/1" > file_noHet.vcf<br><br>#Remove lines where all species are "0/0"<br>bcftools view file_noHet.vcf| grep "1/1" > tmp.vcf<br>bcftools view -h file_noHet.vcf > VCF_header<br>cat VCF_header tmp.vcf > file_noHet_variable.vcf #also added the header<br>rm tmp.vcf<br><br>#Make it a "phased" VCF<br>bcftools view -H file_noHet_variable.vcf | sed 's/\//|/g' > tmp.vcf<br>cat VCF_header tmp.vcf > file_noHet_variable_phased.vcf #also added the header<br>rm tmp.vcf VCF_header<br>bgzip file_noHet_variable_phased.vcf<br><br>#Convert to .geno format<br>python /path/parseVCF.py -i file_noHet_variable_phased.vcf.gz --skipIndels --gtf flag=DP min=2 max=100 | gzip > file_noHet_variable_phased.geno.gz<br><br>#Infer the trees<br>python path/phyml_sliding_windows.py --minPerInd 25 -T $ -g file_noHet_variable_phased.geno.gz --prefix prefix.phyml_bionj.w50 -w 50 --windType sites --model GTR --optimise n<br><br>#Get topology weights<br>python /path/twisst.py -t prefix.phyml_bionj.w50.trees.gz prefix.phyml_bionj.w50.weights.csv.gz -g A -g B -g C -g D -g E -g F -g G -g H --groupsFile groups.tsv<br><br>#Scripts can be found at https://github.com/simonhmartin/twisst</td></tr>
<tr><td>SHAPEIT (v4.2.2)</td><td>#without gmap<br>for i in $(cat scaffold_list);<br>do shapeit4 \<br>--input file.bcf \<br>--region ${i} \<br>--effective-size X \<br>--output prefix_phased_${i}.vcf.gz \<br>--log phasing.log;<br>done</td></tr>
<tr><td>BLAST</td><td>XXX</td></tr>
<tr><td>tsinfer tsdate</td><td>#create samples using tsinfer_create_input_files.py<br><br>#run tsinfer using tsinfer_infer.py<br><br>#run tsdate using tsdate_infer_part1.py</td></tr>
</table>

<table>
<tr><td></td><td>#extract TMRCA using tsinfer_extract_info_v2.py</td></tr>
</table>

**Analysis of SVs**

<table>
<tr><td>Step</td><td>Command</td></tr>
<tr><td>EMA (v0.6.2)</td><td>#Interleave read files<br>parallel -j20 --bar 'paste <(pigz -c -d {} | paste - - - -) <(pigz -c -d {= s:_R1_:_R2_: =} | paste - - - -) | tr "\t" "\n" | ema count -w ./barcode_list.txt -o {/.} 2>{/.}.log' ::: *_R1_*.gz<br><br><br>#Preprocess 10x data and insert BX:Z tags<br>paste <(pigz -c -d *_R1_*.gz | paste - - - -) <(pigz -c -d *_R2_*.gz | paste - - - -) | tr "\t" "\n" | ema preproc -w ./barcode_list.txt -b -n 500 -t 20 -o output_dir *.ema-ncnt 2>&1 | tee preproc.log<br><br><br>#Concatenate files<br>cat ema-bin-* > Hgutt_DEDUP_REGEN_S1_Interleaved_BXed.fastq</td></tr>
<tr><td>LRez (v2.2.4)</td><td>#Build barcode index<br>LRez index fastq -f Hgutt_DEDUP_REGEN_S1_Interleaved_BXed.fastq -o Barcode_Index.bci -t 20</td></tr>
<tr><td>BWA (v0.7.17)</td><td>#Map linked-reads to the reference genomes Hgutt_V1 and HguttRefA<br>bwa mem -C -t 20 Hippocampus_guttulatus_v1.fasta -p Hgutt_DEDUP_REGEN_S1_Interleaved_BXed.fastq.gz > Hgutt_10XLR_Aligned_V1.sam<br>bwa mem -C -t 20 GCA_025802095.1_ASM2580209v1_genomic.fna -p Hgutt_DEDUP_REGEN_S1_Interleaved_BXed.fastq.gz > Hgutt_10XLR_Aligned_RefA.sam<br><br><br>#Sort and convert to bam and make index<br>samtools sort Hgutt_10XLR_Aligned_V1.sam -@ 4 -O bam -l 0 -m 2G -o Hgutt_10XLR_Aligned_V1.bam<br>samtools index -b Hgutt_10XLR_Aligned_V1.bam<br>samtools sort Hgutt_10XLR_Aligned_RefA.sam -@ 4 -O bam -l 0 -m 2G -o Hgutt_10XLR_Aligned_RefA.bam<br>samtools index -b Hgutt_10XLR_Aligned_RefA.bam</td></tr>
<tr><td>MTG-Link</td><td>#Generate the input GFA file</td></tr>
</table>

| | |
|---|---|
| | bed2gfa.py -bed Hgutt_14_1698000_1700000.bed -fa Hippocampus_guttulatus_v1.fasta -out Hgutt_14_1698000_1700000.gfa<br><br>#Run MTG-Link<br>mtglink.py DBG -gfa Hgutt_14_1698000_1700000.gfa -bam Hgutt_10XLR_Aligned_V1.bam -fastq Hgutt_DEDUP_REGEN_S1_Interleaved_BXed.fastq -index Barcode_Index.bci -k 61 51 41 31 21 -t 20 |
| Minimap2 (v2.26) | # Align assembled contig on the reference genome<br>minimap2 -a Hippocampus_guttulatus_v1.fasta Hgutt_14_1698000_1700000.gfa.14_0-1698000-L+_14_1700000-9878394-R+.g2000.flank10000.occ2.k61.a3.bxu.insertions_filtered_quality.fasta > 3kb_contig.sam |
| Leviathan (v1.0.2) | #Extract data mapping to Chr4.1 of HguttRefA<br>samtools view -b Hgutt_10XLR_Aligned_RefA.bam JAOYMQ010000004.1 > Hgutt_10XLR_Aligned_RefA_4.1.bam<br>samtools index Hgutt_10XLR_Aligned_RefA_4.1.bam<br><br>#Build LRez barcode index<br>LRez index bam -p -b Hgutt_10XLR_Aligned_RefA_4.1.bam -o Hgutt_10XLR_Aligned_RefA_4.1.bci<br><br>#Run Leviathan on Chromosome 4.1<br>LEVIATHAN -b Hgutt_10XLR_Aligned_RefA_4.1.bam -i Hgutt_10XLR_Aligned_RefA_4.1.bci -g GCA_025802095.1_ASM2580209v1_genomic.fna -o Hgutt_10XLR_Aligned_RefA_4.1.vcf |
